# Horse gluteus is a null-sarcolipin muscle with enhanced sarcoplasmic reticulum calcium transport

**DOI:** 10.1101/688531

**Authors:** Joseph M. Autry, Christine B. Karim, Bengt Svensson, Samuel F. Carlson, Mariana Cocco, Sudeep Perumbakkam, Zhenhui Chen, L. Michel Espinoza-Fonseca, Carrie J. Finno, David D. Thomas, Stephanie J. Valberg

## Abstract

We have analyzed gene transcription, protein expression, and enzymatic activity of the Ca^2+^-transporting ATPase (SERCA) in horse gluteal muscle. Horses are bred for peak athletic performance but exhibit a high incidence of exertional rhabdomyolysis, with myosolic Ca^2+^ suggested as a correlative linkage. To assess Ca^2+^ regulation in horse gluteus, we developed an improved protocol for isolating horse sarcoplasmic reticulum (SR) vesicles. RNA-seq and immunoblotting determined that the *ATP2A1* gene (protein product SERCA1) is the predominant Ca^2+^-ATPase expressed in horse gluteus, as in rabbit muscle. Gene expression was assessed for four regulatory peptides of SERCA, finding that sarcolipin (*SLN*) is the predominant regulatory peptide transcript expressed in horse gluteus, as in rabbit muscle. Surprisingly, the RNA transcription ratio of *SLN*-to-*ATP2A1* in horse gluteus is an order of magnitude higher than in rabbit muscle, but conversely, the protein expression ratio of SLN-to-SERCA1 in horse gluteus is an order of magnitude lower than in rabbit. Thus, the *SLN* gene is not translated to a stable protein in horse gluteus, yet the supra-high level of *SLN* RNA suggests a non-coding role. Gel-stain analysis revealed that horse SR expresses calsequestrin (CASQ) protein abundantly, with a CASQ-to-SERCA ratio ∼3-fold greater than rabbit SR. The Ca^2+^ transport rate of horse SR vesicles is ∼2-fold greater than rabbit SR, suggesting horse myocytes have enhanced luminal Ca^2+^ stores that increase intracellular Ca^2+^ release and muscular performance. The absence of SLN inhibition of SERCA and the abundant expression of CASQ may potentiate horse muscle contractility and susceptibility to exertional rhabdomyolysis.

## Introduction

The horse has been selectively bred for thousands of years to achieve remarkable athletic ability, in part conferred by a naturally-high proportion (75–95%) of fast-twitch myofibers in locomotor muscles that provide powerful contractions and high speed (1). Calcium (Ca^2+^)^1^ is the signaling molecule that controls muscle contraction by activating the actomyosin sliding-filament mechanism (2). In turn, myosolic Ca^2+^ is controlled predominantly by the sarcoplasmic reticulum (SR), a membrane organelle that connects with transverse tubules as membrane junctions and also surrounds actomyosin myofibrils as longitudinal sheaths (3). Thus, SR is a dual structure/function membrane system comprising junctional SR with the ryanodine receptor Ca^2+^ release channel (RYR) acting to initiate contraction, and longitudinal SR with the Ca^2+^ transporting ATPase (SERCA) acting to induce relaxation (4, 5). For muscle contraction, the bolus of released Ca^2+^ originates almost solely from SR through the RYR channel, and this released Ca^2+^ is subsequently re-sequestered in the SR lumen by SERCA (6). The strength of muscle contraction is controlled by the amount of Ca^2+^ released from SR, which is in turn modulated by the Ca^2+^ load in the SR lumen (7). Calsequestrin (CASQ), which binds up to 80 Ca^2+^ ions (mol/mol), is the high-capacity Ca^2+^ storage protein localized to junctional SR (8, 9). Per its luminal Ca^2+^ load, CASQ regulates RYR opening and closing (10, 11).

SERCA is a prototypical member of the P-type ion-motive transport ATPase family: central to the long-term understanding of active membrane transport (12, 13). Importantly, the Ca^2+^ transport activity of SERCA is regulated by a family of single-pass transmembrane peptides in SR, including sarcolipin (SLN) and phospholamban (PLN). SLN and PLN were discovered in the early 1970s (14, 15) and have since been characterized as potent regulators of contractility, dependent upon post-translational modifications and protein expression level (16–20). In the past few years, five additional peptide regulators of SERCA have been reported in mouse: the inhibitor myoregulin (MRLN) in muscle, the inhibitor small ankyrin 1 (sAnk1) in muscle, the activator dwarf open reading frame (DWORF) in ventricles, the inhibitor endoregulin (ELN) in endothelial and epithelial cells, and the inhibitor ‘another-regulin’ (ALN) in non-muscle tissues (21–25).

In larger mammals such as rabbit, dog, and pig, SLN is the primary regulatory peptide expressed in fast-twitch skeletal muscle (18,26–28). SLN reportedly inhibits SERCA activity through multiple enzymatic mechanisms: by decreasing maximal velocity (V_max_), by decreasing Ca^2+^ binding affinity (1/K_Ca_), and by decreasing the number of Ca^2+^ ions transported per ATP molecule hydrolyzed (coupling stoichiometry) below the optimal Ca^2+^/ATP coupling ratio of two (29–32). SLN inhibition is relieved in part by phosphorylation and de-acylation (17,33,34). In mouse slow-twitch muscle, genetic knock-out of SLN expression results in enhanced ATP-dependent Ca^2+^ uptake by SERCA (35). In patients with atrial fibrillation or heart failure with preserved ejection fraction, decreased expression of SLN correlates with increased Ca^2+^ uptake in atrial homogenates; however, it is unknown if decreased SLN inhibition and concomitant SERCA activation are compensatory or causative in disease progression (36, 37). Aberrant Ca^2+^ signaling from SR Ca^2+^ stores in skeletal myocytes has been detected in malignant hyperthermia due to RYR mutation and an experimental model of increased sarcolemmal Ca^2+^ entry that leads to store overload-induced Ca^2+^ release (SOICR) through RYR (38, 39). Thus, SERCA activity and its regulation must be tuned to match the functional requirements of Ca^2+^ transport kinetics for contraction and relaxation of each specific muscle type.

Valberg *et al*. reported recently the gene expression level and the deduced protein sequence of *SLN*, *MRLN*, and *DWORF* in skeletal muscle of three horse breeds: Thoroughbred, Standardbred, and Quarter Horse (40). Similar to other large mammals, the *SLN* gene in horse gluteus is the highest expressed RNA transcript in the family of SERCA regulatory peptides (40). The horse SLN peptide encodes 29 residues (40) versus a consensus 31-residue length for orthologous SLN peptides encoded by 100+ species from the five vertebrate classes: mammal, bird, reptile, amphibian, and fish (18,41,42). Uniquely, horse SLN is the only reported ortholog missing four regulatory sites (18,40–42) that are common in other species: Ser4 and Thr5 for SLN phosphorylation and relief of SERCA inhibition (17, 33), Cys9 for SLN acylation and relief of SERCA uncoupling (18, 34), and Tyr31 for SERCA interaction, organelle targeting of SLN, and potential luminal nitration of SLN (31,43,44), with residues numbered using the consensus 31-residue length, e.g., rabbit, mouse, and human (18,40–42).

In comparison, horse MRLN is a 46-residue transmembrane peptide, similar to human, rabbit, and mouse orthologs, with 5–7 potential phosphorylation sites (Ser or Thr) in the N-terminus, as deduced per amino acid sequences (40). Before the *MRLN* gene product was identified as a sORF coding for a translated MLRN peptide that inhibits SERCA in mouse muscle, the *MRLN* transcript was identified as a functional long non-noncoding RNA (lncRNA) that controls myogenic differentiation, classified as *Linc-RAM* and *LINC00948* in mouse and human genomes, respectively (21,45,46). Similarly, the DWORF peptide was discovered as a small open reading frame (sORF) using bioinformatic analysis of the mouse transcriptome, thereby re-classifying the previously-assigned *DWORF* lncRNA transcript (*NONMMUG026737*) as a dual lnc/mRNA that also encodes a sORF translated into DWORF, a 35-residue transmembrane peptide that activates SERCA in mouse heart (22, 24).

The role of SERCA regulatory peptides in horse SR function is unknown. Microsomal membranes enriched in SR vesicles have been used traditionally since the 1960s to study SERCA Ca^2+^ transport and ATPase activities *in vitro*. These SR vesicles are purified from muscle homogenates using differential centrifugation, yielding sealed right-side-out SR vesicles, albeit with varying degrees of purity. Rabbit fast-twitch muscle is the standard model; early preparations of rabbit muscle SR vesicles were contaminated by significant amounts of glycogen granules, nucleic acids, and cellular debris (“glycogen-sarcovesicular fraction”) (47), yet further developments yielded highly-purified SR vesicles that have been well-characterized and widely-used (47–50). Protocols for preparation of SR vesicles have also been developed for skeletal muscles of mouse, rat, squirrel, chicken, dog, pig, and cow, yet SR vesicles from rabbit muscle has remained the field standard for use in biochemistry, biophysics, and crystallography (13,51,52). Purification of SR vesicles from horse muscles has been achieved, and these SR preparations have been utilized to assess RYR and SERCA activity (53–57). However, SR vesicles purified from horse muscle show low SERCA activity compared to SR vesicles from rabbit muscle, which has seemed inconsistent with the high fast-twitch fiber muscular composition of horse breeds.

Horses are highly susceptible to exertional rhabdomyolysis from a variety of causes including glycogen storage disorders, malignant hyperthermia, and other abnormalities in myosolic Ca^2+^ regulation (58). Recurrent exertional rhabdomyolysis (RER) is one of the most common cause of poor performance and economic loss in racehorses (59, 60). The molecular etiology of RER in Thoroughbred racehorses has been suggested to involve SR Ca^2+^ release and excitation-contraction coupling (57,58,61,62). We determined recently that gene transcription of *CASQ1* (skeletal muscle isoform) is downregulated in the gluteal muscle of male or female horses with RER compared to healthy male or females horses (controls), while CASQ1 was also downregulated to a greater extent in the gluteal muscle of male horses with RER compared to female horses with RER (40).

Horse encodes unique Ca^2+^ regulatory genes that likely have been selected for athletic performance and thus could impact rhabdomyolysis, so the goal of the present study was to examine Ca^2+^ transport regulation in horse SR vesicles. We hypothesized that horse, through natural selection as prey animals and subsequent selective breeding, has highly-adapted mechanisms for Ca^2+^ transport regulation that contribute to powerful muscular performance. This hypothesis builds upon phylogenomic analyses of SLN across the 131 vertebrate species SLN dataset of Gaudry *et al*. that identified novel deletions and replacements in the equid peptide that are predicted to control its regulation and function (41, 42).

Here we add multi-disciplinary approaches (transcriptomic, computational, and biochemical) to investigate Ca^2+^ transport mechanisms specific to horse gluteus medius, a predominant locomotor muscle. Enzymatic properties of horse SERCA were compared to SERCA from a laboratory-bred rabbit breed that is a standard model of skeletal muscle SR function. Our research identified unique modes of *SLN* transcription/translation, SERCA Ca^2+^ transport activity, and robust CASQ expression in healthy horse muscle, and these findings were interpreted in light of the high susceptibility of horses to exertional rhabdomyolysis. We propose that comparative studies of gene expression and biochemical regulation of SR enzymes will increase the broader understanding of selective adaptation of horse and human muscle, with a specific focus towards performance and disease.

## Results

### Transcription of *ATP2A* genes (encoding SERCA proteins) in horse gluteal muscle

SERCA Ca^2+^-transporting ATPases are encoded by a family of three genes (*ATP2A1*, *ATP2A2*, and *ATP2A3*) that encode protein isoforms SERCA1 (typically expressed in fast-twitch myocytes), SERCA2 (slow-twitch myocytes, cardiomyocytes, and non-muscle cells), and SERCA3 (smooth myocytes, platelet cells, and non-muscle cells), respectively. RNA transcriptome shotgun sequencing (RNA-seq) of the middle gluteal muscle of Arabian horses (63) was used to assess expression of the *ATP2A* gene family, per quantification in transcripts per million reads (TPM) reported as mean ± standard deviation (SD) (**Figure 1**), including comparison to muscle myocyte composition. Histological analysis determined a mean ± SD of 4.9 ± 0.9 fold greater abundance of fast-twitch myocytes (type 2A = 52 ± 8% plus type 2X = 31 ± 8%) versus slow-twitch myocytes (type 1 = 17 ± 2%) in the middle gluteal muscle of six healthy Arabian horses (N = 6) (64). Here RNA-seq analysis of middle gluteal muscle from the same six Arabian horses determined that the *ATP2A1* gene encoding SERCA1 is expressed at 7.3 ± 2.8 fold greater level (mean ± SD) than the *ATP2A2* gene encoding SERCA2 (**Figure 1*A,C***). These novel RNA-seq results (**Figure 1**) are consistent with histological analysis of myocyte type (64), and it is well-documented that the SERCA1 protein is expressed at higher density than the SERCA2 protein in fast-twitch muscles from a wide range of non-equine mammalian species (26). The middle gluteal muscle showed insignificant expression of the horse *ATP2A3* gene, which is transcribed at a level comprising ∼ 0.04% of total transcripts from the *ATP2A* gene family (TPM/TPM) (**Figure 1*A***). RNA-seq data generated from gluteal muscle of the set of Arabian horses were deposited in the National Center for Biotechnology Information (NCBI) Sequence Read Archive (SRA) database (65) with accession number <u>SRP082284</u>^2^.

**Figure 1.**
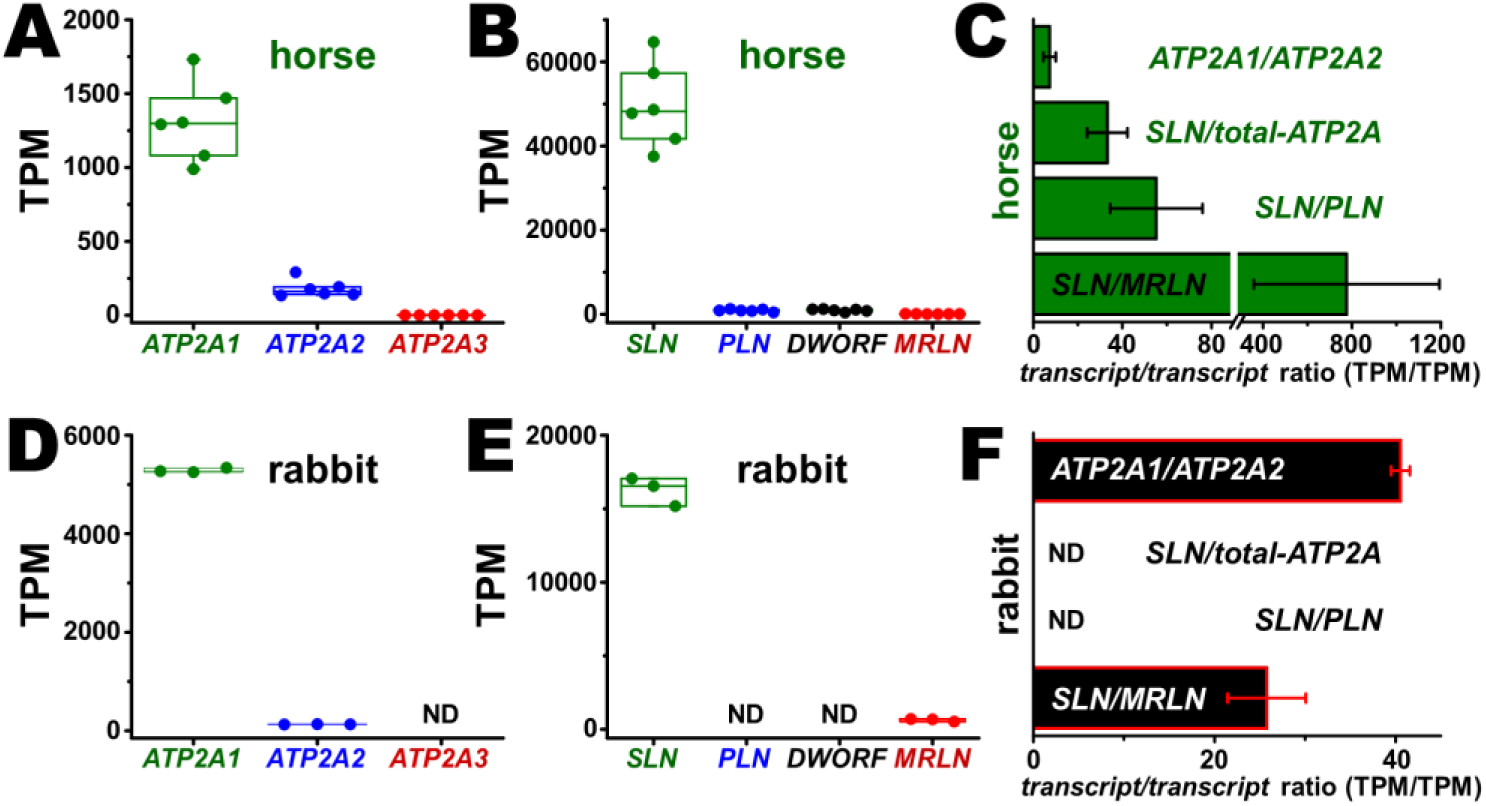
RNA-seq quantitation of gene transcription for calcium regulatory proteins in horse muscle. *A*, *B*, *C*, *D*, the number of target transcripts per million reads (TPM) are reported, using a box plot to indicate the mean, maximum, minimum, and range (N = 6 Arabian horses and N = 3 *O. cuniculus* rabbits). *A*, *D*, *ATP2A1* (SERCA1), *ATP2A2* (SERCA2), and *ATP2A3* (SERCA3) in horse gluteus and rabbit muscle, respectively. *B*, *E*, *SLN*, *PLN*, *DWORF*, and *MLRN* in horse gluteus and rabbit muscle, respectively. *C*, *F*, the relative ratio of *transcript*/*transcript* expression (TPM/TPM) in horse gluteus and rabbit muscle, respectively. The relative ratio of *transcript*/*transcript* expression (TPM/TPM) of *SLN* to the sum of the *ATP2A* gene family is also reported (*SLN/total-ATP2A*). The expression level of horse genes were mined from RNA-seq data reported by Valberg *et al*. (64) with SRA accession number SRP082284^2^. The expression level of rabbit genes were mined from RNA-seq data reported in SRA accession number SAMN00013649^2^. ND indicates ‘not determined’.

For comparison, histological analysis of seven types of rabbit back and leg muscles determined a combined predominance of fast-twitch myocytes (type 2A = 21 ± 7.6% plus type 2B = 76 ± 8.7%) versus slow-twitch myocytes (type I = 3.1 ± 3.1%) (66). Here, we mined RNA-seq data of *ATP2A1* and *ATP2A2* gene expression in rabbit muscle from SRA accession number SAMN00013649^2^ (rabbit species *Oryctolagus cuniculus*). A compilation of RNA-seq data from three laboratory rabbits (SAMN00013649^2^) demonstrated that the *ATP2A1* gene (SERCA1) is expressed at 41 ± 1.0 fold greater level than *ATP2A2* (SERCA2) (**Figure 1*D,F***). Thus, both horse gluteal muscle and rabbit back and leg muscles express predominantly the *ATP2A1* gene and are composed mostly of fast-twitch myocytes, thereby providing a valid platform for comparative transcriptomic and biochemical analyses of SR function.

### Transcription of *SLN*, *PLN*, *MRLN*, and *DWORF* genes (encoding SERCA regulatory peptides) in horse gluteal muscle

We previously used quantitative reverse-transcriptase polymerase chain reaction (qRT-PCR) to quantify transcription of four genes (*SLN*, *PLN*, *MRLN*, and *DWORF*) encoding SERCA regulatory peptides in horse gluteus muscle, finding that *SLN* is the predominant regulatory peptide gene expressed in the middle gluteal muscle of Thoroughbred horses (40). Here we mined RNA-seq data (64) to quantify transcription of the same four genes encoding SERCA regulatory peptides (*SLN*, *PLN*, *MRLN*, and *DWORF*) in the middle gluteal muscle of six Arabian horses. RNA-seq determined that *SLN* is the predominant regulatory peptide gene transcribed in the gluteal muscle of Arabian horses, with >55-fold greater expression than *PLN* and *DWORF* genes, and 780 ± 420 fold greater expression than *MRLN* (**Figure 1*B,C***). RNA-seq also determined that transcription of the *SLN* gene is 38 ± 11 fold greater than *ATP2A1* (SERCA1), thereby demonstrating that the *SLN* gene is highly expressed in the horse gluteus muscle (**Figure 1*A‒C***).

For comparison, we mined reported RNA-seq data for rabbit muscle reported in SRA accession number <u>SAMN00013649</u>^2^ to determine relative gene expression of SERCA regulatory peptides in horse versus rabbit muscle. A compilation of data from three laboratory rabbits indicated that the *SLN* gene is transcribed as the predominant regulatory peptide, with 26 ± 4.3 fold greater gene expression than *MRLN* (**Figure 1*C***). RNA-seq data further indicated that *PLN* gene expression was undetected in rabbit muscle (SRA <u>SAMN00013649</u>^2^). Thus, RNA-seq from horse and rabbit muscles provides results consistent with previous qRT-PCR studies which report that in non-rodent mammals such as rabbit, pig, and horse, *SLN* is the predominantly regulatory gene expressed in fast-twitch muscle (26, 40). On the other hand, *MRLN* is the primary regulatory peptide gene expressed in mouse muscles (21). When comparing the relative ratio of gene expression of *SLN* to the sum of the *ATP2A* genes (i.e., the SERCA1 and SERCA2 proteins), rabbit muscle has a transcription ratio of 3.1 ± 0.19 for *SLN*-to-*ATP2A1* (TPM/TPM), a ratio which is ∼12-fold lower than the *SLN*-to-*ATP2A1* ratio in horse muscle (**Figure 1**). Thus, horse gluteal muscle and rabbit fast-twitch muscle express *SLN* as the predominant regulatory peptide gene, verifying the utility of our experimental platform for comparative analyses of SR function.

### Amino acid sequence analysis and predicted isoelectric point of SLN orthologs

The horse *SLN* gene product encodes a 29-residue transmembrane peptide that is missing four canonical regulatory residues (S4, T5, C9, and Y31; numbered per the consensus amino acid sequence), as determined for Thoroughbred, Standardbred, Quarter Horse, and Przewalski’s Horse, plus zebra and donkey (40,41,64). Here RNA-seq was used to deduce the amino acid sequence of SLN of the Arabian horse (N = 6) (**Figure 2**), which encodes the identical 29-residue sequence found in other members of the genus *Equus* (40). The uniqueness of the amino acid sequence of horse SLN is emphasized by a sequence alignment of 131 SLN orthologs, where the consensus SLN is a 31-residue peptide encoding phosphorylation site(s) at positions 4 and/or 5, and a terminal Y31 residue (41) that is involved in SERCA inhibition and intracellular targeting (31). At position 9, the majority of SLN orthologs encode residue F9 (∼67%), while a large minority of orthologs (∼33%) encode residue C9 (41), which is a site for s-palmitoylation and s-oleoylation of rabbit and pig SLN (18). However, horse SLN lacks these regulatory residues, instead encoding R4, ΔT5, F8, and ΔY30, per horse residue numbering (**Figure 2**). For comparison, the most-closely related families to *Equus* are *Tapiridae* (tapirs) and *Rhinocerotidae* (rhinoceros), which both encode a 31-residue SLN with an amino acid sequence of high consensus: potential phosphorylation sites at T4 and T5, a C-terminal Y31 residue, yet F9, a residue that precludes acylation (18,40,41). Thus, the unique amino acid sequence of SLN shared by equine orthologs suggests novel molecular and structural mechanisms for regulation of SERCA activity in horse muscle (40).

**Figure 2.**
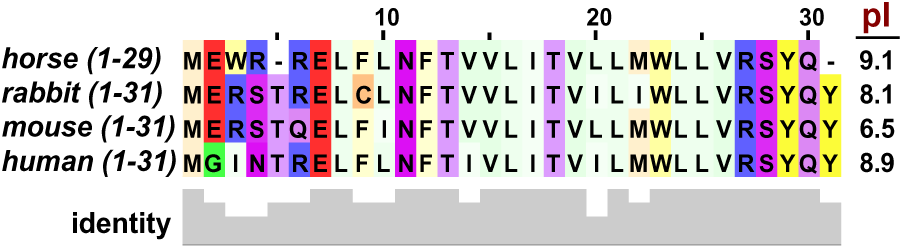
Amino acid sequence analysis and predicted isoelectric point of SLN orthologs. The amino acid sequence of horse SLN was deduced using genomic DNA sequencing and RNA-seq (40, 64). Residue numbering is indicated *above*, according to the consensus length of 31 amino acids, per rabbit, mouse, and human orthologs. The predicted isoelectric point (pI) of SLN orthologs is indicated on the *right*. Sequence identity at each residue position is indicated by the bar graph *below*. Amino acids are highlighted using a color scale that shows sulphur-containing residues in *orange* (M, C), negatively-charged residues in *red* (E), positively-charged in *blue* (R), polar in *purple* (S, T, N, Q), aromatic in *yellow* (W, F, Y), and aliphatic in *green* (I, L, V, G). The intensity of individual colors is scaled relatively for each assignment class: increased hydrophilicity is brighter, whereas increased hydrophobicity is dimmer.

The N-terminus of SLN is reported to SERCA binding, V_max_ inhibition, K and Ca^2+^ transport/ATP hydrolysis (17,29,31–33,67–71). Here we extend sequence analysis of horse SLN using calculation of the isoelectric point (pI) orthologs: horse, rabbit, mouse, and hu **2**). The online program Isoelectric Poi (IPC), developed by Kozlowski, is an software package that predicts the a peptides and proteins based on a calculation from 13 algorithms, each usi scale for side-chain ionization (pK_a_) (7 IPC algorithm, horse SLN is predicted t basic of the four orthologous peptid (**Figure 2**), with an average pI of 9.1 molecular charge of +1.0 at pH 7.4. In comparison, the predicted pI of human SLN is 8.9 (molecular charge of +1.0 at pH 7.4), rabbit SLN pI is 8.1 (molecular charge of +0.6 at pH 7.4), and mouse SLN pI is 6.5 (molecular charge of 0.0 at pH 7.4). Thus, the molecular charge of SLN orthologs is predicted by IPC to range from highly basic to slightly acidic, as assessed at a physiologically-relevant pH of 7.4 (72). Differences in SLN orthologs (amino acid sequence, charge distribution, and isoelectric point) suggest species-dependent differences in the regulatory interaction of SLN and SERCA (40-42,73) (**Figure 2**). As noted in this regard by Gaudry *et al*. plus Campbell and Dicke, the traditional Met^1^ start codon of SLN is mutated to Leu (ATG to CTG) in extant two- and three-toed sloths, with the mutational event occurring ∼30 million years ago—suggesting that sloth *SLN* transcripts do not translate a functional peptide—although sloth *sln* DNA sequences have remained highly conserved (41, 42). The implication of this remarkable evolutionary conservation is further detailed in the Discussion section below, per the potential of equid *SLN* transcripts (and perhaps sloth also?) to act as functional long non-coding RNA (lncRNA), in light of the supra-abundant level of *SLN* gene transcription in horse gluteal muscle (**Figure 1**) and the highly conserved sloth DNA sequences without the traditional start codon (41, 42).

For additional structural comparison of SERCA regulatory peptides, Valberg *et al*. used a combination of Sanger DNA sequencing, genomic DNA mining, and RNA-seq to deduce the amino acid sequence of horse PLN (inhibitor), horse MRLN (inhibitor), and horse DWORF (activator) (40, 64). Here we extend these analyses by using the IPC algorithm (72) to calculate the pI and molecular charge (pH 7.4) of these three regulatory peptides, including ortholog comparisons. The horse *PLN* transcript encodes a sORF predicted to be translated as a basic transmembrane peptide with pI = 8.4 and molecular charge of +1.7 at pH 7.4. The horse PLN peptide comprises 52 residues, which is the same length as rabbit, mouse, and human PLN, with 98–100% sequence identity among the four orthologs (40). The horse *MRLN* transcript encodes a sORF predicted to be translated as an acidic transmembrane peptide with pI = 4.3 and molecular chare of –2.5 at pH 7.4. The horse MRLN peptide comprises 46 residues, which is the same length as rabbit, mouse, and human MRLN, with ∼80% sequence identity among the four orthologs (40). The horse *DWORF* transcript encodes a sORF predicted to be translated as an acidic transmembrane peptide with pI = 6.1 and molecular chare of –0.5 at pH 7.4. The horse DWORF peptide comprises 35 residues, which is similar to the length of rabbit, mouse, and human DWORF (34–35 residues), with ∼80% sequence identity among the four orthologs (40). Thus, the horse genome encodes the SERCA regulatory peptides PLN, MRLN, and DWORF that exhibit similarities in length, sequence, and charge within each specific peptide-type. On the other hand, the four SERCA regulatory peptides SLN, PLN, MRLN, and DWORF exhibit distinct differences when cross-compared per peptide (i.e., paralogs), particularly in the myosolic regions: N-terminal length, molecular charge, and proposed regulatory sites. Within this key set of SERCA subunits, there appears to be no corre charge and mode of regulati with a pI of 9.1, PLN is an in MLRN is an inhibitor with a is an activator with a pI investigation is needed int peptides, including the roles tissue-specific expression, modification(s), in order to el on Ca^2+^ transport regulat contractility.

### Amino acid sequence analysis of SERCA1 orthologs

The deduced amino acid sequence of horse SERCA1a (adult fast-twitch muscle isoform) encoded by the horse *ATP2A1* gene is 993 residues, with 95% identity with rabbit, mouse, and human SERCA1a, which encode 994-residue proteins (see amino acid sequence alignment in **Figure S1**^3^, plus a compiled list of residue variations in **Table S1**^3^). The four SERCA1 orthologs are acidic proteins (993/994 residues) (**Figure S1**) with pI of 5.0–5.1 and molecular charge of –38 to –39 at pH 7.4, as predicted by IPC (72). Four variations in the amino acid sequence of horse SERCA are noted here (primary sequence analysis), and then assessed by molecular modeling and molecular dynamics (MD) simulation, as described below.

(1) Horse SERCA1a encodes a 1-residue deletion in the nucleotide-binding (N) domain, missing the S504 residue encoded by consensus sequences such as rabbit, mouse, and human SERCA1a; thus, horse SERCA1a (993 amino acids) is one residue shorter than rabbit, mouse, and human orthologs (994 amino acids). (2) Horse and mouse SERCA1a encodes V54 on transmembrane helix (TM) 1, whereas rabbit and human orthologs encode I54 on TM1. The position of residue 54 on TM1 is proximal to the cytosolic Ca^2+^-entry portal that is lined by four acidic residues: E55, E58, and D59 on TM1, plus E108 on TM2 (74-77). (3) Horse SERCA1a encodes V85 on TM2, while rabbit, mouse, and human SERCA1a encode I85. This conservative variation in aliphatic amino acids (V/I) at position 85 occurs at and near residue pairs predicted to mediate regulatory interactions of the dimeric complex from rabbit muscle: I85-SERCA1 and Y31-SLN; F88-SERCA1 and Y29-SLN; F92-SERCA1 and R27-SLN (31,78–80). (4) Horse SERCA1a encodes residue E959, with a negatively-charged sidechain, at the analogous position 960 in orthologs which encode a positively-charged sidechain: K960 in rabbit and mouse SERCA1a, and R960 in human SERCA1a. The analogous position of 959/960 residues of SERCA1a orthologs is located between TM9 and TM10, on a luminal loop (L9–10) in the polar/charged region of the water/phospholipid interface (81).

### Molecular modeling and structural analysis of horse SERCA

To analyze the structural implications of the variations among the amino acid sequences of SERCA orthologs, we generated a structural homology model of horse SERCA1a (**Figure 3**) using the x-ray crystal structure of rabbit SERCA1a in the E1•Mg bound structure with PDB accession ID <u>3W5A</u>^4^ (78, 79). Using this molecular model for comparative structural analysis, we observed that horse and rabbit SERCA1a encode residue variations (*purple spacefill*) that are clustered in peripheral regions of SERCA: surface loops of the actuator (A) domain (comprising residues 1–43 and 124–242), surface loops of the N-domain (comprising residues 359–602), large luminal loop L7–8 (residues 858–895), small luminal loop L9–10 (residues 947–964), and the myosolic half of TM10 (residues 965–985) (**Figure 3**). Four variations in the primary sequence of the four SERCA1 orthologs are assessed here.

**Figure 3.**
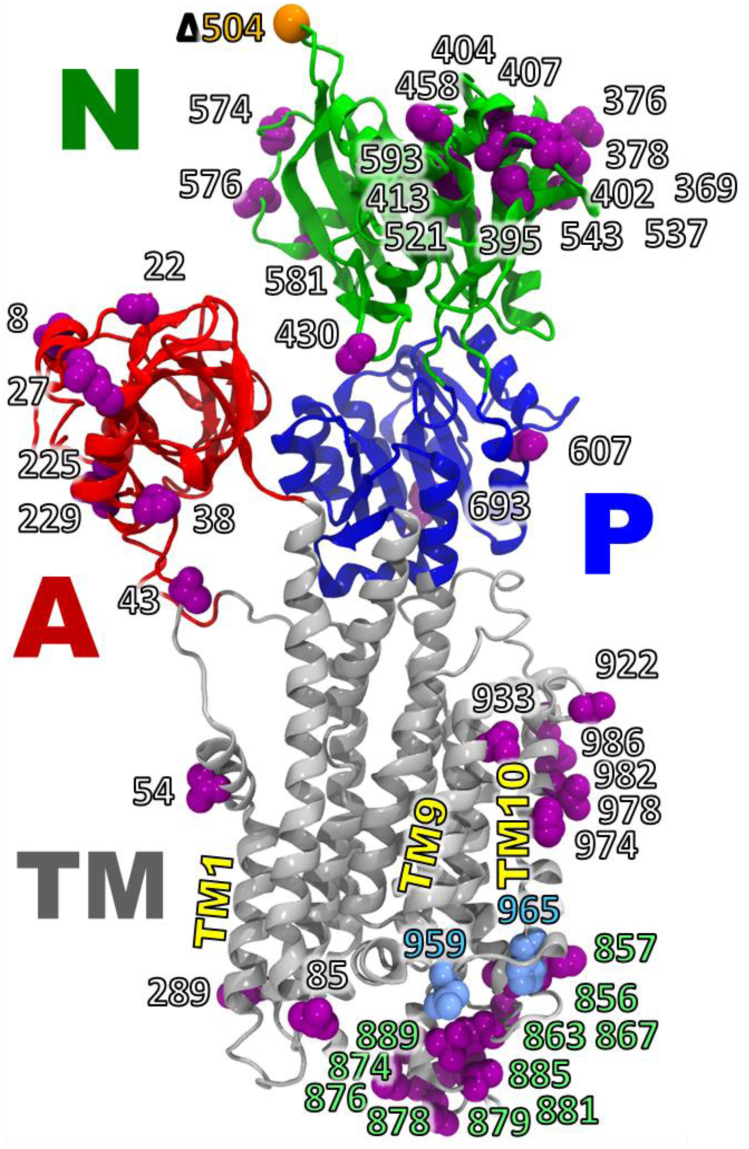
Molecular model of horse SERCA. A homology model of horse SERCA was constructed using the x-ray crystal structure of SERCA purified from rabbit fast-twitch muscle: PDB ID 3W5A^4^ (78). Horse SERCA is rendered in *cartoon* representation with the actuator domain (A) in *red*, the nucleotide-binding domain (N) in *green*, the phosphorylation domain (P) in *blue*, and the transmembrane domain (TM) in *grey*. Residues differing between horse and rabbit orthologs are shown as *purple spacefill*, except for horse residues E959 and H965 on luminal loop L9–10, which are shown in *light blue*. Residue differences in the neighboring luminal loop L7–8 are indicated using *green labels*. The residue deletion at position 504 in horse SERCA1 is shown as an *orange sphere*. Additional detail on SERCA1 sequence variations are shown in **Figure S1** (amino acid alignment of horse, rabbit, mouse, and human orthologs) and **Table S1** (reported functional effects of site-directed mutagenesis at residue positions that show ortholog sequence variation).

(1) Horse SERCA1a encodes a 1-residue deletion at sequence number 504 (Δ504) as compared to rabbit, mouse, and human orthologs, which encode S504 located on a solvent-exposed loop that connects helices N8 and N9 of the N-domain (**Figure 3, Figure S1, Table S1**). The single-residue deletion in horse SERCA1 likely has no significant functional or structural effect, as suggested by (a) bovine SERCA1a (993 residues) encodes the same Δ504 deletion (82), (b) mammalian SERCA2a orthologs encode a proximal deletion (Δ506) located in the same N-domain capping loop (83, 84), and (c) insertion of green fluorescent protein (GFP) at position 509 in this N-domain loop of canine SERCA2a has no effect on enzyme function (85–87).

(2) In the molecular model of horse SERCA1a (**Figure 3**), residue V54 on TM1 is lipid-facing and located in the predicted water–bilayer interface of the myosolic phospholipid leaflet (**Figure S2*A***), similar to the location of residue I54 of rabbit SERCA1a in the x-ray crystal structure PDB ID <u>3W5A</u>^4^ (78). However, there are two reported differences in SERCA activity encoding a V or I residue variation at position 54: (a) in rabbit SERCA1a, the site-directed mutant I54V exhibits decreased Ca^2+^-activated ATPase activity when expressed in COS cells (88), and (b) in human SERCA2b, the spontaneous I54V variation is associated with Darier’s skin disease (89).

To further examine possible functional effects due to the location and identity of residue 54, we reviewed the structural position of residue I54 in rabbit SERCA1a crystalized in three additional structural states assigned to intermediates in the enzymatic cycle: E2•TG (Ca^2+^-free enzyme stabilized by thapsigargin), E1•2Ca (enzyme with two bound Ca^2+^ ions), and E1P•2Ca•ADP (covalent phospho-Asp enzyme with bound ADP and two Ca^2+^). Structural analysis of rabbit SERCA1a indicate that the I54 sidechain in the E2•TG and E1P•2Ca•ADP structures (PDB ID <u>1IWO</u>^4^ and <u>2ZBD</u>^4^, respectively) is located in the phospholipid bilayer at the water–headgroup interface of the myosolic leaflet (75, 90), similar to the position of the I54 sidechain in E1•Mg^2+^ crystal structure of rabbit SERCA1a and the V54 sidechain in E1•Mg^2+^ model of horse SERCA1a (**Figure S2*A***). On the other hand, in the E1•2Ca crystal structure of rabbit SERCA1a with PDB ID <u>1SU4</u>^4^, the rabbit I54 sidechain is located in the interface between TM1 and TM3, whereby rabbit I54 makes van der Waals contacts with the sidechains of residues W50 and V53 on TM1, plus the C_α_ atom of G257 on TM3 (91). For the model of horse SERCA1a in E1•2Ca, the V54 sidechain may not reach far enough to contact the C_α_ atom of G257, thereby decreasing van der Waals force between TM1 and TM3 in horse SERCA1a.

(3) Horse SERCA1 encodes residue V85, a conservative variation at this position, as compared to residue I85 encoded by rabbit, mouse, and SERCA orthologs (**Figure S1**). Residue V85 of rabbit SERCA1 is predicted to interact with residue Y31 of rabbit SLN (80). However, it is unknown if the I85 variation in horse SERCA would perturb regulatory interactions with horse SLN (29 amino acids), which does not encode residue Y31 (40).

(4) Structural comparison of residues E959 and K960 in horse and rabbit SERCA1a (equivalent position in the amino acid sequence of the two orthologs). In the molecular model of horse SERCA1a, residue E959 (*light-blue spacefill*, **Figure 3**) is located on the short luminal loop L9–10 between TM9 and TM10 segments (**Figure S3B**). Modeling of SERCA1 predicts that horse E959 and rabbit K600 sidechains show different positions near the water–phospholipid bilayer luminal interface due to opposite electrostatic force (repulsive versus attractive) with residue E894/E895 on TM8, per horse and rabbit residue numbering, respectively (**Figure S3B**). Analysis of available crystal structures of rabbit SERCA1a shows residue K960 on L9–10 forming a salt bridge with rabbit residue E895 on TM8 (**Figure S2**). Horse residue E959 does not form an equivalent salt bridge to nearby residues; instead, horse residue E959 is part of a negatively-charged patch, and together with E894, could possibly bind a cation in the membrane-water interface in the luminal side of SERCA. Differences in the electrostatic potential surface in luminal loops L7–8 and L9–10 have been proposed to differences in SERCA activity between rabbit and cow SERCA1a (82). The short L9–10 luminal loop interacts with the large L7–8 luminal loop between TM7 and TM8 segments (**Figure 3**, *green* labels), which in bovine SERCA1a is predicted to decrease the maximal enzyme velocity (V_max_) by –35%, as compared to rabbit SERCA1a (82). For another species-specific comparison, chicken SERCA1a encodes T960 at this position on loop L9–10, and the engineered site-directed mutant T960A partially uncouples Ca^2+^ transport from ATPase activity for chicken SERCA1a, producing a lower efficiency of pump energetics, i.e., a decreased Ca^2+^/ATP coupling ratio, less than the optimal ratio of two (92).

### Atomistic molecular dynamics simulations of horse SERCA in a hydrated lipid bilayer

The structural model of horse SERCA1 was generated by homology modeling *in vacuo*. To validate the structural model of horse SERCA, we used the NAMD2 program to perform all-atom MD simulations of horse SERCA embedded in a hydrated phospholipid bilayer composed of 1,2-dioleoyl-sn-glycero-3-phosphocholine (DOPC). Three independent MD simulations indicated that the molecular model of horse SERCA1 is stable for 200 ns, thus demonstrating the fidelity of structural model construction. Next, the 200 ns trajectories of horse SERCA were compared to those previously analyzed for rabbit SERCA, as performed with similar computational input parameters, finding that residue V54 of horse SERCA show similar structural positioning and side-chain dynamics as residue I54 of rabbit SERCA (76,77,93–95). Thus, molecular modeling and MD simulation were used to correlate an amino acid sequence variation and predicted protein structural dynamics. However, initial computational analyses did not provide insight into activity differences due to the position or identity of residue 54, as identified by site-directed mutagenesis (horse SERCA1) or Darier’s disease variation (human SERCA2) (88, 89). Luminal loops of SERCA are also implicated in the control of activity of SERCA isoforms and species; for example, comparative structural studies of cow SERCA1 that identified luminal loop L7–8 interacting with luminal loops L3–4 and L9–10 (PDB ID <u>3TLM</u>^4^), with loop positionings proposed to lower the V_max_ of the bovine ortholog, as compared to rabbit SERCA1 (82). We predict that additional molecular modeling and comparative MD simulations of horse SERCA, using orthologous SERCA proteins with and without SLN as bound subunit, will yield functional and structural hypotheses that predict mechanisms for the enhanced Ca^2+^ transport of SERCA1 in horse muscle, as identified experimentally and reported below.

### An improved fractionation protocol increases the purity of SR vesicles isolated from horse muscle

To investigate SERCA activity and regulation in horse gluteal muscle, we aimed to develop further our protocol for purifying a subcellular microsomal fraction enriched in SR vesicles (57). Tissue samples were collected from the middle gluteal muscle of four horses: three Quarter Horses and one Thoroughbred (see **Experimental Methods**). The middle gluteal muscle of these two horse breeds has a slightly higher proportion (∼90–95%) of fast-twitch muscle fibers (96) than the six control Arabian horses (∼85%) studied by qRT-PCR of SERCA regulatory peptides and Ca^2+^ release proteins (40), by RNA-seq of cytosolic redox proteins (64), and by RNA-seq of SERCA gene transcription and regulatory peptides here (**Figure 1**). We also reviewed previous reports on isolation of SR from horse muscle (53–57), summarized in (**Table 1**). In general, previous horse SR studies have typically isolated unfractionated SR vesicles comprised of longitudinal and junctional SR using a high-speed centrifugation step (ranging from 38,000 to 165,000 × *g*), preceded by a clarification step of moderate-speed centrifugation (ranging from 8,000 to 12,000 × *g*) (**Table 1**). However, these previous studies do not show electrophoretic analysis of the protein profile of horse SR preparation, nor immunoblot analysis of SERCA protein content. Instead, these previously studies rely on Ca^2+^-ATPase and Ca^2+^ transport assays as an indicator of horse SERCA in the SR preparations, which exhibit only 2–28% the activity of rabbit SERCA in unfractionated SR vesicles from rabbit skeletal muscle (**Table 1**). Similarly, horse SR shows low activity compared to unfractionated SR from rat skeletal muscle, with horse SERCA having only 12% the activities (Ca^2+^ ATPase and Ca^2+^ transport) compared to rat SERCA (55). The mechanistic origin of low SERCA activity in horse SR has been unclear (55). Thus, our goal in the present study was to increase the enrichment of SR vesicles purified from horse gluteus, including validation by key parameters such as a high content of SERCA protein, a high level of SERCA activity, and a low content of contaminating proteins and enzyme activities from other horse muscle fractions and membranes.

**Table 1.** Reported Ca^2+^-ATPase and Ca^2+^ transport activities in SR vesicles purified from horse fast-twitch muscles. Published results of horse SERCA activities are listed from studies utilizing unfractionated SR vesicles purified from horse muscle, as assessed using ionophore-facilitated ATPase assay or oxalate-facilitated transport assay. Our improved protocol for isolating horse SR vesicles from gluteal muscle (*this study*) provided a Ca^2+^-activated ATPase activity of 4.0 ± 0.39 IU at 37 °C and a Ca^2+^ transport activity of 4.0 ± 1.0 IU at 25 °C. Prior to this study, the maximum reported activities of horse SR vesicles were Ca^2+^-ATPase of 0.73 ± 0.14 IU at 37 °C and Ca^2+^ transport of 0.55 ± 0.16 IU at 37 °C. For comparison, the Ca^2+^-ATPase and Ca^2+^ transport activities of unfractionated SR vesicles purified from rabbit fast-twitch muscle are also listed, assayed under the same conditions utilized for horse SR vesicles in this study.

| Horse SR fractionation protocols | Centrifugation Spins ($\text{g-force} \times 1000$)<sup>a</sup> | | $\text{Ca}^{2+}$-ATPase assay + A23187<sup>b</sup> (IU) | $\text{Ca}^{2+}$ transport assay + oxalate<sup>c</sup> (IU) |
| --- | --- | --- | --- | --- |
|  | Pre-clear | SR collect |  |  |
| **this study<sup>d</sup>** | **4** | **10** | **$4.0 \pm 0.39$** | **$4.2 \pm 0.66$** |
| *Am J Vet Res* 2000<sup>e</sup> | 2.6 | 140 | $0.71 \pm 0.42$ | n.r.<sup>l</sup> |
| *Equine Vet J Suppl* 1998<sup>f</sup> | 10 | 38 | $0.16 \pm 0.01$ | $0.18 \pm 0.02$ |
| *J Anim Sci* 1995<sup>g</sup> | 12 | 165 | $0.16 \pm 0.01$ | $0.19 \pm 0.02$ |
| *J Anim Sci* 1995<sup>g</sup> | 7 | 160 | unsuccessful<sup>k</sup> | n.r.<sup>l</sup> |
| *Res Vet Sci* 1993<sup>h</sup> | 8 | 12 | n.r.<sup>l</sup> | n.r.<sup>l</sup> |
| *J Appl Physiol* 1989<sup>i</sup> | 10 | 38 | $0.73 \pm 0.14$ | $0.55 \pm 0.16$ |
| **\*rabbit SR: this study<sup>j</sup>** | **12** | **23** | **$7.6 \pm 0.45$** | **$2.4 \pm 0.45$** |

<sup>a</sup>The g-force of centrifugation spins (pre-clear and SR collection steps) are reported in relative centrifugal force ( $\text{g-force} \times 1000$ ).
<sup>b</sup>A23187, a $\text{Ca}^{2+}$ ionophore, was added to the ATPase assay in order to release accumulation of intravesicular $\text{Ca}^{2+}$ .
<sup>c</sup>Oxalate, a $\text{Ca}^{2+}$ -precipitating anion, was added to the transport assay in order to trap accumulation of intravesicular $\text{Ca}^{2+}$ .
<sup>d</sup>SR vesicles from horse gluteal muscle (see also **Figure 4**, **Figure 5**, **Figure 6**).
<sup>e</sup>Ward et al., *Am. J. Vet. Res.* 2000 (57)
<sup>f</sup>Wilson et al., *Equine Vet. J. Suppl.* 1998 (56)
<sup>g</sup>Wilson et al., *J. Anim. Sci.* 1995 (55)
<sup>h</sup>Beech et al., *Res. Vet. Sci.* 1993 (54)
<sup>i</sup>Byrd et al., *J. Appl. Physiol.* 1989 (53)
<sup>j</sup>SR vesicles from rabbit fast-twitch skeletal muscle purified by slight adaptation of the protocol from Ikemoto et al., *J Biol Chem* 1971 (50) (see also **Figure 6** and **Figure S3**).
<sup>k</sup>SR purification and $\text{Ca}^{2+}$ -ATPase activity assay were reported as “unsuccessful” (55).
<sup>l</sup>n.r. = not reported.

As previously, parameters of differential centrifugation were used here to examine subcellular fractions from horse muscle, in the search for a microsomal membrane preparation enriched in longitudinal and junctional SR vesicles (unfractionated SR). We examined two muscle fractions isolated at common centrifugal speeds that often yield membrane vesicles, albeit a combination of types such as SR, nuclear, and/or mitochondrial microsomes, depending on the muscle type, species, and prep solutions. First, we isolated a horse muscle fraction using a centrifugation step at 10,000 × *g*, which was preceded by a homogenate clarification step using centrifugation at 4,000 × *g*, thereby yielding a pellet which we termed the 10K fraction (10KP) (see *green box* in **Figure 4**). The 10KP pellet from horse muscle fractionation was beige, soft, and easily resuspended, indicative of a centrifugal pellet comprised of membrane vesicles. The yield of the 10KP fraction from 200 g of horse gluteus was ∼200 mg protein, i.e., ∼1 mg protein/g tissue. Second, we isolated a horse muscle fraction using a centrifugation step at 100,000 × *g*, which was preceded by the centrifugation step at 10,000 × *g*, thereby yielding a pellet which we termed the 100K fraction (100KP) (see *red box* in **Figure 4**). The 100KP pellet from horse muscle fractionation was colorless, translucent, hard, and difficult to resuspend, suggestive of a centrifuged pellet comprised of glycogen granules (97). The yield of the 100KP fraction from 200 g of horse gluteus was ∼50 mg protein, i.e., ∼0.25 mg protein/g tissue.

**Figure 4.**
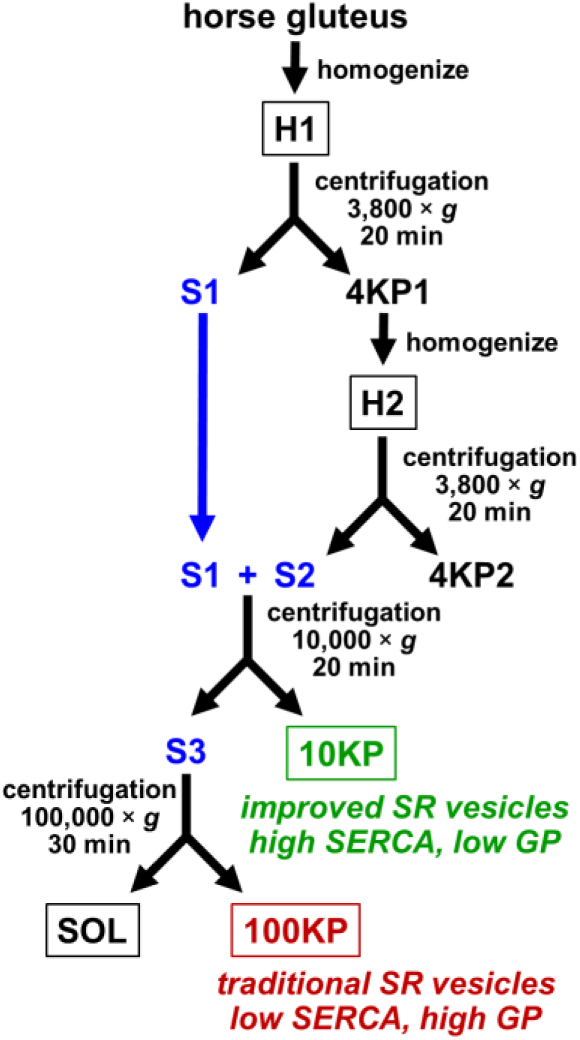
Improved protocol for purification of SR vesicles from horse muscle. Five fractions from horse muscle were isolated and characterized biochemically, as identified by *boxed text*: homogenate 1 (H1), homogenate 2 (H2), 10,000 × *g* pellet (10KP), 100,000 × *g* pellet (100KP), and 100,000 × *g* supernatant (SOL). Intermediate fractions include supernatants 1–3 (*blue text*).

For comparative biochemical assays of horse and rabbit SERCA, homogenization and differential centrifugation of rabbit muscle were also used to purify a microsomal fraction enriched in longitudinal and junctional SR vesicles (unfractionated preparation). The method of Ikemoto *et al*. (1971) was used to purify SR vesicles from rabbit skeletal muscle (50), with slight modifications (see **Experimental Methods**). Rabbit SR vesicles were collected using a centrifugation step at 23,000 × *g*, which was preceded by a homogenate clarification step using centrifugation at 12,000 × *g*, thereby yielding a pellet which we termed here the 23KP fraction, i.e., unfractionated SR vesicles (see *green box* in **Figure S3*A*** and gel lanes labeled *SR* in **Figure S3*B***). The 23KP pellet comprised of SR vesicles from rabbit muscle was beige, soft, and easily resuspended, as expected for a centrifuged pellet comprised of membrane vesicles. The yield of unfractionated SR vesicles from rabbit muscle was ∼1 mg of SR protein per g of tissue (N = 6 preps). Rabbit SR vesicles isolated by this method are highly purified, well characterized, and widely utilized, with robust SERCA activity (98).

### Immunoblotting and Coomassie staining identify SERCA and glycogen phosphorylase as co-migrating gel-bands that are differentially enriched in the 10KP and 100KP fractions, respectively

SDS-PAGE and immunoblot analysis were used for initial characterization of five subcellular fractions from horse muscle: homogenate 1 (H1), homogenate 2 (H2), membrane pellets 10KP (4,000–10,000 × *g*) and 100KP (10,000–100,000 × *g*), and the 100,000 × *g* supernatant (SOL) (**Figure 4**, **Figure 5**). Immunoblotting for SERCA detection utilized mouse monoclonal antibody (mAb) anti-SERCA1 VE12_1_G9, which identified SERCA1 protein in all five fractions from horse gluteal muscle, with the 10KP fraction showing many-fold greater SERCA1 content than the other four horse muscle fractions (**Figure 5*A***). Semi-quantitative immunoblotting using mAb VE12_1_G9 (**Figure S4**) demonstrated that the level of SERCA1 protein in the 10KP fraction is enriched 11 ± 1.1 fold compared to the initial H1 homogenate of horse gluteal muscle, by defining the SERCA expression of the H1 fraction as 1.0 (± 0.37) (**Figure 5*A*, Figure S4**). The 10KP fraction also showed 5.0 ± 0.54 fold greater level of SERCA1 protein than the 100KP fraction, and the 10KP fraction showed ≥ 10-fold level of SERCA1 protein than the H2 and SOL fractions (**Figure 5*A*, Figure S4**). Thus, immunoblot analysis of horse muscle fractions demonstrated high enrichment of SERCA1 protein in the 10KP fraction; a unique observation, since the majority of reported horse SR preps did not include this centrifugal cut, but instead collected SR vesicles that pelleted at ≥ 10,000 × *g* (**Table 1**). Thus, immunoblotting and Coomassie densitometry indicate that the 10KP fraction (4,000–10,000 × *g* cut) from horse muscle is an improved membrane vesicle preparation for functional and structural assays of SERCA.

**Figure 5.**
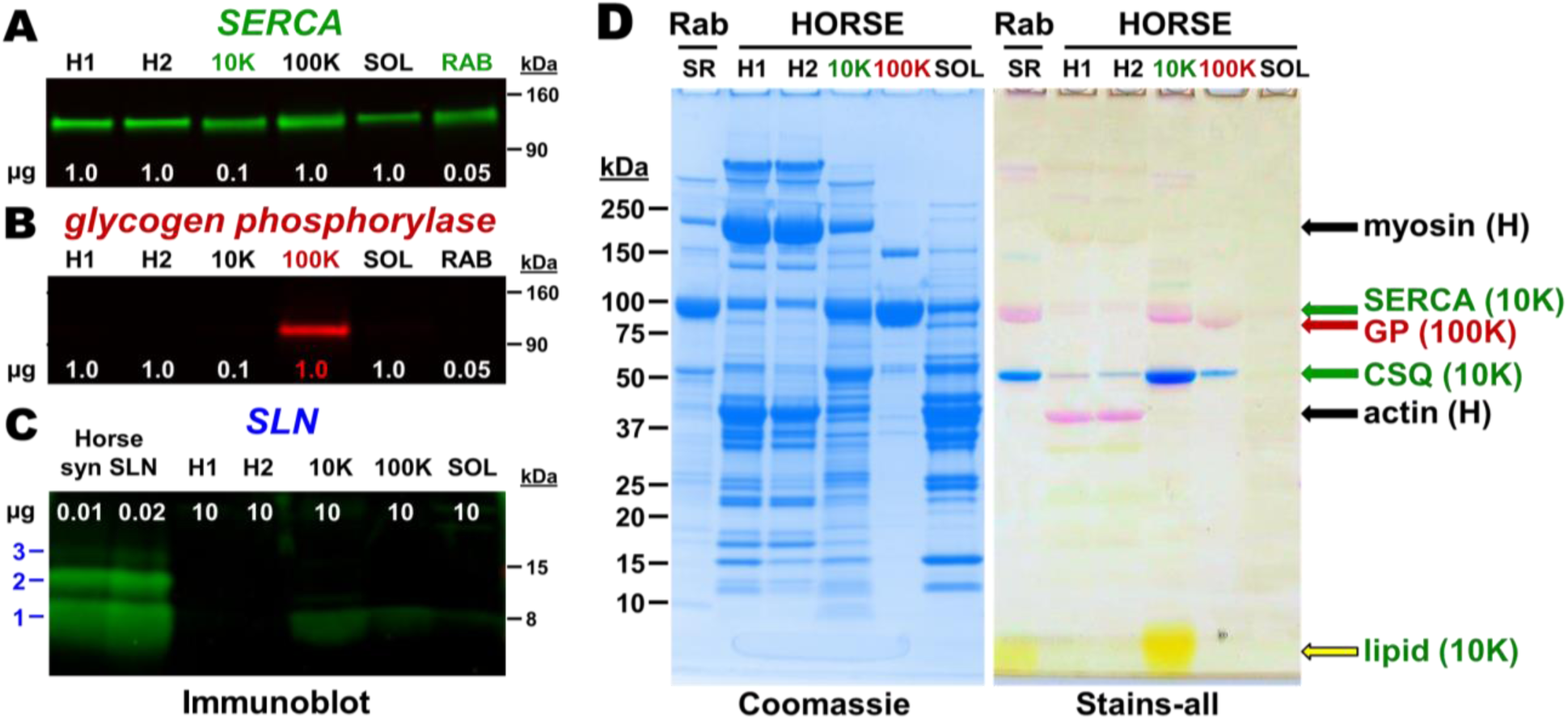
Immunoblot and gel-stain analyses of horse muscle fractions. Five horse muscle fractions were assayed: H1, H2, 10KP, 100KP, and SOL. Samples were electrophoresed through 4–15% Laemmli gel for SERCA and GP blots, and 4‒20% Laemmli gel for SLN blot. The protein load (µg) in each lane is indicated in *white text*. Protein molecular mass markers are indicated on the *right*. *A*, SERCA immunoblot using mAb VE121G9 (primary) and goat anti-mouse-IgG pAb labeled with 800-nm fluorophore (secondary). SR vesicles isolated from rabbit skeletal muscle (RAB) were used as a quantitative positive control (prep flowchart and gel analysis of fractions in Figure S4). *B*, GP immunoblot using mAb ab88078 (primary) and goat anti-mouse-IgG pAb labeled with 680-nm fluorophore (secondary). Rabbit SR was also assayed (RAB). *C*, SLN immunoblot using anti-horse-SLN pAb GS3379 (primary) and goat anti-rabbit-IgG pAb labeled with 800-nm fluorophore (secondary). Synthetic horse SLN was used as a positive control (0.01–0.02 µg), as compared to horse muscle fractions (10 µg). Molecular species of SLN (monomer, dimer, and trimer) are indicated on the *left*. *D*, Gel-stain analysis of horse muscle fractions. Samples were electrophoresed through 4‒15% Laemmli gels and stained with Coomassie blue (*left*) or Stains-all (*right*). The amount of horse protein loaded per lane is 30 µg for homogenate 1 (H1) and homogenate 2 (H2), 20 µg for the 10,000 × *g* pellet (10K), 10 µg for the 100,000 × *g* pellet (100K), and 30 µg for the 100,000 × *g* supernatant (SOL). Rabbit SR vesicles (RAB) were used as positive control (10 µg protein load). Gel bands are identified by arrows to the *right*, along with the predominant muscle fraction for each. Protein molecular mass markers are indicated on the *left*.

Rabbit muscle expresses almost exclusively the SERCA1 isoform, with undetectable amounts of SERCA2 or SERCA3 proteins (99), while horse gluteal muscle expresses predominantly the *ATP2A1* transcript (**Figure 1*A***). Thus, the collection of SERCA proteins (mostly SERCA1) expressed in horse and rabbit muscle will hereafter be referred to as “<u>SERCA</u>” for simplicity.

Immunoblot analysis of glycogen phosphorylase (GP) in horse muscle was also performed, because GP is a common contaminant in SR preparations (47), and because of the apparent nature of the 100KP pellet, which was clear, glassy, and sticky, i.e., reminiscent of glycogen granules, instead of membrane vesicles. GP is a soluble protein that remains bound to glycogen granules during muscle homogenization and fractionation by differential centrifugation. Horse muscle is rich in glycogen, comprising 1.5–2.0% of wet weight muscle (96). GP and SERCA co-migrate at 100 kDa on SDS-PAGE, thus leading to possible misidentification of GP as SERCA in muscle fractions, and/or not accounting for GP protein contribution at 100 kDa when performing Coomassie densitometry for SERCA in various muscle fractions. This problem was recognized early in the rabbit SR field, and protocols were developed to separate SR membranes from glycogen granules from an initial glycogen-sarcovesicular fraction (47). Our Western blotting indicates that the 100KP fraction is highly enriched in GP, as compared to initial homogenates H1 and H2 of horse gluteal muscle (**Figure 5*B***). Thus, GP immunoblotting provides an additional indicator, along with the appearance and physical nature of the 100KP pellet, that glycogen granules are a considerable component of the 100KP fraction from horse gluteal muscle.

Horse muscle fractions obtained by differential centrifugation were also analyzed using electrophoretic analysis and in-gel staining by Coomassie blue (**5*D***). Gluteus homogenates H1 and H2 express abundant Coomassie-stained bands that migrated at 200 kDa (myosin) and 40 kDa (actin), plus a moderately-abundant band at 100 kDa band, likely a mix of SERCA, GP, and other proteins. The 10KP fraction (4,000–10,000 × *g* pellet) and the 100KP fraction (10,000–100,000 × *g*) from horse muscle express abundant band(s) at ∼100 kDa, which appear to be an unresolved doublet comprising one or two proteins. Although Coomassie blue is utilized frequently to identify protein bands by gel mobility, this dye cannot be utilized to identify protein bands that co-migrate on SDS-PAGE. On Laemmli-type gradient gels, SERCA runs at 100 kDa, as does GP, a common contaminant in SR vesicles (47). Thus, traditionally-studied preparations of horse SR which are isolated at higher centrifugal forces (between 10,000 and 165,000 × *<u>g</u>*) (**Table 1**) are probably non-optimal for examining Ca^2+^-ATPase and Ca^2+^ transport activities due to enrichment of GP and glycogen granules, plus de-enrichment of SERCA protein (**Figure 5**). Interestingly, the 10KP fraction of horse muscle expresses a Coomassie-stained band that migrates faster, i.e., lower molecular mass, than any Coomassie-stained band in rabbit SR (**Figure S5**). The identity of this low molecular mass protein (< 10 kDa) in horse 10KP is unknown. We propose that the 10KP fraction (4,000–10,000 × *g* cut) from horse muscle is an improved membrane vesicle preparation enriched in SR vesicles that will provide more effective assays of SERCA function, structure, and regulation.

We utilized quantitative immunoblotting and Coomassie densitometry to compare the relative amount of SERCA protein in SR vesicles from horse versus rabbit muscle. For standardization, the SERCA content of SR vesicles from rabbit fast-twitch (white) muscle has been determined to be 55‒70% of total protein (weight/weight), i.e., 5.0‒6.4 nmol SERCA/mg rabbit SR protein (98,100–102). Immunoblotting with anti-SERCA1 mAb VE12_1_G9 demonstrated that SR vesicles from horse gluteal muscle express ∼35% the relative amount of SERCA1 as compared to rabbit SR (**Figure 5*A*, Figure S5**), although this is probably a slight underestimate of total SERCA content, since horse SR (10KP) also expresses a small amount of *ATP2A2* (SERCA2) (**Figure 1**), which is incompatible with the use of anti-SERCA1 mAb VE12_1_G9. Coomassie densitometry demonstrated that SR vesicles from horse gluteal muscle express ∼55% the relative content SERCA compared to rabbit SR (**Figure S4**), although this is probably a slight overestimate, since horse SR (10KP) may contain a small amount of GP and/or additional proteins that co-migrate with SERCA at ∼100 kDa on SDS-PAGE (**Figure 5*B,D***). Despite the minor limitation of each gel-based method, quantitative immunoblotting and Coomassie densitometry are two orthogonal assays that provide similar results, indicating that horse SR vesicles express 35‒55% the SERCA content of rabbit SR vesicles. Assuming that (i) the expression level of SERCA in rabbit SR is 6.0 nmol mg SERCA/mg SR proteins, a common value, and (ii) horse SR contains (a relative median value of) 45% the amount of SERCA protein as rabbit SR, then the density of SERCA expressed in horse SR is 2.7 nmol SERCA/mg SR protein. Multiple SR preps from individual horses and rabbits were assayed by densitometry and immunoblotting, demonstrating that there are only slight variations in the high content of key SR proteins (e.g., SERCA and CASQ) and in the low content of contaminating proteins (e.g., myosin) within the type of SR preparation from each animal (**Figures S4, S5, S6, S10**). Our next goals are to increase the purity of horse SR vesicles and to increase the number of SR preps purified from horse and rabbit muscles, in order to account for individual variation(s) in muscle proteogenomics. In this initial report, we detected a small amount of uncertainty in measured SERCA content, yet as demonstrated below, this uncertainty does not affect the reported results; for example, horse SR vesicles have enhanced oxalate-facilitated Ca^2+^ transport compared to rabbit SR vesicles, as determined without normalization to SERCA content.

### Stains-all identifies calsequestrin and phospholipids as SR markers enriched in the 10KP fraction, but not the 100KP fraction

Additional electrophoretic analysis was performed to corroborate the identification of SR distribution in horse muscle fractions. Calsequestrin (CASQ) is a high-capacity Ca^2+^-binding protein localized to junctional SR with RYR; the CASQ1 isoform is expressed in horse gluteus and rabbit muscles (40, 103). A useful tool to identify CASQ and phospholipids on SDS-PAGE is the cationic carbocyanine dye Stains-all (**Figure 5*D***). Stains-all is a metachromatic dye (i.e., having an environmentally-sensitive absorbance) which stains most proteins pink or red, yet also provides unique staining of proteoglycans as purple, phospholipids as yellow, and highly-acidic proteins as blue (e.g., high-capacity Ca^2+^-binding proteins such as CASQ) (9,103,104). Electrophoretic analysis with in-gel staining by Stains-all illustrates the protein profile of horse muscle fractions. In H1 and H2, actin stains as an abundant pink band at 40 kDa. In the 10KP, there are high abundance band(s) staining pink at 90–100 kDa (SERCA and/or GP), two low abundance bands staining faint purple at >250 kDa in the 10KP fraction (possibly ‘CASQ-like’ proteins) (9), and one abundant band staining intensely-blue at 50 kDa, i.e., CASQ (**Figure 5*D***). In the 100KP, there are high abundance band(s) staining pink at 85–95 kDa (GP and/or SERCA) and one low abundance band staining intensely-blue at 50 kDa (CASQ). Thus, the abundant CASQ band in the 10KP fraction serves as a subcellular marker of SR vesicles. Stains-all also detected the expression of CASQ at 55 kDa in fractions H1 and H2 from horse muscle, demonstrating dye sensitivity and specificity. Comparison of Stains-All and Coomassie gels indicated that CASQ is the second-most abundant protein in the 10KP fraction from horse muscle, following SERCA (**Figure 5*D***).

In horse muscle fractions, CASQ was less abundant in 100KP than 10KP, indicating preferential enrichment of SR vesicles in the 10KP fraction. Only a small amount of CASQ staining was detected in the 100KP fraction from horse muscle, in spite of an abundant ∼100 kDa Coomassie-staining protein band (here identified as GP by immunoblotting in **Figure 5*B***, yet often mis-identified as SERCA, due to co-migration). In rabbit muscle fractions, Stains-All identified CASQ1 in H1, H2, P3 (unwashed 23KP), and P5 (washed 23KP) (**S3, Figure 5*D***). CASQ1 was most enriched in the washed 23KP fraction, defined as “rabbit SR vesicles” (**Figure S3B, Figure 5*D***). Coomassie staining of the sets of horse preps (10KP; N = 4) and rabbit preps (washed 23KP; N = 5) utilized in this study indicated that horse SR expresses ∼20% greater CASQ protein than rabbit SR (**Figure S4**). Stains-all gels corroborated similar levels of CASQ protein expression in horse and rabbit SR (**Figure 4*D***), assuming that horse and rabbit CASQ orthologs bind the Stains-All dye with similar affinity and metachromatic effect, per molecular electronegativity. Both horse and rabbit CASQ1 have the same predicted pI of 3.8, as assessed by IPC (72). When normalized to SERCA content, the CASQ/SERCA ratio (protein/protein) in horse SR (10KP fraction) is ∼3-fold greater than the CASQ/SERCA ratio (protein/protein) in rabbit SR (washed 23KP fraction), as determined by Coomassie blue densitometry (**Figure 5*D*, Figure S4**). We propose that the relatively high level of CASQ protein expression in horse gluteus contributes to enhanced SR Ca^2+^ handling and potentially high muscular performance.

Phospholipids, which are stained yellow by Stains-all, migrate near the bromophenol blue dye-front on SDS-PAGE (103). For rabbit muscle, Stains-all demonstrated that phospholipids are most enriched in the P5 fraction (washed 23KP = SR vesicles) (**Figure S4, Figure 5**). For horse muscle, Stains-all demonstrated that phospholipids are enriched in the 10KP fraction from horse muscle, indicating the abundance of SR vesicles in this fraction (**Figure 5**). Stains-all also demonstrated that phospholipids are de-enriched in the 100KP fraction from horse muscle, indicating the deficit of SR vesicles in this fraction. Thus, Stains-all gel identification of CASQ and phospholipids in horse muscle fractions corroborates immunoblotting results demonstrating that SERCA protein expression is enriched in the 10KP fraction. Furthermore, Stains-all gel analysis is consistent with immunoblotting results that identified the 100 kDa band is predominantly GP in the 100KP fraction from horse muscle, instead of SERCA. Thus, the 10KP fraction from horse muscle homogenate shows the most enrichment of three distinct SR markers (SERCA, CASQ, and phospholipids), so we define the 10KP fraction isolated from horse gluteal muscle as “<u>SR vesicles</u>”.

### Horse gluteal muscle expresses a minor level of SLN protein relative to SERCA, as detected by multiple anti-SLN immunological approaches

RNA-seq determined that horse muscle expresses a high level of the *SLN* gene transcript (**Figure 1**). To determine if SLN peptide expression correlates with *SLN* gene transcription, immunoblotting was used to quantitate the expression level of native SLN peptide in horse muscle (**Figure 5*C***). Due to the unique amino acid sequence of horse SLN (**Figure 2**), a custom anti-horse-SLN pAb was ordered (GS3379) using horse SLN residues ^1^MEWRRE^6^ (N-terminal peptide) as immunogen. The pAb GS3379 was produced by four rounds of rabbit immunization, with titer assessments by ELISA assay and a final affinity purification. In addition, chemical synthesis of full-length horse SLN (residues 1–29) was used to produce a positive control and gel-loading standard (**Figure 5*C***, **Figure S6**). On immunoblots, anti-horse-SLN pAb GS3379 detected synthetic horse SLN loaded at 10‒20 ng SLN/lane (**Figure 5*C***) and 2.5‒10 ng SLN/lane (**Figure S6**), thereby validating pAb GS3379 as an avid antibody against horse SLN.

Per immunoblotting of horse muscle fractions, anti-horse-SLN pAb GS3379 showed little to no detection of native SLN in homogenates H1 or H2, nor 100KP or SOL (**Figure 5*C***). However, pAb GS3379 detected native SLN expression in horse SR vesicles (23KP) when loaded at 10 µg SR protein/lane (**Figure 5*C***, **Figure S6**), albeit at a lower intensity level compared to 10 ng of synthetic horse SLN, i.e., SLN comprises < 0.1% of the total protein in horse SR vesicles. Semi-quantitative immunoblot analysis of four horse SR preps using pAb GS3379 and a synthetic horse SLN standard determined that native SLN is expressed at 0.16 ± 0.015 nmol SLN/mg SR protein (**Figure S6**). The expression level of SERCA in horse SR is estimated at 2.7 nmol SERCA/mg SR protein (see above), per content assessed by Coomassie densitometry (**Figure S4**) and SERCA immunoblotting (**Figure 5**, **Figure S5**) using rabbit SR as a relative SERCA standard, with 5.0‒6.4 nmol SERCA/mg SR protein (98,100,101). Thus, quantitative immunoblotting using the anti-horse-SLN pAb 3379 and a synthetic horse SLN peptide standard determined that SR vesicles from horse gluteal muscle express a minor ratio of 0.059 ± 0.005 SLN/SERCA (mol/mol protein).

On immunoblots, the small amount of horse SLN in native SR vesicles migrates as a monomer on SDS-PAGE (**Figure 5*C***, **Figure S6**), as is typically found for rabbit SLN in SR vesicles, and also other orthologs (mouse, dog, and pig) in respective SR vesicles (26,28,78,102). On the other hand, purified SLN (native, recombinant, or synthetic) on SDS-PAGES migrates often as a mix of oligomeric species: abundantly-populated monomers with a variable amount of dimers, and sometimes with lowly-populated trimers and/or tetramers (29,31,105–107) (**Figure 5**, **Figure S9**). Thus, denaturing SDS-PAGE maintains a variable amount of self-association for purified SLN. In biomimetic membranes, dimerization of rabbit SLN has been detected in Sf21 insect cells via baculovirus infection, yet the degree of dimerization was dependent on a combination of conditions such as sample-SDS concentration, solubilization temperature, and gel type (108). Regardless of SDS-PAGE, self-association of SLN into dimers and higher-order oligomers in SR is proposed to play a key role in SERCA regulation, as SLN oligomerization competes directly with the binding of inhibitory SLN monomers to SERCA (31,106,108–110). A similar competition of peptide–pump interactions has been well characterized for the PLN–SERCA system, whereby PLN self-association into pentamers competes with the assembly of PLN monomers and SERCA into the inhibitory hetero-dimeric complex, with the avidity depending on competing binding equilibria that are dynamically controlled by PLN phosphorylation and SERCA Ca^2+^ binding (83,111,112). Thus, oligomerization of synthetic horse SLN on SDS-PAGE suggests that self-association of horse SLN plays a role in the availability of SLN monomers for inhibition of horse SERCA.

Further attempts to detect SLN in horse and rabbit muscle utilized additional commercial and custom-made anti-SLN antibodies. Horse SLN shows high sequence divergence and a 1-residue deletion (ΔS4) at the N-terminus, plus a 1-residue deletion (ΔY30) at the C-terminus (**Figure 2**). The commercial anti-rabbit/mouse/human-SLN pAb ABT13, with immunogen comprising the consensus C-terminus of SLN (residues ^27^RSYQY^31^), successfully detected SLN expression in rabbit SR (0.5 µg SR protein/lane), but pAb ABT13 did not detect SLN in horse SR (10 µg SR protein/lane) (**Figure S7**). The custom anti-rabbit/mouse/human-SLN pAb PFD-1 generated by Desmond *et al*., with immunogen comprising a 7-residue stretch of the consensus C-terminus of SLN (residues ^25^LVRSYQY^31^) (113), successfully detected rabbit SLN expression in SR vesicles (0.1 µg SR protein/lane) and also synthetic rabbit SLN (0.01 µg SLN/lane), but pAb PFD-1 did not detect horse SLN in gluteal SR vesicles (10 µg SR protein/lane) nor synthetic horse SLN (25 ng SLN/lane) (**Figure S8**). The commercial pAb 18-395-1-AP, with immunogen comprising human SLN (residues 1–31), did not detect SLN in rabbit SR (0.5 µg SR protein/lane) nor horse SR (10 µg SR protein/lane) (**Figure S9**). Two additional commercial pAbs N-15 and E-15 (raised against human and mouse SLN, respectively) did not detect rabbit SLN in SR vesicles (2 µg SR protein/lane). In our assessment, the custom anti-horse-SLN pAb GS3379 gives suitable and reproducible immunoblot results for detecting horse SLN. Furthermore, the commercial anti-rabbit/mouse/human-SLN pAb ABT13 and the custom anti-rabbit/mouse/human-SLN pAb PFD1-1, provided by Bloch lab (23, 113), gives suitable and reproducible immunoblot results for detecting rabbit SLN.

When correlating the gene expression of *SLN* versus the protein expression of SLN in horse muscle (normalized per *ATP2A1* or SERCA1, respectively), we find that the high level of *SLN* RNA, with *SLN*/*total*-*ATP2A* = 9.1 ± 3.7 (TPM/TPM) (**Figure 1*C***), produces only an insignificant amount of SLN regulatory peptide, with a protein expression ration of SLN/SERCA = 0.059 ± 0.005 (mol/mol) (**Figure 5**, **Figures S4‒S6**). On the other hand, rabbit skeletal muscle produces a relative gene expression ratio of *SLN*/*total-ATP2A* = 3.0 ± 0.2 (TPM/TPM) (**Figure 1*C***) and a protein expression ratio of 0.75–1.2 SLN/SERCA (mol/mol) (29, 102). Thus, horse muscle lacks correlation of gene and protein expression levels for SLN and SERCA, unlike rabbit muscle. We propose that horse muscle utilizes translational and/or degradation mechanism to mediate a low basal level of SLN protein, and that the high level of *SLN* RNA may possess a cellular function besides translation.

### Horse gluteal muscle expresses an insignificant level of PLN protein compared to SERCA

RNA-seq determined that horse muscle expresses a minor level of the *PLN* gene transcript (**Figure 1**). To determine if PLN peptide expression correlates with *PLN* gene transcription, immunoblotting was used to quantitate the expression level of native PLN peptide in horse muscle (**Figure S10**). The commercial anti-PLN mAb 2D12, with an epitope against the N-terminus of PLN that encodes an identical sequence among horse, rabbit, mouse, and human PLN orthologs, was used to quantitate PLN peptide expression in horse SR. The anti-PLN mAb 2D12 is robust and has been used effectively in multiple types of experiments: immunoblotting, ELISA, and histochemistry, plus functional assays to disrupt PLN inhibition of SERCA in SR vesicles and live cells (114–117). Quantitative immunoblotting with anti-PLN mAb 2D12 and a synthetic PLN peptide standard determined that the 10KP fraction (SR vesicles) from horse gluteal muscle expresses <0.01 PLN/SERCA (mol/mol) (**Figure S10**). The binding affinity of an inhibitory PLN monomer to SERCA is ∼40-fold less than the binding affinity of a PLN monomer for the competing PLN self-association into non-inhibitory homo-pentamers (118), thereby reducing greatly the available amount of functional PLN monomers (≪ 0.01 mol/mol SERCA). Thus, we conclude that the level of PLN peptide expressed in SR vesicles from horse gluteal muscle is insignificant compared to the expression of SERCA protein.

### Ca^2+^-activated ATPase activity by SERCA is enriched in the 10KP fraction, but not the 100KP fraction, from horse muscle

SERCA activity in horse muscle fractions was characterized by ATPase assay and compared to rabbit SR (**Figure 6*B***). ATP hydrolysis was measured using a spectrophotometric linked-enzyme assay, whereby production of ADP was coupled stoichiometrically to oxidation of NADH, as detected by a decrease in absorbance at 340 nm. ATPase activity was measured at 37 °C with saturating nucleotide concentration (5 mM Mg-ATP) in the presence and absence of 100 μM Ca_2+_, the difference being SERCA activity (Ca^2+^ - activated ATPase = total ATPase minus basal ATPase). The Ca^2+^ ionophore A23187 was added to prevent back-inhibition of SERCA by high luminal Ca_2+_ concentration. ATPase assays were performed in the presence of 5 mM azide, which inhibits the ATP synthetase running in reverse mode (proton pumping ATPase activity) in disrupted mitochondrial fragments present in muscle homogenates and fractions. Horse 10KP had a Ca^2+^-activated ATPase activity 4.0 ± 0.4 IU, as attributed to SERCA. Addition of the SERCA inhibitor thapsigargin (1 µM TG) eliminated Ca^2+^-activated ATPase activity in horse fractions H1, H2, 10KP, and 100KP, attributing the predominant contribution of SERCA to Ca^2+^-ATPase activity in these fractions. In comparison, rabbit SR had a Ca^2+^ activated ATPase activity of 7.6 ± 0.45 IU, as attributed to SERCA, using the same assay conditions (i.e., rabbit SR vesicles have ∼1.9-fold greater Ca^2+^-ATPase activity than the 10KP fraction of horse gluteal muscle, which is highly enriched in SR vesicles). The Ca^2+^-activated ATPase by the 100KP fraction from horse muscle reported here (1.4 ± 0.26 IU in **Figure 6*B***) is equal to or greater than the Ca^2+^-activated ATPase activity of horse SR preparations previously reported (**Table 1**), as identified by extensive online searches using PubMed and Google. Thus, the 10KP fraction of horse muscle exhibits high SERCA expression and high Ca^2+^-activated ATPase activity.

**Figure 6.**
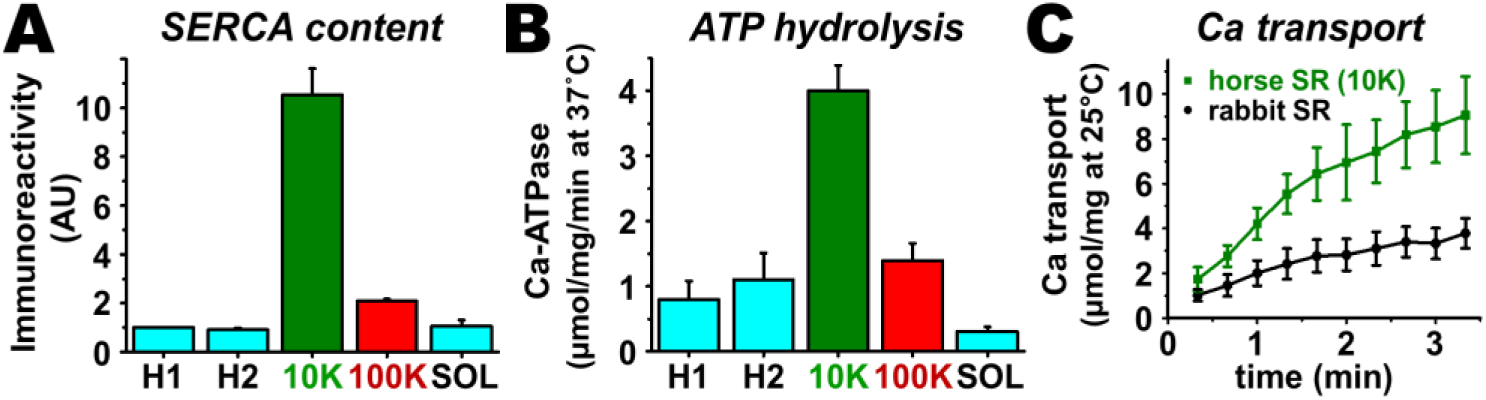
SERCA protein content, Ca^2+^-ATPase activity, and Ca^2+^ transport in horse muscle fractions. *A*, Immunoblotting was used to measure the relative SERCA content in horse muscle fractions and rabbit SR vesicles. The amount of SERCA protein in horse muscle fraction was calculated relative to the immunoreactivity of the H1 fraction, as determined per individual preparation with the initial H1 fraction set to an arbitrary unit (AU) of 1.0. Data are compiled from experiments performed as in Figure 5A, i.e., samples were assayed in a single-lane for relative comparison, with the per-sample protein loading amount adjusted to give a similar immunoreactive signal of SERCA/lane. *B*, Ca^2+^-activated ATPase activity was measured in horse muscle fractions and rabbit SR. The ATPase assay was run at 37 °C in the presence of the Ca^2+^ ionophore A23187, with saturating substrate concentration of Ca^2+^ and Mg-ATP (i.e., V_max_ assay). For comparison, the Ca^2+^-activated ATPase activity of rabbit SR vesicles is 7.6 ± 0.45 IU. *C*, ^45^Ca transport activity was measured for horse and rabbit SR vesicles. The transport assay was run at 25 °C in the presence of the Ca^2+^-precipitating anion oxalate, with saturating substrate concentration of Ca^2+^ and Mg-ATP (Vmax assay).

### Horse SR vesicles show enhanced Ca^2+^ transport compared to rabbit SR vesicles

ATP-dependent Ca^2+^ transport was measured using a radiometric filtration assay. ^45^Ca^2+^ transport was measured at 25 ⁰C under V_max_ condition, i.e., in the presence of saturating concentration of substrates: 3 mM ATP, 3.3 mM MgCl_2_, and an ionized Ca^2+^ concentration ([Ca^2+^]_i_) ∼ 2.4 µM (which was set using an EGTA/Ca^2+^ buffering system). Oxalate and azide were also added. Oxalate, a Ca^2+^-precipitating anion that diffuses into SR vesicles through anion channels, was added to the transport assay to remove back-inhibition of SERCA by high luminal Ca^2+^ concentration. Addition of oxalate allows Ca^2+^ transport by SERCA to proceed for minutes, thus providing a steady-state V_max_ measurement. Under these assay conditions, Ca^2+^ transport is specific to SR vesicles in muscle homogenate fractions because (i) Ca^2+^ uptake by mitochondria is inhibited by azide (119), and (ii) Ca^2+^ uptake by sarcolemmal vesicles is not facilitated by oxalate (120). The specificity of ATP-mediated Ca^2+^ transport was demonstrated by lack of ATP, which eliminated >95% of Ca^2+^ transport by horse SR vesicles. The specificity of SERCA-mediated transport was demonstrated by addition of TG, a SERCA-specific inhibitor (121) that eliminated >95% of Ca^2+^ transport by horse SR vesicles.

Using these assay conditions, the initial rate of Ca^2+^ transport by horse and rabbit SR vesicles was linear for ∼60–90 s (**Figure 6*C***). Ca^2+^ accumulation by horse and rabbit SR vesicles continued to increase for a total of ∼5 min until plateauing, when the measured rate of net ^45^Ca^2+^ uptake became zero. After that, the amount of accumulated Ca^2+^ decreased, e.g., 10‒15 min, due to bursting of SR vesicles from the growth of luminal Ca^2+^–oxalate deposits beyond vesicular capacity. Horse SR vesicles transported Ca^2+^ at a maximum rate of 4.2 ± 0.66 µmol Ca^2+^/mg protein/min (IU) at 25 °C, and rabbit SR vesicles transported Ca^2+^ at a maximum rate of 1.8 ± 0.51 IU (**Figure 6*C***). Thus, horse SR has a ∼2-fold greater rate of Ca^2+^ transport than rabbit SR, even though rabbit SR has ∼2-fold greater SERCA content and ∼2-fold greater Ca^2+^-ATPase activity (**Figure 6, Figure S4, Figure S5**). The high rate of Ca^2+^ transport by horse SR indicates that the membrane vesicles in the 10KP fraction, isolated by this protocol, are mostly intact and sealed (**Figure 4**). **Table 1** compiles published V_max_ rates of Ca^2+^ transport from previous reports of purification of SR vesicles from horse muscle (53–57). The Ca^2+^ transport rate of horse SR vesicles (**Figure 6*C***) is 8‒23-fold greater than the Ca^2+^ transport rates of horse SR vesicles reported by multiple laboratories (**Table 1**). Thus, this improved protocol for isolating SR vesicles from horse muscle provides a useful system for studying Ca^2+^ transport by SERCA1. We propose that the high V_max_ rate of Ca^2+^ transport by horse SERCA is facilitated by the lack of SLN protein and the abundance of CASQ protein in horse SR vesicles.

### Horse and rabbit SERCA show similar temperature-dependence of Ca^2+^-ATPase activity

The internal temperature of actively-contracting horse muscle is 42–44 °C (122). Since temperature controls the structure and activity of enzymes, we compared the temperature dependence of the Ca^2+^-activated ATPase activity for SERCA in horse and rabbit SR (**Figure S11**). At V_max_ conditions, with saturating substrate concentrations of Ca^2+^ and Mg-ATP, both horse and rabbit SERCA show a biphasic effect of enzyme activation and inactivation, with a robust ∼6-fold increase in activity when raising the temperature from 20 °C to 45 °C, followed by a steep ∼19-fold decrease in activity when raising the temperature from 45 °C to 50 °C (**Figure S11**). The 50% transition temperature from peak activity to thermal inactivation (T_i_) is ∼48 °C for both horse and rabbit SERCA (**Figure S11**). For comparison, rat SERCA in SR vesicles from rat skeletal muscle exhibits a 50% thermally-induced inactivation at T_i_ = 47 ± 0.7 °C (123). The similarities in thermal activation and inactivation of SERCA from horse and rabbit muscle indicate a similarity in protein structure and function of the two orthologs, such that global structure and temperature-dependence probably do not account for the increased relative rate of Ca^2+^ transport by SERCA in horse SR vesicles (**Figure 6*B, C***). The temperature-dependence assay also demonstrates that horse SERCA in the natural absence of horse SLN shows a similar biphasic thermal profile of Ca^2+^-ATPase activity as rabbit SERCA in the natural presence of rabbit SLN. We propose that the robust Ca^2+^ transport of SERCA in horse SR vesicles compared to rabbit SERCA is due, in part, to the lack of SLN inhibition and/or uncoupling of SERCA in horse SR, and possibly due to local differences in the amino acid sequence or horse SERCA.

### Horse and rabbit SERCA show similar Ca^2+^-dependent cleavage by Proteinase K

To assess the structure of horse SERCA, we utilized Proteinase K (ProtK), a protease which produces conformation-specific cleavage of rabbit SERCA at A domain residues in amino acid segments near the membrane boundary (124, 125). For example, in the presence of Ca^2+^ (E1•2Ca biochemical state), ProtK cuts rabbit SERCA selectively at residue T242 on stalk segment (SS3) leading out of the A domain into TM3, yielding a major C-terminal fragment comprising residues 243–994 (molecular mass of 83 kDa) (**Figure S12**). On the other hand, in the absence of Ca^2+^ (E2 biochemical state stabilized by TG), ProtK cuts rabbit SERCA selectively at residue L119 on stalk segment 2 (SS2) above the water/membrane interface of TM2 leading into the A domain, yielding a major C-terminal fragment comprising residues 120–994 (molecular mass of 95 kDa), with a low amount of secondary cleavage at T242, i.e., also producing a minor fragment of 83 kDa (**Figure S12**). As in the amino acid sequence of rabbit SERCA, horse SERCA1 also encodes residues L119 and T242 (**Figure 3**, **Figure S1**).

Here, ProtK digestion demonstrated that horse SERCA shows a similar Ca^2+^-dependent conformational change as rabbit SERCA, providing similar fragment intensities of the 83 kDa and 95 kDa bands as rabbit SERCA in the presence and absence of Ca^2+^, respectively (**Figure S12**). Thus, the ProtK assay demonstrates that horse SERCA in the absence of SLN shows a similar Ca^2+^-dependent conformation change as rabbit SERCA in the presence of SLN. We propose that (i) the structural dynamics of A domain stalk linkers S2 and S3 of horse SERCA is similar to that of S2 and S3 linkers rabbit SERCA, and (ii) the absence of SLN as SERCA-bound protein subunit does not affect the equilibrium of Ca^2+^ bound versus Ca^2+^ free states of horse SERCA in ligand-induced intermediates, i.e., non-cycling enzyme stabilized in E1•2Ca or E2•TG. We conclude that the ProtK assay demonstrates similarity in the structural dynamics of horse and rabbit SERCA, suggesting that the lack of SLN in horse SR helps enhance Ca^2+^ transport by horse SERCA.

As further validation of the reported protocol for purification of horse SR, the ProtK assay provides almost complete digestion of horse and rabbit SERCA (110 kDa bands) in 15 min at 23 °C (**Figure S12**), indicating that almost all of the SR vesicles in horse and rabbit preps are oriented right-side out, with an extravesicular location of the SERCA headpiece in SR vesicles from both species. We conclude that the reported protocol for isolation of SR vesicles from horse muscle (**Figure 4**, **Table 1**) provides a preparation suitable for functional and structural comparisons of horse SERCA to orthologs such as rabbit and human enzymes (13,50,81).

## Discussion

This study is a comparative assessment of gene expression and enzyme function of Ca^2+^ transport proteins from horse and rabbit muscle, thereby providing quantitative measurements of fundamental parameters of SR function at the molecular level. Our specific purpose was to analyze muscle Ca^2+^ transport regulation in horse, a species that has long-been selected for speed by evolution and breeding. Here we report quantitative analysis of RNA transcription of Ca^2+^ regulatory genes in horse muscle, *in silico* structural analysis of horse SERCA and SLN proteins, and biochemical analysis of Ca^2+^ transport in horse SR vesicles, with comparison to widely-utilized experimental and computational models from rabbit muscle. The generated results provide new insights on Ca^2+^ transport regulation in horse muscle, including proposed mechanisms for high muscular performance.

### Improved protocol for purification of SR vesicles from horse muscle

To date, there have been six reports of purification of SR vesicles from horse muscle with measurement of steady-state SERCA activity **(Table 1**). The most recent report of purification of horse SR vesicles allowed successful analysis of RYR and Ca^2+^-ATPase activities in muscle membranes from control horses and horses susceptible to recurrent exertional rhabdomyolysis; no differences were found in muscle membranes between the two sets (57). Here we further characterized subcellular fractions from horse muscle, finding that the 10,000 × *g* pellet (10KP fraction) has 6–50-fold greater SERCA activity than ‘SR vesicles’ previously reported (**Table 1**). The 10KP fraction isolated here expresses 10-fold higher SERCA content than the gluteal muscle homogenate, and this 10KP fraction is highly enriched in SR markers such as SERCA, CASQ, and phospholipids. Here the 10KP fraction (which we define as “SR vesicles”) of horse muscle has enabled detailed characterization of SERCA transport and ATPase activities, plus quantitation of the expression of Ca^2+^ regulatory proteins SLN, PLN, and CASQ. We also demonstrate that the traditionally-studied high-speed differential-centrifugation fractions from horse muscle, e.g., the 100,000 × *g* pellet (100KP) previously defined as “SR vesicles”, are less than optimal for examining the activity of horse SERCA, due to a low content of SERCA and a high content of GP and glycogen, thereby reducing the utility of functional and structural assays of Ca^2+^ transport regulation in these samples. Previous reports of SR preparations from horse muscle did not illustrate electrophoretic analysis of high-speed pellets using Coomassie blue, Stains-all, nor immunoblotting for SERCA, GP, regulatory peptides, or CASQ.

The protocol developed here, based in part on a key initial study (57), enables the selective determination of SERCA Ca^2+^ transport and Ca^2+^-ATPase activities in SR vesicles from horse muscle, while providing a similar ease of purification and high yield. With such low SERCA activity and probable high glycogen contamination in other reported SR preps, it is possible that the SR preparations isolated by these methods are a small subfraction of SR vesicles from horse muscle, and may not be representative of the SR system as a whole. Furthermore, SERCA activity is likely perturbed by high-glycogen contamination in these previously-reported preparations of horse SR vesicles; for example, addition of the non-ionic detergent C_12_E_8_ to 100KP increased Ca^2+^-activated ATPase activity of SERCA ∼2-fold, probably by dissociating aggregates of glycogen granules and SR vesicles. Thus, our new preparation of horse SR vesicles (10KP protocol) will be useful in assessing, with high fidelity, the protein level and enzyme activity of SERCA in SR vesicles, including the determination of the expression level and regulatory effect of peptide inhibitors in gluteal muscle from healthy and myopathic horses.

### Comparative analysis of SERCA activity in horse and rabbit SR vesicles

In this study, we compared SERCA activity from horse gluteus and rabbit skeletal muscles using an optical assay of ATP hydrolysis and a radiometric assay of Ca^2+^ transport. SERCA back-inhibition by luminal Ca^2+^ was relieved by the Ca^2+^ ionophore A2317 in the ATPase assay and the Ca^2+^-precipitating anion oxalate in the transport assay. The maximal Ca^2+^-activated ATPase activity of SERCA was lower in horse SR vesicles than in rabbit SR vesicles (measured per mg of SR protein), but the specific activity of horse SERCA was greater than rabbit SERCA (measured per relative content of SERCA/mg SR protein). The maximal ATP-dependent Ca^2+^ transport for horse SR vesicles was greater than that detected for rabbit SR vesicles (mg horse SR protein/mg rabbit SR protein), and the relative Ca^2+^ transport activity of horse SERCA was increased even further than rabbit SERCA (measured per relative content of SERCA/mg SR protein).

The differences in SERCA1 V_max_ could potentially be attributed to the lack of SLN peptide in horse. Another possibility for increased transport is sequence variation(s) in horse versus rabbit SERCA that affect the equilibrium distribution between biochemical intermediates along the enzymatic cycle. For example, single-residue variations R164H and R559C, each of which individually decrease the activity of bovine SERCA1, are sufficient to induce congenital pseudomyotonia syndrome in cattle (126–128). The lack of difference in the temperature dependence of Ca^2+^-ATPase activity by horse and rabbit SERCA does not support a transport effect by phospholipids: either directly through altered lipid composition and protein interaction, or indirectly through membrane fluidity and bilayer phase modulation.

Additional biochemical assays of SERCA structure and function included the ProtK cleavage assay and the thermal activation/inactivation enzyme assay, which provided additional information to help interpret results from standard Ca^2+^-ATPase and transport assays. The ProtK assay revealed a similar pattern of protein cleavage for horse and rabbit SERCA in Ca^2+^-bound and Ca^2+^-free biochemical states (**Figure S16**), indicating a similar residue accessibility and backbone dynamics of the two alternate ProtK sites in the A domain (residue T242 in E1•2C and residue L119 in E2•TG) for the two orthologous transporters. Horse and rabbit SERCA showed a similar activation of ATP hydrolysis upon increasing assay temperature from 20 °C to 45 °C using 5 °C steps (**Figure 15**), indicating that the two orthologous transporters have a similar standard free energy of enzyme activation (E_a_), which is a key parameter that determines enzyme rate. The similar profile of temperature-dependent enzyme activation is consistent with the similar rate of Ca^2+^-ATPase activity for the two SERCA orthologs measured at 37 °C, when normalized to the relative amount of SERCA content/total protein in SR vesicles (**Figure 6*A,B***). Thus, the intrinsic rate of Ca^2+^-dependent ATPase activity appear to be similar for horse and rabbit SERCA. Increasing the assay temperature from 45 °C to 50 °C inactivated both horse and rabbit SERCA almost completely, with each ortholog showing a similar 50% thermal inactivation at T_i_ ∼ 48 °C (**Figure 15**), suggesting that SERCA from the two species share a similar global structure and folding energetics. Although neither of these two biochemical assays provided information on molecular mechanism(s) that produce robust Ca^2+^ transport by horse SR, both assays demonstrated functional and structural similarities of SERCA orthologs from horse gluteus and rabbit muscle.

### Comparative sequence and structural analysis of horse SERCA luminal residues E894 and E959

Molecular modeling indicates that the global architecture of rabbit and horse SERCA are similar, as expected by high amino acid sequence identity (**Figure 3**). Comparing the sequences of horse and rabbit SERCA, there are 49 residue variations, plus a 1-residue deletion in horse SERCA (**Figure 3**, **Figure S1, Table S1**). Of the 49 variations in sequence between the two orthologs, 39 variations are conservative substitutions, while 9 variations are non-conservative substitutions (often a difference with His or Gly in either ortholog). One non-conservative amino acid substitution stands out for possible V_max_ control of horse SERCA: the negatively-charged residue E959 (**Figure S1**, **Figure S2*B***).

At the analogous positions, rabbit SERCA encodes K960 (L9–10) that forms a salt-bridge with E895 (TM8). The K960A mutation in rabbit SERCA does not qualitatively disrupt activity, yet the effect of this mutation on enzymatic parameters has not been reported (129); for example, is the V_max_ of rabbit SERCA increased when the E895–K960 salt-bridge is lost in the K960A mutant? At the analogous positions, chicken SERCA1 encodes T960 and E895 with the potential for hydrogen bonding. The T960A mutation in chicken SERCA reduces the stoichiometry of Ca^2+^ ions transported per ATP molecule hydrolyzed (92). At the analogous positions, cow SERCA encodes residues Q959 and E894 with the potential for hydrogen bonding. However, the x-ray crystal structure of the bovine ortholog shows different electrostatic interactions of luminal loops, which may be responsible for a lowered V_max_ (82). Thus, SERCA function depends on the identity of the amino acid at position 959/960 and its interactions with other luminal residue(s), which probably result in structural allosteric effects that control equilibria of biochemical intermediates and overall transport activity.

In horse SERCA, molecular modeling and MD simulation predict that there is not a salt-bridge between residues E894 and E959; instead, there is electrostatic repulsion (**Figure S2**). Since luminal loop residues are involved in the maintenance of transport/ATPase coupling in chicken SERCA1 and the decrease in V_max_ of bovine SERCA1, it is possible that the horse amino acid substitution E959 and the concomitant loss of TM8 interaction contribute to the enhanced V_max_ of horse SERCA. Furthermore, without this luminal salt-bridge in horse SERCA, these two negatively-charged carboxylate side chains in horse SERCA are available for alternative interactions. Luminal Ca^2+^ binding to SERCA in the E2•2Ca conformation has been well characterized in solution (124), and it has been proposed that SERCA provides significant luminal Ca^2+^ buffering in mammalian skeletal muscle myocytes (10). Since the concentration of SERCA in myocytes is ∼200 µM (as elegantly reviewed in (130)), it is likely that the side chains of E894 and E959 are luminally accessible with the potential to contribute up to 800 µM of liganding-oxygens for cation binding; as such, enhanced luminal Ca^2+^ binding by horse SERCA may act to increase the Ca^2+^ load stored in horse SR (**Figure 6*C***). Thus, we suggest that the E959 amino acid substitution contributes to the increased rate of Ca^2+^ uptake by horse SERCA and the increased level of Ca^2+^ load in horse SR.

### Comparative sequence analysis of SLN orthologs and potential mechanisms of SERCA regulation

In terms of steady-state protein expression (dependent on the rates of transcription, translation, and degradation, plus appropriate organelle targeting), it remains an outstanding question why such a supra-abundant level of *SLN* RNA is not translated into a stable SLN protein in horse gluteal muscle (compare *SLN* RNA transcription in **Figure 1** to SLN protein expression in **Figure 5** and **Figures S6–S9**). For rabbit SLN, the luminal C-terminus (^27^RSYQY^31^) and TM domain (residues ∼10–26) are important for targeting heterologous SLN to the endoplasmic reticulum of mammalian cells (COS-7 and HEK-293) grown in a dish (131, 132), serving as an experimental model to recapitulate protein targeting to SR in skeletal and cardiac myocytes in human tissues. In these cell culture models, rabbit SLN is correctly targeted to endoplasmic reticulum, except that cellular mal-targeting occurs upon deletion of the five C-terminal residues (Δ5 residues = rabSLN^1–26^) or upon addition of seven Leu residues to the C-terminal tail (rabSLN^1–31^-LLLLLLL^38^). An additional aspect of SLN sequence determinants of ER protein retention is co-expression of SERCA, whereby the mal-targeting of further-truncated C-terminal mutants of SLN (Δ6–9 residues) was rescued by heterologous over-expression of SERCA, with rabSLN^1-22^ (C-term Δ9) being correctly targeted and stably expressed with rabbit SERCA in the endoplasmic reticulum in HEK-293 cells. Since horse encodes a SLN ortholog (residues 1–29) of near-consensus length with a standard TM domain and only a single-residue deletion at the C-terminal tail (ΔY30), as expressed in muscle SR in the presence of a high level of endogenous SERCA (many-fold over than the level of SERCA protein in HEK-293 cells), we suggest that the lack of SLN peptide expression in horse gluteal muscle is <u>not</u> due to SLN mal-targeting or rapid degradation caused by the ΔY30 variation.

To examine structural mechanisms by which rabbit SLN inhibits rabbit SERCA, we previously utilized MD simulations, which predicted that (i) K_Ca_ inhibition is mediated by hydrogen-bond formation between N11-SLN and G801-SERCA, and (ii) transport/ATPase uncoupling is mediated by salt-bridge formation between E7-SLN and R324-SERCA, plus E2-SLN and K328-SERCA (67, 68). Compared to the rabbit system, horse SLN and horse SERCA encode the same residue pairs mediating these three inhibitory interactions: N10–G800, E2–K328, and E6–R324, per numbering of horse amino acid sequences (**Figure 2**, **Figure S1**) (40). We also used fluorescence resonance energy transfer (FRET) and site-directed mutagenesis to probe regulatory interactions of rabbit SLN and rabbit SERCA, finding that (i) SLN exists as multiple molecular species in membranes: SERCA-free (monomer, dimer, oligomer) and SERCA-bound (inhibitory dimer), and (ii) SLN residue I17 plays a dual role in determining the distribution of SLN homo-oligomers and stabilizing the formation of SERCA–SLN heterodimers (108). Compared to the rabbit system, horse SLN also encodes the analogous I16-SLN zipper residue that mediates these competing oligomeric interactions (**Figure 2**, **Figure S1**) (40). Thus, horse SLN and horse SERCA encode multiple, key residues that mediate inhibitory interactions.

To identify potential protein sequences responsible for horse exertional rhabdomyolysis, we previously evaluated sequences of four SERCA regulatory peptides from horses of three breeds (Thoroughbred, Standardbred, Quarter Horse) that developed recurrent exertional rhabdomyolysis compared to healthy horses of a similar breed (Przewalski’s Horse), plus two additional equine species (zebra and donkey) (40). SLN sequence was identical between horses susceptible to recurrent exertional rhabdomyolysis and controls. Here the amino acid sequence of SLN from Arabian horses was deduced (**Figure 2**), finding the identical 29-residue sequence as previously-characterized equine SLNs by Valberg *et al*. (40). For comparison, these horse SLN sequences were compared to an amino acid sequence alignment of 131 SLN orthologs, including three horse species, reported by Gaudry *et al*. (41). Five key differences are readily apparent. (i) All non-equine SLN orthologs encode ≥30 amino acids. (ii) Equine SLN are the only orthologs missing all proposed phosphorylation sites (S4, T4, T5) and the acylation site (C9) at the N-terminus. (iii) Equine SLN are the only orthologs with a single-residue deletion at the N-terminus, occuring at amino acid position 4 (Δ4). (iv) Equine SLN are the only orthologs that encode one Trp residue and two Arg residues at the N-terminus, suggesting a unique position (depth or tilt) in the membrane bilayer (SLN monomer), unique electrostatic control of self-association (SLN dimer and higher-order oligomers), and unique side chain interactions in the inhibitory groove of SERCA (SERCA‒SLN regulatory dimer). (v) Equine SLN are the only reported orthologs (^26^RSYQ^29^) that are missing the final residue (typically Y31) of the consensus C-terminal tail (^27^RSYQY^31^). The Y31 residue of consensus-length SLN may be accessible to the SR lumen for nitration, such as proximal nitration of SERCA luminal loop L3‒4 residues Y294 and Y295. Thus, we propose that horse SLN inhibition is not regulated in the short-term by post-translational modification. We also propose that the potency of SLN inhibition of SERCA in horse muscles and heart chambers depends on the relative expression level of SLN and SERCA proteins, which are likely regulated temporally through cellular mechanistic checkpoints such as transcription, translation, and degradation (133–135).

To advance these ideas, the amino acid sequence alignment of SLN by Gaudry *et al*. was used to assess additional structural/functional hypotheses (41). Via sequence analysis, two- and three-toed sloths 1ack the lack the Met^1^ start codon in *SLN*, thereby questioning the functional role of the SLN peptide in this clade (41), while also suggesting a possible non-coding role for *SLN* RNA. As noted, the grey short-tailed opossum #2 is the only reported species encoding a SLN peptide ortholog that does not encode an Asn residue at position 11 (i.e., position 10 in the numbering of equine SLN); instead, SLN from opossum #2 encodes S11, along with E2 and E7 (41). Here we predict, per MD simulation of rabbit SLN‒SERCA interactions by Espinoza-Fonseca *et al*. (67), that SLN from grey short-tailed opossum #2 does not inhibit the K_Ca_ of SERCA (due to the lack of N11 in SLN), but that SLN from grey short-tailed opossum #2 still acts to induce non-shivering thermogenesis by SERCA (due to the presence of E2 and E7 in SLN). Thus, comparative biochemical analysis of Ca^2+^ transport regulation in native SR systems may yield further insight on the mode(s) of SLN inhibition of SERCA.

### Positive and negative correlations of gene transcription and protein expression of SR Ca^2+^ transport regulators in horse gluteal muscle

Transcriptome sequencing is a powerful tool for identifying gene transcription and changes that occur, such as with disease. Previous work has shown that SLN protein expression correlates with SLN mRNA expression in mouse leg muscle (136). Here we examined the relationship between transcription and protein levels in horse muscle. mRNA transcript abundance is often an excellent proxy for protein expression; that is, for whether or not that protein is detectable within the cells (137, 138). To infer more exact cellular concentrations of proteins is imprecise because on average protein correlates with the abundances of their corresponding mRNAs with a squared Pearson correlation coefficient of 0.40 (139). Stronger correlations for Ca^2+^ regulatory and ATPase transcript abundance and protein expression, however, have been shown in a mouse microarray-proteomic study that included calsequestrin (CASQ2 r = 0.999893) and the sodium/potassium-transporting ATPase catalytic subunit (ATP1A1 r = 0.901647) (140). Vangheluwe *et al*. found that SLN mRNA expression in skeletal muscle correlates positively to SLN expression in Western blots in mouse, rat, rabbit, and pig skeletal muscle (26). Horses have exceptionally high *SLN* transcription relative to other regulatory peptides (*PLN*, *MRLN*, *DWORF*) and SERCA genes (*ATP2A1*, *ATP2A2*, *ATP2A3*). Thus, it was quite remarkable that our current immunoblotting results demonstrated that there is insignificant expression of SLN or PLN peptides in horse SR, as compared to SERCA (**Figure 5, Figure S4, Figure S5, Figure S6, Figure S10**). Horse SLN shows sequence divergence at the N-terminus and a sequence deletion at the C-terminus, which are regions that border the TM domain (**Figure 2**). To detect horse SLN in gluteal muscle fractions, we tested additional commercial and custom-made antibodies. The pAb ABT13, with immunogen the consensus C-terminus of SLN (^27^RSYQY^31^), showed little to no immunoreactivity for horse SLN. Immunoblot analysis using anti-SLN pAb ABT13 was successful in identifying SLN in rabbit SR (**Figure 5*C***). Our custom antibody GS3379 successfully detected synthetic horse SLN, yet found minimal level of SLN expressed in horse SR. By correlating the gene expression of *SLN* with the protein expression of SLN, we find that the high level of *SLN* RNA transcripts does not produce significant SLN protein. For comparison, rabbit skeletal muscle is reported to express 0.75 or 1.2 SLN per SERCA (mol/mol) (29, 102). Thus, we suggest that horse gluteus is the first demonstration of a native skeletal muscle that shows insignificant-to-undetectable expression of known regulatory peptides of SERCA in muscle. We acknowledge that additional research is needed to support this claim; for example, horse SR vesicles express a fast-migrating peptide on Coomassie-stained SDS-PAGE gel (**Figure S4**).

### Is there a known mammalian striated muscle that expresses SERCA in the absence of regulatory transmembrane peptide?

The long-held traditional view on Ca^2+^ regulation in striated muscles (skeletal and cardiac) has been that in each specific myocyte type, SERCA is co-expressed with one specific regulatory transmembrane peptide (either SLN or PLN) to produce a reversible inhibitory interaction, forming an heterodimeric complex whereby SERCA activity is inhibited until post-translational modification of the regulatory peptide submit (for an excellent review, see Primeau *et al*. (141)). More recently, there have been reports of three-way co-expression of SERCA, SLN, and PLN together, as assessed at the single-cell level, in human leg postural muscle (vastus lateralis) and Takotsubo cardiomyopathy patients (left ventricle), with enhanced SERCA inhibition (27, 142). These results support MacLennan’s early prediction of ‘super-inhibitory’ trimeric complexes comprised of one catalytic pump (SERCA) and two inhibitory subunits (SLN and PLN) expressed in native muscle and heart (71, 143), as assessed in cell culture models with epitope-tagged peptide regulators, which have received the newly appropriate family terminology as “regulins” (133, 144). As such, it is becoming clear that SERCA and its regulatory peptides are expressed broadly in tissue-specific patterns, with concomitant single and dual physiological coupling to SERCA (27,70,113,143).

In horse gluteal muscle, we made two observations that were surprising: (i) SLN and PLN proteins were detected at miniscule amounts in horse SR, i.e., SERCA is apparently unregulated by inhibitory transmembrane peptides in equine skeletal muscle, and (ii) the supra-abundant transcription of *SLN* gene RNA is disconnected from stable expression of SLN protein, which does not correlate with the greatly sub-stoichiometric level of SLN to SERCA protein in horse SR, i.e., horse gluteal muscle does not follow the central dogma of molecular biology: DNA to RNA to protein (**Figure 1*B***, **Figure 5*C***, **Figures S6–S10**).

Regarding surprising observation number one (i): the lack of significant levels of SLN and PLN in horse gluteal has generated novel questions, such as:

a. *Is there a reported mammalian skeletal or cardiac muscle that expresses SERCA in the absence of known regulatory transmembrane peptides?* We ran extensive literature searches looking for a case such as this, but we were unable to find a report identifying a mammalian striated muscle that expresses SERCA without a regulatory transmembrane peptide from the family known to regulate contractility, i.e., SLN, PLN, MRLN, sAnk1, or DWORF. Although we have extensive collective experience in the field, we have no knowledge of a mammalian striated muscle that expresses SERCA without one of these five regulatory transmembrane peptides.
b. *Does horse gluteal muscle express the three newly-discovered regulatory peptides from mouse muscle and heart:* MRLN, DWORF, or sAnk1? qRT-PCR and RNA-seq assays indicated that MRLN and DWORF genes are minimally transcribed in horse muscles (45-lower level than the SLN gene), so it is unlikely that MRLN and DWORF peptides are expressed in horse gluteus (Figure 1) (40). sAnk1 from mouse muscle acts a slight inhibitor of SERCA K_Ca_, producing about half the Ca^2+^ affinity inhibition as SLN or PLN (23, 113). sAnk1 enhances K_Ca_ inhibition of SLN or PLN when assembling as heterotrimers with SERCA. There is no commercial antibody available for horse sAnk1. sAnk1 is a key mediator stabilizing SR ultrastructure, yet considering the small inhibitory effect of sAnk1 on SERCA K_Ca_ (about half the effect of SLN or PLN) and the high activity of horse SERCA in gluteal SR, we suggest that it is unlikely that SERCA is primarily regulated by sAnk1 in horse muscle.
c. *Does horse gluteal muscle express an undiscovered regulatory peptide?* Unfractionated SR (10KP fraction) from horse muscle contains a low molecular mass Coomassie-staining band that migrates faster on SDS-PAGE than any band in unfractionated SR from rabbit muscle (Figure 5D, Figure S4). The identity of this low molecular mass peptide (< 10 kDa) in the horse 10KP fraction is unknown. This horse peptide was not detected by Coomassie-staining of the 100KP fraction, which contains a small amount of SERCA but relatively few other proteins, particularly with mass below 50 kDa (Figure 5D). This horse peptide was only partially-extracted from SR vesicles by the non-ionic detergent C_12_E_8_ (unlike SERCA, which was mostly solubilized), so it is unlikely that this low molecular mass horse peptide is derived from longitudinal SR and co-localized with SERCA. Wealso used multiple transcriptomic and biochemical assays in the search for regulatory transmembrane peptides of SERCA in our improved preparation of horse SR vesicles (10KP fraction), yet we did not identify any peptide regulator of SERCA in horse SR. We propose that horse gluteal muscle expresses SERCA without a transmembrane regulatory peptide, which accounts in part for the high rate of Ca^2+^-transport ATPase activity in horse SR vesicles. Per our comparative biochemical analysis of SERCA function in SR vesicles from horse and rabbit, we suggest that it may be informative to determine if SERCA is expressed without inhibitory transmembrane peptide in superfast striated muscles, such as rattlesnake shaker tail, bat sonar esophagus, hummingbird flight wing, and toadfish sonic swimbladder.

### Physiological implications of null-SLN and supra-CSQ expression on horse muscular performance

We propose that the low SLN/SERCA protein ratio in horse SR vesicles enhances the rate of SR Ca^2+^ reuptake and the loading level of SR Ca^2+^ stores (**Figure 5**, **Figure 6**, **Figures S4–S6**). Horse SR shows ∼2-fold greater Ca^2+^ transport than rabbit SR, in spite of a ∼2-fold lower SERCA content. These results are consistent with transgenic mouse models, where knock-out of SLN increases SERCA activity and over-expression of SLN decreases SERCA activity (35,36,145). The 2-fold greater Ca^2+^ load and higher CASQ expression in horse SR vesicles suggest that horse myocytes may be able to store and release more intracellular Ca^2+^ than rabbit myocytes. CASQ is the primary Ca^2+^ storage protein in JSR, and CASQ is tightly linked to the regulation of Ca^2+^ cycling in muscle. Since SRCa^2+^ release is sensitized by luminal Ca^2+^, an increase of luminal Ca^2+^ stores and associated RYR1 Ca^2+^ release could result in a more rapid and more powerful muscle contraction, i.e., producing a positive inotropic effect *in vivo*. Furthermore, the correlation of a higher rate of SERCA transport in SR vesicles with a more rapid rate of clearance of myosolic Ca^2+^ transient in myocytes could produce a faster rate of muscle relaxation, i.e., producing a positive lusitropic effect *in vivo*.

One reason why a lack of SERCA regulatory peptide inhibitor may have evolved in horses is to produce strong acceleration, enhanced by rapid muscular relaxation, giving horses an advantage in escaping predators. Horses have since been bred selectively to further enhance their speed and with that has come an incidence of exertional rhabdomyolysis ranging from 3 to 7% in horses (60, 146). Elevated myosolic Ca^2+^ has been detected in primary myocytes isolated from horse muscle during an episode of acute exertional rhabdomyolysis (147), and in cell culture myocytes differentiated from muscles from horses susceptible to recurrent exertional rhabdomyolysis, followed by exposure to increasing caffeine concentrations (148). Whereas slightly-elevated myosolic Ca^2+^ is an advantage that enhances muscle power output, excessively-elevated myosolic Ca^2+^ is a disadvantage that decreases contractility, and a heritable crossover between these two contractile conditions may occur following genetic pressure such as performance-based breeding(**Figure 7**).

**Figure 7.**
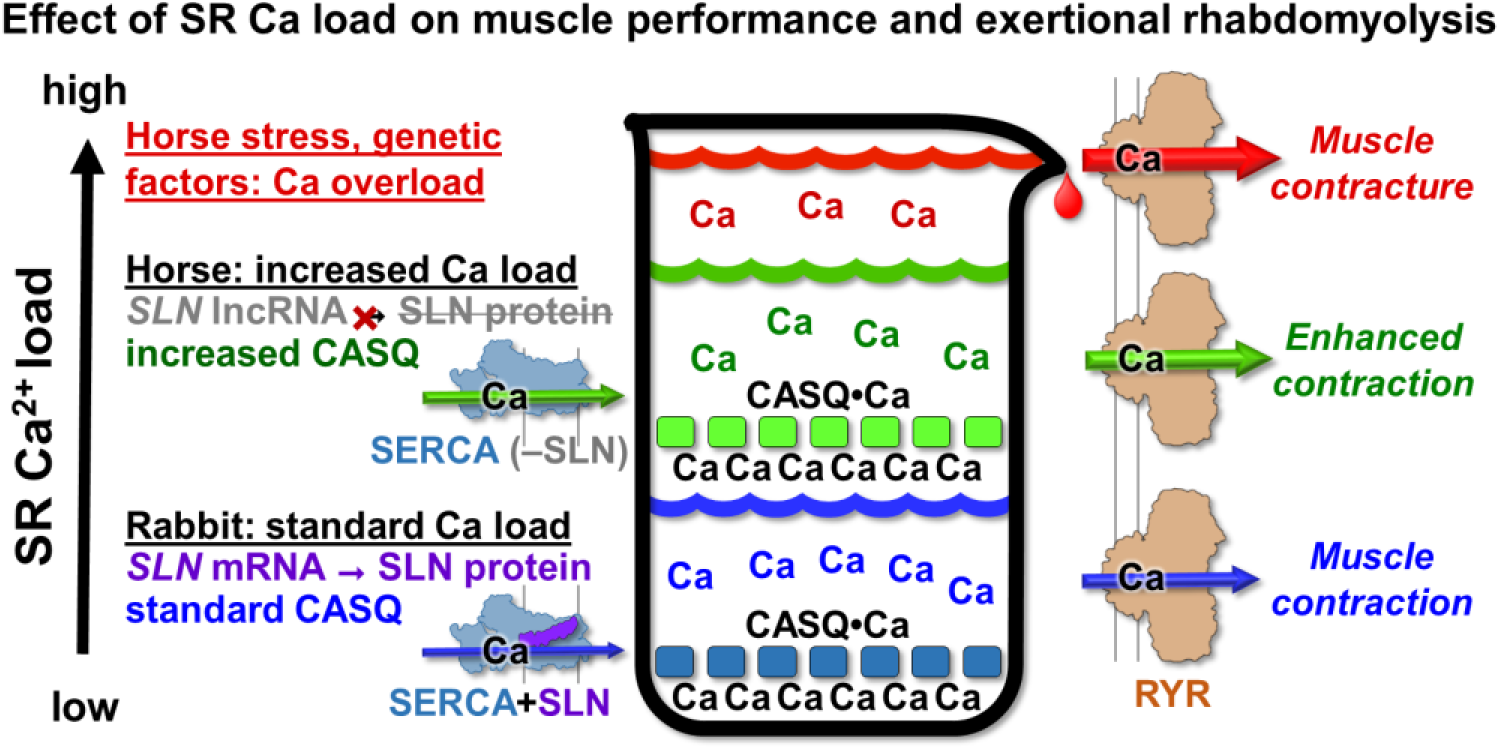
Proposed model for the role of SR calcium regulation in horse muscle performance and exertional rhabdomyolysis. This schematic diagram illustrates that lack of SERCA inhibition by SLN and increased level of CASQ expression enhance luminal Ca^2+^ stores in horse myocytes. High luminal Ca^2+^ promotes store-overload induced Ca^2+^ release (SOICR) through the RYR Ca^2+^ channel (7), which combined with stress-induced RYR Ca^2+^ leak, increases the incidence of muscle contracture events. Subsequent Ca^2+^-induced activation of proteases, lipases, and oxidative stress could contribute to the etiology of equine exertional rhabdomyolysis.

We propose that the low SLN/SERCA protein ratio in horse SR vesicles enhances the rate of SR Ca^2+^ reuptake and the loading level of SR Ca^2+^ stores, thereby enabling more rapid and more powerful contraction of horse gluteal muscle *in vivo*. An increase in the intrinsic rate of Ca^2+^ transport by horse SERCA (i.e., a possible lack of SLN uncoupling and/or an enhanced V_max_), coupled with a robust level of CASQ expression, may also serve as molecular mechanisms that potentiate SR contributions to functional contractility in horse muscle.

### Does the *SLN* gene transcript act as a functional non-coding RNA in horse gluteal muscle?

Transcriptional regulation is the primary mechanism for the control of eukaryotic protein expression, yet throttling of protein translation from mRNA is also common, including effects by encumbered 5’ noncoding sequences, regulation by micro-RNA’s, and inhibitory protein(s) binding to mRNA or ribosomes (149). At first, our unique detection of the dissociation between *SLN* RNA and SLN protein levels in horse muscle seemed surprising (**Figure 1*B***, **Figure 5*C***, **Figures S6–S9**). However, there is precedence in the literature: human airway smooth muscle cells express abundant *PLN* RNA but do not express PLN protein (150). Assays in this study included immunoblotting and immunoprecipitation of human airway smooth muscle homogenates, and immunohistochemistry of enzymatically-isolated human airway smooth muscle myocytes. These antibody-based approaches were corroborated by functional assay of cultured human ASM cells, whereby siRNA-PLN-knockdown had no effect on SR Ca^2+^ uptake in live cells or culture homogenates (150). Similarly, our functional assays indicated that SLN protein is minimally expressed in horse gluteus muscle (**Figure 1**), since the anti-horse-SLN pAb 3379 has no effect on Ca^2+^ transport by SR vesicles from horse gluteal muscle, consistent with the lack of SLN protein expression. These results indicate that the small level of SLN peptide expression detected in horse SR (∼1 SLN molecules per 13–16 SERCA molecules) has a minimal effect on the regulation of SERCA activity in gluteal myocytes *in vivo*.

Three recent articles report transcriptional, translational, and/or protein degradation mechanisms whereby lncRNA regulate SERCA. Tese newly-proposed molecular mechanisms may provide functional insights into the high level of *SLN* gene expression yet the low level of SLN peptide expression in horse muscle. First, the lncRNA *ZFAS* in mouse heart is proposed to be a dual-mode inhibitor of SERCA that acts by decreasing *ATP2A2* gene expression and by directly inhibiting SERCA2a enzyme activity (151). Thus, *ZFAS* lncRNA decreases the quantity and quality of Ca^2+^ transport in mouse SR. Analogously, we suggest the possibility that the high level of *SLN* RNA (sans SLN peptide) in horse gluteal muscle acts to regulate gene expression and/or activity of Ca^2+^ regulatory protein(s), including SERCA as a potential target. Second, hundreds of lncRNA with sORF (e.g., between 99 and 300 nucleotides) in human heart are reported to be translated as a peptide/small protein comprising 33 to 100 residues (also known colloquially as “micropeptide” or “microprotein”), including SR-targeted peptides that may act as regulators of cardiac contractility and development (152). Thus, lncRNA may switch between functional modes: non-coding, coding, or both. Third, the lncRNA *DACH1* is reported to directly bind and increase ubiquitination of SERCA in heart failure, thereby enhancing SERCA protein degradation, decreasing SR Ca^2+^ re-uptake, and inhibiting cardiac contractility (153). Thus, lncRNA control the steady-state level of cellular proteins by multiple mechanisms: transcriptional, translational, and/or degradation. These three studies indicate that lncRNA regulate a variety of central cellular functions through key molecular mechanisms, including Ca^2+^ regulation as a checkpoint hub (151–153). Indeed, there may be a species-specific array of currently-unidentified lncRNA that act in distinct roles to regulate myocyte contractility and muscle adaptation in health and disease.

### Proposed role of SLN expression in horse exertional rhabdomyolysis and relationship to human muscular dystrophies

With natural selection of horses as fast moving prey animals: why would they evolve to have a unique SLN amino acid sequence and lack of expression that are not present in closely-related ungulates such as rhinoceros? One possible advantage for the lack of a SERCA regulatory peptide inhibitor includes a genetic improvement that generates an increased luminal Ca^2+^ load to provide a greater ability to escape predators due to stronger acceleration and stronger contraction. Per breeding over millennia, could this enhanced Ca^2+^ cycling and contractile abilities have been selected to the extreme in some breeds, with *SLN* RNA upregulated and SLN protein downregulated? Thoroughbred horses with recurrent exertional rhabdomyolysis (RER) have 50% faster relaxation times than horses that are not predisposed to recurrent exertional rhabdomyolysis (154). Furthermore, horses with recurrent exertional rhabdomyolysis have faster racing speeds than those not predisposed to exertional rhabdomyolysis (59). Thus, two parameters of horse muscular performance are associated with horse susceptibility to RER.

In human muscular dystrophies, the effect of modulating SLN expression level and inhibitory function is unknown. In compelling mouse and canine models, enhancing SLN protein expression has been proposed as an effective therapy, as reported by Babu et al. (155). Indeed, SLN gene expression and protein levels are increased many-fold in standard mouse models of Duchenne’s muscular dystrophy, e.g., *mdx* and *mdx*/*utr*-*dko* (36). However, decreasing SLN expression in mouse models is reported to have beneficial or deleterious effects, depending on the genetic model and etiology studied (155–157). To help delineate these disparate effects, additional correlations should be determined among Ca^2+^ transport regulation and muscular performance in animal models and human disease, which may then provide insights on utilization of species-dependent mechanisms for contractile and therapeutic control.

### Remaining knowledge gaps and key directions for future horse SR research

The reported results are well supported and significant, but due to the breadth and novelty of our examination of Ca^2+^ regulation in horse muscle, there remain further questions. Below we discuss current limitations and future research priorities.

1. The number of horses utilized for SR isolation is small (N = 4); however, the amount of muscle utilized to obtain SR vesicles was large (∼200 g) and therefore necessitated euthanasia. However, we demonstrated the reproducibility of the protein expression profile of SR vesicles isolated from the gluteal muscle of four horses: three Quarter Horses and one Thoroughbred (Figure S5). Future research should investigate the effect of horse breed, sex, and age on SR vesicle purification, SERCA activity, and expression of regulatory peptides and CASQ. SR vesicles should also be isolated from horses with exertional rhabdomyolysis.
2. We assayed only one muscle: horse gluteus. Horse have 700+ muscles, which is greater than the number of human muscles. Among different species and muscle types, the SR network within myocytes is distinctly heterogeneous (3). SR vesicles from a myriad of horse striated muscles need to be tested, including alternative skeletal muscles (locomotive and postural) and the heart (four chambers), in order to better correlate gene expression of ATP2A and SLN, protein expression of SERCA and SLN, and Ca^2+^ transport/ATPase activities in horse SR vesicles.
3. Although possessing high SERCA activity, our unfractionated horse SR vesicles (10KP) comprise a mix of vesicle types, as demonstrated by the ∼2.2-fold lower SERCA content than unfractionated rabbit SR vesicles, and by the presence of Ca^2+^-independent ATPase activity (azide-sensitive) that is characteristic of contaminating mitochondrial vesicles (FoF1-ATPase running in reverse mode). For examples, the protein yield, protein composition, activity, and regulation of Ca^2+^ transport in SR vesicles purified by this reported protocol should allow further detailed biochemical assays of Ca^2+^ transport, Ca^2+^ storage, and Ca^2+^ release.
4. Multiple antibodies were used to assess SLN peptide expression in horse SR, finding little to no expression. However, it is possible that all antibodies used for immunoblotting are unable to recognize SLN peptide expression in horse muscle due to unknown reason(s). It is also possible that horse muscle expresses other known peptide inhibitor(s) of SERCA. Here we were unable to detect significant protein expression of SLN or PLN, or gene expression of MLRN or DWORF. Expression of ALN, ELN, or sAnk1 was not assessed. With the recently growing family of SERCA regulatory peptides, it is possible that horse muscle expresses an undiscovered peptide effector of SERCA that is an activator or inhibitor. The lowest molecular mass peptide in horse SR migrates faster than any Coomassie-staining band in rabbit SR (Figure 4D, Figure S4). The identity of the low molecular mass peptide in horse SR is unknown.
5. Two separate enzyme assays were used to assess the dual functions of SERCA under two different conditions. (i) Ca^2+^ transport assay at 25 °C with addition of oxalate to prevent luminal overload by Ca^2+^ precipitation inside the SR vesicles, thus allowing V_max_ determination by measuring the initial transport rate from 0–90 s. (ii) Ca^2+^-activated ATPase assay at 37 °C with addition of A23187 to prevent luminal overload by Ca^2+^ leakage from the SR vesicles, thus allowing V_max_ determination by measuring the initial Ca^2+^-ATPase rate from 30‒90 s. Using these two distinct conditions for steady-state enzyme assays of SR vesicles in vitro, the Ca^2+^/ATP coupling ratio of SERCA transport energetics cannot be calculated.
6. Activity assays of SR vesicles from horse and rabbit muscle measured V_max_ of SERCA, but not K_Ca_. There are many studies reporting variable effect of SLN regulation on SERCA K_Ca_ (inhibition or no effect) and SERCA V_max_ (activation, inhibition, or no effect). Here we focused on Ca^2+^ transport and Ca^2+^-ATPase assays measured at saturating substrate concentrations (Ca2+ and Mg-ATP), thus comparing horse and rabbit SERCA orthologs. Additional information on the enzymatic parameters is needed for horse SERCA in the absence of SLN, including determination of K_Ca_, Hill coefficient, and Ca/ATP coupling ratio, with comparison to rabbit SERCA in the presence of SLN.
7. Our transport assay detected Ca^2+^ uptake by SR vesicles, which is not a direct measure of SERCA activity. Instead, the 45Ca transport assay reports a combination of functional interdependence of SERCA, RYR, and CASQ, i.e., uptake, release, and storage of Ca^2+^, the three functions of SR. The goals are to improve the purity of horse SR vesicles by sucrose gradient subfractionation into longitudinal SR vesicles (with SERCA) and junctional SR vesicles (with RYR and CASQ). Future studies should investigate Ca^2+^ release from horse SR vesicles using endogenous RYR activators and inhibitors, and the effect of luminal store Ca^2+^ overload on RYR activity.
8. Additional SR proteins accessory to Ca^2+^ transport in muscle, such as junctin and triadin, were not assessed in horse SR vesicles. Other contributing proteins and regulatory control mechanisms in horse muscle may contribute to the robust Ca^2+^ transport by horse SR. There may also be unknown cellular effects due to compensatory molecular responses resulting from the lack of SLN peptide expression in horse gluteal myocytes. Useful information may result from future studies that investigate the proteomic profile and physiological control points of horse muscle SR.

### Summary

Our integrative methodologies (transcriptional, computational, and biochemical) provide novel information on SERCA activity in horse SR, which is a unique physiological system with high-capacity Ca^2+^ transport. Information reported here contribute to the understanding of Ca^2+^ cycling in myocytes, with relevance to muscular performance, adaptability, and disease. Additional functional studies are needed to dissect further the mechanistic roles of SLN, CASQ, and SERCA in horse muscle contractility, including correlations with gene and protein expression levels. We continue our experimental and computational analyses of horse Ca^2+^ transport in order to enhance the basic understanding and therapeutic utilization of Ca^2+^ control in muscle contractility.

## Experimental Procedures

### Materials

The Enzyme Commission (EC) number of SERCA is <u>7.2.2.10</u>^5^. Three enzymes were purchased from Sigma-Aldrich Corp.: pyruvate kinase (EC <u>2.7.1.40</u>^5^), L-lactate dehydrogenase (EC <u>1.1.1.27</u>^5^), and proteinase K (EC <u>3.4.21.64</u>^5^). ^45^Ca(CaCl_2_) was purchased from New England Nuclear Corp. The BCA Protein Assay Kit was purchased from Pierce Biotechnology, Inc. Laemmli-type SDS-PAGE gels were purchased from Bio-Rad Laboratories, Inc. Primary and secondary antibodies for immunoblotting were purchased from specific companies, as described below. Standard chemicals and Stain-all dye were purchased from Sigma-Aldrich Corp.

### Animals

The Research Animal Resources Facility at the University of Minnesota (U of MN) complies with the USDA Animal Welfare Act Regulations and the NIH Public Health Service Policy on Humane Care and Use of Laboratory Animals. U of MN has an approved Animal Welfare Assurance (A3456-01) on file with the NIH Office of Laboratory Animal Welfare. U of MN received accreditation renewal from the Association for Assessment and Accreditation of Laboratory Animal Care (AAALAC) in November 2015. All animal research was reviewed and approved by the Institutional Animal Care and Use Committee (IACUC) at U of MN, with IACUC protocol # 1511-33199A for horses and IACUC protocol # 1611-34327A for rabbits. Muscle samples were obtained from four horses (ages 10–18 yr) that were donated to U of MN for euthanasia due to orthopedic disease and were otherwise healthy: three castrated males (two Quarter Horses and one Thoroughbred) and one female (Quarter Horse). Horse owners provided written consent for obtaining samples for this research.

New Zealand white female rabbits (junior does less than six months of age) were provided by the Research Animal Resources Facility at U of MN. Standard protocol ensures that rabbits were housed for a minimal time. RAR staff performed euthanasia of rabbits with appropriate palliative care consistent with American Veterinary Medical Association guidelines. Rabbits were anesthetized prior to euthanasia to ensure minimal pain and distress; specifically, rabbits were given a subcutaneous injection of two sedatives: Acepromazine (1 mg/kg body weight) and Torbugesic (1 mg/kg body weight). After sedation, the rabbit’s ear was dabbed with wintergreen oil as a topical anesthetic, and a lethal dose of pentobarbital (100 mg/kg body weight) was injected through an ear vein catheter.

### Construction of a RNA transcriptome library from horse muscle

RNA-seq analysis was performed on snap-frozen gluteal muscle obtained by percutaneous needle biopsy from six healthy Arabian horses, as part of a previous study (64). Frozen muscle was ground to powder using a Biopulverizer (BioSpec Products, Inc.), and total RNA was extracted using TRIzol/chloroform, as previously described (158). To eliminate genomic DNA contamination, DNase treatment was performed on column (Direct-zol RNA MiniPrep Plus Kit, Zymo Research Corp.) using TURBO DNase (Thermo Fisher Scientific, Inc.). Quantification and quality of RNA, along with the degree of ribosomal RNA contamination, was assessed using the Pico chip on the Agilent Bioanalyzer 2100, and samples with a RNA integrity number (RIN) ≥ 7 were used for library preparation and quantification. The library was constructed using a strand-specific polyA^+^ capture protocol (TruSeq Stranded mRNA Kit, Illumina, Inc.).

### RNA sequencing, mapping, and assembly of the horse muscle transcriptome

Sequencing was performed using the Illumina HiSeq 2000 genome analyzer (100 base-pair paired-ends) at a targeted 35 million reads/sample. All paired-end RNA-seq reads were assessed for quality using the FastQC program (https://www.bioinformatics.babraham.ac.uk/projects/fastqc/). A total of 26,993 transcript sequences from the Ensembl EquCab 2.86 database (http://www.ensembl.org/Equus_caballus/Info/Index) were used to create the index file and map RNA-seq reads. Samples that passed the quality threshold (Q ≥ 30) were aligned to the horse transcriptome index using Salmon program 0.11.0 (159). Count data, generated as transcripts per million reads (TPM) by Salmon processing, were used to compile gene expression between each individual samples (N ≥ 5 horses per transcript) and for relative comparative analyses with rabbit gene expression. The previous RNA-seq datasets (64) that were analyzed during the current study (**Figure 1**) are available in the NCBI Gene Expression Omnibus (160) with GEO accession number GSE104388 (64).

### RNA-seq quantitation of gene expression in horse muscle

Gene expression of 7 targets (*ATP2A1*, *ATP2A2*, *ATP2A3*, *SLN*, *PLN*, *DWORF*, and *MRLN*) was extracted from RNA-seq data, normalized based on the length of each transcript and the overall sequencing depth per individual horse, and expressed as transcripts per million (TPM). The values for mean, range, and standard error of gene expression are reported as TPM (**Figure 1**).

### Deduced amino acid sequence of horse muscle proteins using genomic sequencing, genomic mining, and RNA sequencing

The NCBI genome browser was searched to identify the coding sequences of *ATP2A1, SLN*, *PLN*, *MRLN,* and *DWORF* from horse, rabbit, mouse, and human. Nucleotide and protein sequences for horse *ATP2A1* (SERCA1a) were determined from the horse reference genome EquCab2, Ensembl version 2.86. Our RNA-seq data (**Figure S1**) verified the coding sequence of horse *ATP2A1* (SERCA1a) reported in the horse reference genome EquCab2.86. The protein sequence of SERCA1a orthologs from rabbit, mouse, and human were obtained from the NCBI database with GenBank accession number <u>ABW96358.1</u>^6^ (Autry, *et al*., 2016), <u>NP_031530.2</u>^6^, and <u>NM_004320.4</u>^6^ (84), respectively. The protein sequence of SLN orthologs from rabbit, mouse, and human were respectively obtained from the NCBI database with Genbank accession number <u>NP_001075856.1</u>^6^ (73, 108) <u>AAH28496.1</u>^6^, and <u>AAB86981.1</u>^6^ (73). Protein sequences were aligned manually: SERCA1a orthologs in **Figure S1** and SLN orthologs in **Figure 2**.

### Molecular modeling of horse SERCA1

The x-ray crystal structure of rabbit SERCA1a (PDB ID <u>3W5A</u>^4^ (78) was used as the starting template for structural homology modeling of horse SERCA1a. The horse SERCA1a protein sequence (this work) and the rabbit SERCA1a protein sequence (GenBank accession number <u>ABW96358.1</u>^6^) were aligned manually, identifying 48 residue variations and a 1-residue deletion at position 504 in horse (**Figure S1**). The Modeller 9.17 software package (161, 162) was used to construct the homology model of horse SERCA1a (**Figure 3**), as previously utilized to construct molecular models of rabbit SERCA1a labelled with small-molecule fluorophore or GFP derivative (98, 163). During the modeling procedure, only the horse residues that were substituted from the rabbit amino acid sequence were allowed to be optimized, plus the two residues on each side of the deletion 504 in horse SERCA1a. The optimization and refinement protocol was set to ‘very-thorough’, as described in the Modeller 9.17 software documentation. A total of 50 molecular models of horse SERCA1a were generated and then evaluated using the DOPE score (164), and the top-scoring model (**Figure 3**) was used as a starting structure for further molecular simulation studies (next section).

### MD simulation of horse SERCA1

We used all-atom MD simulation to produce structures of horse SERCA1 embedded in a 1,2-dioleoyl-sn-glycero-3-phosphocholine (DOPC) phospholipid bilayer at simulated physiological conditions: 100 mM KCl, 3 mM MgCl_2_, and pH 7.0 (67). Three independent MD simulations of horse SERCA1 were run using NAMD (165), as previously described for rabbit SERCA1 (76,94,166,167). System set-up consisted of periodic boundary conditions (168), particle mesh Ewald for calculating electrostatic interactions in periodic molecular systems (169, 170), and a non-bonded cutoff of 0.9 nm, in order to use the RATTLE algorithm (171) to generate 200 ns trajectories of SERCA with a time-step of 2 fs. CHARMM36 force-field parameters and topologies were used for SERCA (172), DOPC (173), water molecule, potassium and chloride ions, and magnesium ion (174). The NPT ensemble was maintained with a Langevin thermostat (310 K) and an anisotropic Langevin piston barostat (1 atm). The solvated system was energy-minimized, heated to 310 K, and equilibrated for 20 ns. The equilibrated SERCA system was used to run three independent 200-ns MD simulations, producing a total simulation time of 600 ns.

### Isoelectric point calculation for SLN and SERCA orthologs

The online algorithm Isoelectric Point Calculator (IPC) was utilized (72). The average predicted pI is reported for orthologous SLN peptides (**Figure 2**), plus SERCA1 and CASQ1 proteins (**Results**).

### Purification of SR vesicles from horse muscle

Horses were humanely euthanized, and middle gluteal muscle was harvested, rinsed with ice-cold phosphate-buffered saline (PBS) at pH 7.4, and transported in chilled PBS from the veterinary clinic to the biochemistry laboratory within 1 h of euthanasia. Processing steps were performed in a 4 °C cold-room, with samples, tubes, and solutions on ice. Muscle tissue was trimmed of connective tissue, cut into 1 g chunks, and minced into small pieces. Minced muscle (200 g wet weight) was further disrupted using a 2-step Waring blender procedure: (i) blend 100 g tissue in 100 mL of 100 mM KCl, 20 mM MOPS (pH 7.0) for 15 s on high, and (ii) add 200 mL of 100 mM KCl, 20 mM MOPS (pH 7.0) and blend for 35 s on high. Next, the Waring blender procedure was repeated with another 100 g of tissue. The Waring-blended muscle homogenate samples were pooled, split into six SLA-1500 centrifuge bottles, and homogenized in each bottle using a Polytron homogenizer (Kinematica PT1300 with a 1.5-cm sawtooth generator) for 30 s on medium-high, yielding muscle fraction homogenate 1 (H1) (see horse SR protocol flow-chart in **Figure 4**). The H1 fraction was spun at 4,000 × *g_max_* for 20 min at 4 °C, yielding muscle fractions supernatant 1 (S1) and pellet 1 (4KP1). The 4KP1 fraction (6 pellets in 6 bottles) was re-homogenized as before (30 s on medium) in 300 mL solution of 100 mM KCl, 20 mM MOPS (pH 7.0) per bottle, yielding muscle homogenate 2 (H2). The H2 fraction was spun at 4,000 × *g_max_* for 20 min at 4 °C, yielding muscle fractions supernatant 2 (S2) and pellet 2 (4KP2). The S1 and S2 fractions were pooled, filtered through 5 layers of cheesecloth, and spun at 10,000 × *g_max_* for 20 min at 4 °C, yielding muscle fractions supernatant 3 (S3) and pellet 3 (10KP). The S3 fraction was spun at 100,000 × *g_max_* for 30 min at 4 °C, yielding muscle fractions supernatant 4 (SOL) and pellet 4 (100KP). The 10KP and 100KP pellets were resuspended in a solution of 250 mM sucrose, 20 mM MOPS (pH 7.0) using a Potter-Elvehjem glass-teflon homogenizer. Five horse muscle fractions were collected for biochemical analysis: H1, H2, 10KP, 100KP, and SOL (**Figure 4**). For storage, horse muscle fractions were aliquoted, snap-frozen in liquid nitrogen, and kept in a –80 °C freezer. The protein concentration of horse muscle fractions was determined using a colorometric BCA assay in microplate format with bovine serum albumin (BSA) as the standard; fractions were assayed in triplicate by n ≥ 2 assays.

### Purification of SR vesicles from rabbit muscle

SR vesicles were isolated from rabbit fast-twitch skeletal muscle using mechanical homogenization and differential centrifugation (50) (see also protocol flow-chart for rabbit SR purification in **Figure S3*A***). White, fast-twitch muscles from the back and hind legs were harvested (∼300–500 g), minced, and homogenized. A blender was used for homogenization (Waring Products, Inc.), with 100 g of tissue in 300 mL solution of 100 mM KCl and 50 mM MOPS, pH 7.0: 30 s at low-speed, 15 s off, and 20 s at low-speed. Rabbit muscle homogenate 1 (H1) was centrifuged at 4,000 × *g_max_*, yielding muscle fractions supernatant (S1) and pellet 1 (4KP1). The 4KP1 fraction was homogenized as before, yielding homogenate 2 (H2). The H2 fraction was centrifuged as before, yielding muscle fractions supernatant 2 (S2) and pellet 2 (4KP2). Supernatants 1 and 2 (S1 + S2) were pooled, filtered through cheesecloth, and centrifuged at 11,750 × *g_max_* for 30 min at 4 °C, yielding muscle fractions supernatant 3 (S3) and pellet 3 (12KP). To release myofibrillar and glycolytic proteins from rabbit SR vesicles in the S3 fraction, KC1 salt was slowly added to S3 to give a final concentration of 0.6 M, and then the S3 fraction was stirred at 4 °C for 15 min. The S3 fraction was then centrifuged at 23,400 × *g_max_* for 60 min at 4 °C, yielding muscle fractions supernatant 4 (SOL) and pellet 4 (P4 = unwashed rabbit SR vesicles). The P4 fraction was resuspended in a wash solution of 250 mM sucrose, 20 mM MOPS (pH 7.0) and centrifuged at 55,300 × *g_max_*, yielding pellet 5 (23KP = washed rabbit SR vesicles), which was resuspended in a solution of 250 mM sucrose and 20 mM MOPS (pH 7.0) using a Potter-Elvehjem glass-teflon homogenizer. Five rabbit muscle fractions were collected for biochemical analysis: H1, H2, P3 (8KP), P5 (23KP = washed SR vesicles), and SOL (S4 = 23K supernatant). For electrophoretic analysis of rabbit muscle fraction, see Coomassie and Stains-all gels in **Figure S4**. For storage, rabbit muscle fractions were aliquoted, snap-frozen in liquid nitrogen, and kept at –80 °C. The protein concentration of rabbit SR vesicles was determined using a colorimetric BCA assay in microplate format with bovine serum albumin (BSA) as the standard; fractions were assayed in triplicate by n ≥ 2 assays.

### Synthesis of SLN peptide orthologs

Horse SLN (29 residues), rabbit SLN (31 residues), and human SLN (31 residues) were synthesized by New England Peptides, Inc., using Fmoc solid-phase chemistry at 50-mg crude scale (16 µmol), including quality assessment by HPLC and mass spectrometry. SLN peptides were synthesized with an acetylated N-terminus and an amidated C-terminus. Synthetic SLN peptides were further purified in-house by HPLC (175). The concentration of SLN synthetic peptides was assessed by amino acid analysis, BCA assay, and densitometry of Coomassie blue-stained gels (31, 176).

### SDS-PAGE and gel-stain analysis

Muscle fractions were solubilized for 10 min at 23 °C in Laemmli sample buffer (69 mm Tris, pH 6.8, 11% glycerol, 1.1% lithium dodecyl sulfate, 144 mm β-mercaptoethanol, and 0.005% bromophenol blue), electrophoresed through Laemmli gels, and stained with the dye Coomassie blue R-250 or Stains-all dye. Coomassie-stained gels were imaged using a Bio-Rad ChemiDoc MP imaging system, and exposure time was optimized to prevent camera pixel saturation. Coomassie densitometry was performed using a Bio-Rad ChemiDoc MP imaging system. Stains-all was used for in-gel identification of CASQ (stained blue) and phospholipids (stained yellow) in a mix of total proteins (stained pink) (9,103,104). Stains-all gels were imaged using a hand-held color camera. In some cases, sample tubes were placed in a boiling water bath for 2 min prior to electrophoresis, in order to dissociate oligomers of PLB or SLN that are stable on Laemmli gels run at 23 °C.

### Quantitative immunoblotting

Immunoblotting was performed as previously described (83,177–179). Proteins were transferred to PVDF membrane (0.2-µm pore Immobilon-FL) in transfer solution of 25 mM Tris, 192 mM glycine, pH 8.3 Blots were blocked using Odyssey-TBS blocking buffer (LI-COR Biosciences, Inc.). Primary antibodies are described below. Primary antibodies were used typically at 1:1000 dilution, with incubation for 16–18 h at 4 °C. Secondary antibodies (fluorescently-labeled) were used at 1:15,000 dilution, with incubation for 20 min at 4 °C. After incubation with primary or secondary antibody, blots were washed three times with Tris-buffered saline (pH 7.4) with 0.05% Tween-20, and then once with Tris-buffered saline (pH 7.4). Sandwich-immunolabeling of the target protein with primary plus secondary antibodies was detected using an Odyssey laser scanner in near-infrared fluorescence mode, and bands were quantified using Image Studio Lite software (LI-COR Biosciences, Inc.).

### Anti-SERCA antibodies

Three commercial anti-SERCA antibodies were utilized. Immunoblot conditions and validation reports for these three commercial anti-SERCA antibodies are described as follows:

1. Anti-SERCA1 antibody VE121G9 is a mouse mAb (IgG1-kappa isotype) developed by Dr. Kevin P. Campbell (180). The immunogen was rabbit SR vesicles, and the epitope is located between residues 506‒994 of rabbit SERCA1a (180), a sequence which is 94% identical to horse SERCA1a (Figure S1). mAb VE121G9 was purchased as Ascites fluid diluted in PBS (Abcam, P.L.C.). Abcam reports that mAb VE121G9 reacts on immunoblot with rabbit, mouse, human, dog, pig, and amphibian SERCA1. mAb VE121G9 has been validated for SERCA1 immunoblot using human vastus lateralis homogenate (181), human diaphragm homogenate (182), mouse gastrocnemius homogenate (183). Here mAb VE121G9 was used for primary labeling at 1:1000 dilution and validated for horse SERCA1 immunoblot using horse gluteal homogenate (H1) and SR vesicles (100KP) (Figure 5C, Figure S5).
2. Anti-SERCA2 antibody 2A7-A1 is a mouse mAb (IgG2a isotype) developed by Dr. Larry R. Jones (115). mAb 2A7-A1 was purchased as Ascites fluid diluted in PBS (Abcam, P.L.C.). The epitope for mAb 2A7-A1 is residues 386‒396 of dog SERCA2a (184), a protein sequence (GenBank U94345.16) (83) that is identical with horse SERCA2a residues 386‒396 (EquCab3.0 ENSECAG000000076716) and rabbit SERCA2a residues 386‒396 (Genbank EU195061.16). mAb 2A7-A1 was used at 1:1000 dilution.
3. Anti-SERCA1/2/3 is a rabbit pAb (IgG) purchased from Santa Cruz Biotechnologies (H-300) as an affinity-purified pAb from whole serum. pAb H-300 recognizes SERCA isozymes 1, 2, and 3 from human, mouse, and rabbit orthologs, plus SERCA in horse muscle. The immunogen of this pan-isoform antibody is residues 1‒300 of human SERCA1a (Santa Cruz Biotechnologies), a sequence which is ∼95% identical in horse SERCA1a (Figure S1). Here pAb anti-SERCA1/2/3 was used for primary labeling at 1:1000 dilution (0.5 µg/mL). pAb H-300 has been discontinued by Abcam.

### Anti-glycogen-phosphorylase monoclonal antibody

Anti-GP ab88078 is a mouse mAb purified by protein-A chromatography (Abcam, P.L.C.). The immunogen was residues 734–843 of human muscle GP (UniProt reference sequence <u>P11217</u>^6^), which is 95% identical to horse muscle GP (GenBank <u>NP_001138725.1</u>^6^) and 96% identical to rabbit GP (GenBank <u>NP_001075653.1</u>^6^). Abcam reported that mAb ab88078 reacts with human and mouse muscle GP on immunoblot using mAb ab88078 at a concentration of 1 µg/mL. mAb ab88078 has been validated on GP immunoblot using human muscle homogenate (185), mouse muscle membrane vesicles (186), rat brain homogenate (187), and rat muscle homogenate (188). Here mAb ab88078 was used for primary labeling at 1:1000 dilution (0.43 µg/mL), with successful detection of horse GP (**Figure 5*B***). mAb ab88078 has been discontinued by Abcam.

### Custom anti-horse-SLN polyclonal antibody (pAb GS3379)

Due to sequence divergence of horse SLN at the N- and C-termini compared to rabbit, mouse, and human orthologs (**Figure 2**), we designed and purchased an anti-horse-SLN polyclonal antibody from Genscript (pAb GS3379). The immunogen was a 6-residue N-terminal peptide of horse SLN (Q^1^MEWRRE^6^C), with an added ^−1^Gln residue (to mimic an acetylated N-terminus) and an added ^+7^Cys residue (to serve as a C-terminal linker). The horse N-terminal SLN peptide was conjugated to keyhole limpet hemocyanin (KLH) as the carrier protein for rabbit immunization. The anti-horse-SLN pAb GS3379 was purified from rabbit serum using affinity purification on the same horse-SLN-peptide coupled as an affinity-column ligand. Antibody generation, affinity purification, and titer determination were performed by Genscript. Here pAb GS3379 was used for primary labeling at 1:1000 dilution (0.4 µg/mL). pAb GS3379 was validated using synthetic horse SLN as a quantitative immunoblot standard (**Figure 5*C***, **Figure S6**).

### Custom C-terminal anti-rabbit/mouse/human SLN polyclonal antibody (PFD1-1) from Dr. Patrick Desmond and Prof. Robert Bloch

Dr. Patrick F. Desmond and Professor Robert J. Bloch, from the University of Maryland School of Medicine, provided a commercially-generated anti-SLN pAb (113). Two immunogens were used for rabbit immunization: a base 7-residue C-terminal peptide conserved in rabbit/mouse/human SLN with two alternate variations in the SLN peptide at both the N- and C-terminal positions (acetyl-^25^LVRSYQY^31^-amide and C-aminohexanoic acid-^25^LVRSYQY^31^-OH). Both SLN peptides were conjugated to bovine serum albumin (BSA). Antibody generation and affinity purification were performed by 21st Century Biochemicals, Inc. The anti-rabbit/mouse/human-SLN pAb PFD1-1 was validated on immunoblot using mouse muscle homogenate and recombinant mouse SLN (113). Horse SLN encodes ^24^LVRSYQ^29^ (i.e., horse SLN is missing one residue of the seven-residue immunogen: the terminal Tyr residue of the pAb PFD1-1 epitope). Here pAb PFD1-1 was used for primary labeling at 1:1000 dilution. Although horse SLN encodes six of seven residues in the immunogen peptide sequence, horse SLN shows significantly lower labeling than rabbit SLN by pAb PFD1-1 (**Figure S7**).

### Commercial anti-SLN polyclonal antibodies

Five anti-SLN antibodies were purchased. One commercial anti-SLN antibody (ABT13) was raised against the mostly-conserved C-terminus of SLN. Three commercial anti-SLN antibodies (pAb 18395-1-AP, pAb N-15, and pAb bs-9855R) were raised against human SLN (N-terminal peptide or whole protein). One commercial anti-SLN antibody (pAb E-15) was raised against mouse SLN (N-terminal peptide). Immunoblot conditions and validation reports for commercial anti-SLN antibodies are described as follows:

1. Anti-SLN ABT13 is an affinity-purified rabbit pAb (Millipore-Sigma). The immunogen was C-terminal residues ^26^VRSYQY^31^ of rabbit/mouse/human SLN as a sulfolink-peptide (CGG-^26^VRSYQY^31^) conjugated to KLH. Horse SLN encodes 26VRSYQ30 (i.e., missing the terminal Y31 of the rabbit/mouse/human SLN immunogen). Millipore-Sigma reports that ABT13 reacts with rabbit, mouse, and human SLN on immunoblot at 1.5 µg/mL pAb, as corroborated for rabbit SLN using 0.5 µg/mL pAb ABT13 (1:1000 dilution) and fluorescent secondary antibody detection (Figure S8). pAb ABT13 has been validated for SLN immunoblotting using human right atrium homogenate (189) and mouse soleus and diaphragm homogenates (190). Here pAb ABT13 was used for primary labeling at 1:1000 dilution, with positive detection of SLN in rabbit SR (Figure S7), plus synthetic rabbit and human SLN.
2. Anti-human-SLN 18395-1-AP is an affinity-purified rabbit pAb (Proteintech Group, Inc.). The immunogen was human SLN (residues 1–31) conjugated as a fusion protein to glutathione S-transferase (GST). Compared to human SLN protein, horse SLN is 77% identical (5 sequence variations and 2 deletions), and rabbit SLN is 81% identical (6 sequence variations) (Figure 2). Proteintech Group reports that pAb 18395-1-AP reacts with SLN in human and mouse heart homogenates on immunoblot using antibody concertation of 1.6 µg/mL and 0.5 µg/mL pAb, respectively. pAb 18395-1-AP has been validated on SLN immunoblot using mouse heart homogenate (191). Here pAb 18395-1-AP was used for primary labeling at 1:1000 dilution, a condition where pAb 18395-1-AP did not detect rabbit or horse SLN but showed non-specific binding to other proteins in rabbit and horse SR (Figure S9).
3. Anti-human-SLN bs-9855R is a rabbit pAb IgG purified by protein A chromatography (Bioss, Inc.). The immunogen was synthetic human SLN (residues 1–31) conjugated as a fusion protein to KLH. Compared to human SLN protein, horse SLN is 77% identical (5 sequence variations and 2 deletions), and rabbit SLN is 81% identical (6 sequence variations) (Figure 2). Bioss reports that bs-9855R reacts with human, mouse, and rat SLN in immunohistochemistry assay. Here pAb bs-9855R was used for primary labeling at 1:1000 dilution.
4. Anti-human-SLN N-15 is an affinity-purified goat pAb (Santa Cruz Biotechnology, Inc.). The immunogen was a N-terminal peptide of human SLN (proprietary peptide sequence). Santa Cruz Biotechnology reported that pAb N-15 reacts with human and dog SLN on immunoblot using a pAb concentration of 1 µg/mL. Here pAb N-15 was used for primary labeling at 1:500 dilution (0.5 µg/mL). pAb N-15 did not detect rabbit SLN by immunoblot using rabbit SR vesicles (2 µg protein per lane). pAb N-15 has been discontinued by Santa Cruz Biotechnology.
5. Anti-mouse-SLN E-15 is an affinity-purified goat pAb (Santa Cruz Biotechnology, Inc.). The immunogen was a N-terminal peptide of mouse SLN (proprietary peptide sequence). Santa Cruz Biotechnology, Inc. reported that pAb E-15 reacts with mouse and rat SLN on immunoblot using a pAb concentration of 1 µg/mL. Here pAb E-15 was used for primary labeling at 1:500 dilution (0.5 µg/mL). pAb E-15 did not detect rabbit SLN by immunoblot using rabbit SR vesicles (2 µg total protein per lane). pAb E-15 has been discontinued by Santa Cruz Biotechnology.

### anti-PLN monoclonal antibody 2D12

Anti-PLN antibody 2D12 is a mouse mAb (IgG2a isotype) generated by Dr. Larry R. Jones (114). The immunogen of mAb 2D12 was residues 2–25 of dog PLN conjugated to thyroglobulin using glutaraldehyde. The epitope for mAb 2D12 is residues ^7^LTRSAIR^13^ of dog PLN (184), a sequence which is identical for horse and rabbit PLN (40). mAb 2D12 (ab2685) was purchased from Abcam, P.L.C., as an Ascites fluid stock solution diluted in PBS to 1 mg/mL. Abcam reports that mAb 2D12 reacts on immunoblot with PLN from rabbit, mouse, human, pig, sheep, chicken, guinea pig, and dog. mAb 2D12 has been validated for quantitative immunoblotting using multiple PLN orthologs: native and recombinant dog PLN, native mouse PLN, native human PLN, synthetic human PLN, and synthetic pig PLN (83,114,129,177–179,192,193). Here mAb 2D12 was used for primary labeling at 1:1000 dilution, with successful detection of synthetic human PLN (high level) and horse PLN in SR vesicles (low level) purified from horse gluteal muscle (**Figure S11**). In additional control experiments, mAb 2D12 detected PLN protein in SR vesicles purified from horse left ventricle, horse right atrium, and pig left ventricle.

### Secondary immunolabeling with fluorescent anti-IgG antibodies

Secondary antibodies labeled with near-infrared fluorophore (700 nm or 800 nm emission) were purchased from LI-COR Biosciences, including affinity-purified goat antisera against mouse IgG (goat anti-mouse pAb) and affinity-purified goat antisera against rabbit IgG (goat anti-rabbit pAb). Secondary antibodies were used at 1:15,000 dilution (20 min at 4 °C) for sandwich-labeling of target protein through primary antibody binding. Fluorescent bands were detected using an Odyssey laser scanner in near-infrared mode and quantified using Image Studio Lite software (LI-COR Biosciences, Inc.).

### Ionophore-facilitated Ca^2+^-activated ATPase assay

ATP hydrolysis was determined using a spectrophotometric assay of phosphate release. The assay was started in a 2 mL solution of 100 mM KCl, 0.1 mM EGTA, 5 mM MgCl_2_, 5 mM Na_2_ATP, 5 mM NaN_3_, and 50 mM MOPS (pH 7.0) at 37 °C. The Ca^2+^ ionophore A23187 was added (3 μg/mL = 5.7 µM) to eliminate the accumulation of Ca^2+^ inside SR vesicles, i.e., to relieve product inhibition (119). ADP production was coupled to NADH oxidation by a linked-enzyme ATP-regenerating system: 10 U/mL pyruvate kinase, 0.6 mM phosphoenol pyruvate, 10 U/mL lactate dehydrogenase, and 0.2 mM NADH (112). Assays were started with the addition of muscle fractions (5–20 µg protein/mL). After 1-min equilibration, Ca^2+^-independent ATPase activity was measured for 2‒3 min, and then 100 µM CaCl_2_ was added (20 µL of 100 mM stock) to activate Ca^2+^-dependent ATP hydrolysis by SERCA. The rate of ATP hydrolysis was calculated as the rate of NADH oxidation by measuring the decrease in NADH absorbance at 340 nm using an extinction coefficient of 6220 M^−1^ cm^−1^, as detected by a Spectramax 384 spectrophotometer (Molecular Dynamics, Inc.) in cuvette mode. SERCA activity is defined as the Ca^2+^-dependent ATPase activity (i.e., the difference in ATP hydrolysis rate measured in the presence and absence of Ca^2+^) and expressed in international units (1 IU = 1 μmol product/mg protein/min) (194).

### Oxalate-facilitated 45Ca2+ transport assay

Ca^2+^ transport was measured using a filtration assay with ^45^Ca radioactive tracer (83, 195). Assays were conducted at 25 °C with 10 μg of SR protein in 1 mL solution containing 100 mM KCl, 3.3 mM MgCl_2_, 3.0 mM Na_2_ATP, 10 mm K_2_-oxalate, 5 mM NaN_3_, and 50 mM MOPS (pH 7.0), with 2 mm EGTA and 1.8 mm CaCl_2_ (containing trace amounts of ^45^Ca) to give an ionized Ca^2+^ concentration ([Ca^2+^]_i_) of 2.4 µM, i.e., a V_max_ assay with saturating concentration of the substrates Ca^2+^ and Mg-ATP. Transport assays were started with the addition of protein samples to reaction tubes. Ca^2+^ transport was terminated at serial time intervals by vacuum filtering 100 µL of assay solution (1 µg of SR protein) through a glass-fiber filter (Millipore HA, with 0.45 µm pore size), which was washed twice with 5 mL of ice-cold 150 mM NaCl solution. The loading of ^45^Ca inside SR vesicles was determined by liquid scintillation counting. Background ^45^Ca binding (defined as a Millipore filter blank with 1 µg SR protein in the absence of ATP) was subtracted from experimental values (the Millipore filter sample with 1 µg SR protein in the presence of ATP) to yield ATP-dependent Ca^2+^ transport by muscle fractions.

### Proteinase K assay of SERCA conformational state

Limited proteolysis by ProtK was used to assess the ligand-dependent conformational state of horse and rabbit SERCA (124, 125) (**Figure S12**). The standard assay solution contained 50 mM NaCl, 0.5 mM MgCl_2_, and 20 mM MOPS (pH 7.0), with addition of either 0.1 mM CaCl_2_ (E1•2Ca biochemical state) or 2 mM EGTA and 1 μM TG (Ca^2+^-free E2•TG biochemical state) (98). SR vesicles at 500 µg/mL were hydrolyzed with 12.5 µg/mL ProtK (40:1 wt:wt protein) for 15 min at 23 °C. Proteolysis was stopped by the addition of ice-cold trichloroacetic acid (TCA at 2.5% w:v), followed by addition of Laemmli sample buffer (final concentration of 1.1% lithium dodecyl sulfate). SR samples were electrophoresed through a 4–15% Laemmli gel, and stained with Coomassie blue. Protein bands were imaged and quantified using a Bio-Rad ChemiDoc MP imaging system.

### Experimental design, statistical analysis, and data presentation

Biochemical assays were performed using independent SR vesicle preparations from N = 3–4 horses and N = 3–6 rabbits. Data are reported as mean ± standard error (SEM), which were calculated typically from *n* ≥ 2 independent experiments for each SR prep. Experimental design was not randomized, and scientists were not blinded to sample identity during data acquisition or analysis. Data graphs were generated using Origin 9.2 (OriginLab Corporation; Northampton, MA).

## Supporting information

Supplement: 1 Table, 12 Figures, 22 References

## Acknowledgments

We thank Kevin L. Campbell (University of Manitoba) for insightful discussions and manuscript input. We thank Seth Robia, Howard Young, and Naresh Bal for insightful discussions. We thank Samantha Yuen, Morgan Zander, and Ty Gaedtke for technical support. We thank Jonathan Marchant for equipment access. We thank Octavian Cornea, Destiny Ziebol, and Sarah Blakely-Anderson for administrative support. We thank Patrick Desmond and Robert Bloch (University of Maryland) for providing the custom-designed anti-mouse/rabbit/human-SLN polyclonal antibody PFD1-1. RNA sequencing was performed by the University of Minnesota Genomics Center. Computational resources were provided by the University of Minnesota Supercomputing Institute. Spectrophotometric assays were performed in the Biophysical Technology Center, University of Minnesota Department of Biochemistry, Molecular Biology, and Biophysics.

## Contributions

S.J.V.: study design, project administration, transcriptomics, genomics, protein sequence/structure analysis, sample collection, SR purification, biochemical assays, manuscript writing, grant funding, and expert team leadership.

J.M.A.: study design, project administration, protein sequence/structure analysis, sample collection, SR purification, biochemical assays, SLN antibody design, immunoblotting, fluorescence spectroscopy, manuscript writing, manuscript figures, and grant funding.

C.B.K.: quantitative immunoblotting and peptide purification.

B.S.: molecular modeling, MD simulation, protein sequence/structure analysis, manuscript writing, and manuscript figures.

S.F.C.: biochemical assays, fluorescence spectroscopy, and protein sequence analysis. M.C.: biochemical assays.

S.P.: transcriptomics, genomics, and protein sequence analysis. Z.C.: biochemical assays and grant funding.

L.M.E.-F.: MD simulation, molecular modeling, protein sequence/structure analysis, and grant funding.

C.J.F.: transcriptomics.

D.D.T.: study design, project administration, manuscript writing, and grant funding.

All authors contributed research ideas, reviewed results, and approved manuscript submission.

## Funding Sources and Disclaimers

This study was supported in part by a Morris Animal Foundation grant to S.J.V, J.M.A., and D.D.T. (MAF D16EQ-004). Morris Animal Foundation is the global leader in supporting science that advances animal health. This study was supported in part by National Institutes of Health grants to D.D.T. (R01 GM27906, R01 HL129814, and R37 AG26160) and L.M.E.-F. (R01 GM120142). The content is solely the responsibility of the authors and does not necessarily represent the official views of the National Institutes of Health. This study was supported in part by an American Heart Association grant to Z.C. (#18TPA34170284/Chen,Zhenhui/2018). The authors declare that they have no conflicts of interests with the contents of this article.

## Footnotes

^1^Abbreviations and acronyms used are: *ATP2A1*, gene encoding SERCA1 protein isoforms; *ATP2A2*, gene encoding SERCA2 protein isoforms; *ATP2A3*, gene encoding SERCA3 protein isoforms; Ca^2+^, calcium; CASQ, calsequestrin; codRNA, protein-coding RNA; DWORF, dwarf open reading-frame peptide; *g*, gravitational force equivalent of 9.8 m/s^2^ standardized to 1 *g* at average Earth surface; GP, glycogen phosphorylase; IU, international unit of enzyme activity, defined as the production of 1 µmol product per milligram protein per minute; K_Ca_, apparent Ca^2+^ dissociation constant, defined as the Ca^2+^ concentration required for half-maximal activation of SERCA activity; KLH, keyhole limpet hemocyanin; lncRNA, long non-coding RNA; mAb, monoclonal antibody; MD, molecular dynamics; MRLN, myoregulin; mRNA, message RNA; pAb, polyclonal antibody; pI, protein isoelectric point defined as the pH at which the net protein charge is zero; PLN, phospholamban; ProtK, proteinase K; qRT-PCR, mRNA quantitation using real-time reverse transcription polymerase chain reaction; RER, recurrent exertional rhabdomyolysis; RNA-seq, whole transcriptome shotgun sequencing; RYR, ryanodine receptor Ca^2+^ release channel; SEM, standard error of the mean; SERCA, sarco/endoplasmic reticulum Ca^2+^-transporting ATPase; SLN, sarcolipin; sORF, small open reading frame; SOICR, store overload-induced Ca^2+^ release; SR, sarcoplasmic reticulum; TG, thapsigargin; T_i_, the temperature at which 50% transition occurs between peak enzyme activity and thermal inactivation; TM, transmembrane helix; TPM, transcripts per million reads; V_max_, maximal enzyme velocity of SERCA measured at saturating concentration of ionized Ca^2+^ (1–10 µM) and Mg-ATP (1–10 mM).

^2^RNA-seq data from Arabian horse gluteal muscle were deposited in the Sequence Read Archive (SRA) database with accession number <u>SRP082284</u>. Gene expression values from rabbit muscle were mined from RNA-seq data deposited with SRA accession number <u>SAMN00013649</u>.

^3^This article contains **Table S1** and **Figures S1–S12** as **Supporting Information**.

^4^Research Collaboratory for Structural Bioinformatics Protein Databank. (1) Rabbit SERCA1a: PDB # <u>3W5A</u>, <u>1IWO</u>, <u>2ZBD</u>, and <u>1SU4</u>. (2) Cow SERCA1a: PDB # <u>3TLM</u>. The molecule or structure for an accession ID can be identified through the NCBI Entrez utility at http://www.ncbi.nlm.nih.gov/sites/gquery.

^5^Enzyme Commission number for IUBMB Enzyme Database. SERCA Ca^2+^-transporting ATPase = <u>7.2.2.10</u>. Glycogen phosphorylase <u>2.4.1.1</u>. Pyruvate kinase = <u>2.7.1.40</u>. L-lactate dehydrogenase = <u>1.1.1.27</u>. Proteinase K (i.e., peptidase K) = <u>3.4.21.64</u>. The enzyme for an EC number can be identified through the ExplorEnz utility at http://www.enzyme-database.org/about.php.

^6^GenBank accession code for cDNA sequences of seven target proteins. (1) SERCA1a: horse <u>XM_001502262.6</u>, rabbit <u>ABW96358.1</u> (Autry, Winters et al. 2008), mouse <u>BC036292.1</u> (196), and human <u>NM_004320.4</u> (84). (2) SERCA2a: horse <u>XM_023646608.1</u>, rabbit <u>EU195061.1</u> (Autry and Thomas 2008), and dog <u>U94345.1</u> (83). (3) SLN: horse (40), rabbit <u>U96091.1</u> (73), mouse <u>NM_025540.2</u>, and human <u>U96094.1</u> (73). (4) PLN: horse (40), rabbit <u>Y00761.1</u> (197), human <u>M63603.1</u> (198), and dog <u>NM_001003332.1</u> (199). (5) MRLN: horse (40) and human <u>NM_001304732.2</u> (21). (6) DWORF: horse (40) and mouse <u>NM_001369305.1</u>(22). (7) GP: horse muscle <u>NP_001138725.1</u>, rabbit muscle <u>NP_001075653.1</u>, and human muscle <u>AH002957.2</u>. The cDNA sequence for an accession code can be identified through the NCBI GenBank utility at https://www.ncbi.nlm.nih.gov/nuccore/.

## SUPPORTING INFORMATION^3^

### TABLE#

Table S1 Amino acid sequence variations of horse SERCA1a versus rabbit, mouse, and human orthologs, correlated with site-directed mutagenesis and a human skin disease mutation.

### FIGURES#

Figure S1 Amino acid sequence alignment of SERCA1a orthologs: horse, rabbit, mouse, and human.

Figure S2 Molecular modeling of horse SERCA, with structural analysis of two key variations in amino acid sequence versus rabbit SERCA.

Figure S3 Standard purification of SR vesicles from rabbit muscle using mechanical homogenization and differential centrifugation.

Figure S4 Coomassie gel of horse and rabbit SR preps utilized in this study.

Figure S5 Semi-quantitative immunoblot analysis of SERCA protein expression in horse muscle fractions.

Figure S6 Immunoblot analysis using anti-horse-SLN pAb GS3379 detects minimal expression of SLN protein in horse SR.

Figure S7 Immunoblot analysis using anti-rabbit-SLN pAb ABT13 identifies SLN protein expression in rabbit SR, but not horse SR.

Figure S8 Immunoblot analysis using anti-mouse-SLN pAb PFD1-1 identifies SLN protein expression in rabbit SR, but not horse SR.

Figure S9 Immunoblot analysis using anti-human-SLN pAb 18395-1-AP does not detect horse or rabbit SLN, but instead shows non-specific binding to other proteins in horse and rabbit SR.

Figure S10 Immunoblot analysis using anti-universal-PLN mAb 2D12 detects minimal expression of PLN protein in horse and rabbit SR.

Figure S11 Horse and rabbit SERCA show similar temperature dependence of Ca^2+^-activated ATPase activity.

Figure S12 Horse and rabbit SERCA show similar Ca^2+^-dependent cleavage by Proteinase K.

