## Supplement: 1 Table, 12 Figures, 22 References for "Horse gluteus is a null-sarcolipin muscle with enhanced sarcoplasmic reticulum calcium transport"

p. S1 **Table of Contents**

p. S2–S3 **Fig. S1**. Amino acid sequence alignment of SERCA1a orthologs: horse, rabbit, mouse, and human.

p. S4–S5 **Table S1**. Amino acid sequence variations of horse SERCA1a versus rabbit, mouse, and human orthologs, correlated with functional effects of 18 site-directed mutants and one human skin disease mutation.

p. S6 **Fig. S2**. Molecular modeling of horse SERCA, with structural analysis of two key variations in amino acid sequence versus rabbit SERCA.

p. S7 **Fig. S3**. Standard purification of SR vesicles from rabbit muscle using mechanical homogenization and differential centrifugation.

p. S8 **Fig. S4**. Coomassie gel of horse and rabbit SR preps utilized in this study.

p. S9 **Fig. S5**. Semi-quantitative immunoblot analysis of SERCA protein expression in horse muscle fractions.

p. S10 **Fig. S6**. Immunoblot analysis using custom anti-horse-SLN pAb GS3379 detects minimal expression of SLN protein in horse SR.

p. S11 **Fig. S7**. Immunoblot analysis using commercial anti-rabbit/mouse/human-SLN pAb ABT13 identifies SLN protein expression in rabbit SR, but not horse SR.

p. S12 **Fig. S8**. Immunoblot analysis using custom anti-rabbit/mouse/human-SLN pAb PFD1-1 identifies SLN protein expression in rabbit SR, but not horse SR.

p. S13 **Fig. S9**. Immunoblot analysis using commercial anti-human-SLN pAb 18395-1-AP does not detect horse or rabbit SLN, but instead shows non-specific binding to other proteins in horse and rabbit SR.

p. S14 **Fig. S10**. Immunoblot analysis using commercial anti-universal-PLN mAb 2D12 detects minimal expression of PLN protein in horse and rabbit SR.

p. S15 **Fig. S11**. Horse and rabbit SERCA show similar temperature dependence of $Ca^{2+}$-activated ATPase activity.

p. S16 **Fig. S12**. Horse and rabbit SERCA show similar $Ca^{2+}$-dependent cleavage by ProtK.

p. S17–S18 **References**

#### FIGURES and TABLE

**Figure S1**. **Amino acid sequence alignment of SERCA1a orthologs: horse, rabbit, mouse, and human.** Nucleotide and protein sequences for horse *ATP2A1* were determined from the horse reference genome EquCab2, Ensembl version 86. The protein sequence of SERCA1a orthologs from rabbit, mouse, and human were obtained from the NCBI database with GenBank accession code ABW96358.1, NP_031530.2, and NM_004320.4 (1), respectively. The amino acid sequence of horse and rabbit SERCA1a differ at 48 residues, as illustrated below (*yellow highlight*). Horse SERCA1a also has a 1-residue deletion (*orange highlight*) at position 504, as compared to SERCA1a from rabbit, mouse, and human. Amino acid sequence variations between horse and rabbit SERCA1a are further identified in the molecular model of horse SERCA1a (**Figure 2*A***) and the structure-function analysis compiled from reports of SERCA site-directed mutagenesis (**Table S1**).

```
                         |         |         |         |         |         |
horse     MEAAHSKTTEECLAYFGVSETAGLTPEQVKRHLEKYGPNELPTEEGKSLWELVVEQFEDL  60
rabbit    MEAAHSKSTEECLAYFGVSETTGLTPDQVKRHLEKYGHNELPAEEGKSLWELVIEQFEDL  60
mouse     MEAAHSKSTEECLSYFGVSETTGLTPDQVKRHLEKYGPNELPAEEGKSLWELVVEQFEDL  60
human     MEAAHAKTTEECLAYFGVSETTGLTPDQVKRNLEKYGLNELPAEEGKTLWELVIEQFEDL  60

                         |         |         |         |         |         |
horse     LVRILLLAACISFVLAWFEEGEETVTAFVEPFVILLILIANAIVGVWQERNAENAIEALK 120
rabbit    LVRILLLAACISFVLAWFEEGEETITAFVEPFVILLILIANAIVGVWQERNAENAIEALK 120
mouse     LVRILLLAACISFVLAWFEEGEETVTAFVEPFVILLILIANAIVGVWQERNAENAIEALK 120
human     LVRILLLAACISFVLAWFEEGEETITAFVEPFVILLILIANAIVGVWQERNAENAIEALK 120

                         |         |         |         |         |         |
horse     EYEPEMGKVYRADRKSVQRIKARDIVPGDIVEVAVGDKVPADIRILSIKSTTLRVDQSIL 180
rabbit    EYEPEMGKVYRADRKSVQRIKARDIVPGDIVEVAVGDKVPADIRILSIKSTTLRVDQSIL 180
mouse     EYEPEMGKVYRADRKSVQRIKARDIVPGDIVEVAVGDKVPADIRILSIKSTTLRVDQSIL 180
human     EYEPEMGKVYRADRKSVQRIKARDIVPGDIVEVAVGDKVPADIRILAIKSTTLRVDQSIL 180

                         |         |         |         |         |         |
horse     TGESVSVIKHTEPVPDPRAVNQDKKNMLFSGTNIAAGKALGIVAATGVNTEIGKIRDQMA 240
rabbit    TGESVSVIKHTEPVPDPRAVNQDKKNMLFSGTNIAAGKALGIVATTGVSTEIGKIRDQMA 240
mouse     TGESVSVIKHTDPVPDPRAVNQDKKNMLFSGTNIAAGKAVGIVATTGVSTEIGKIRDQMA 240
human     TGESVSVIKHTEPVPDPRAVNQDKKNMLFSGTNIAAGKALGIVATTGVGTEIGKIRDQMA 240

                         |         |         |         |         |         |
horse     ATEQDKTPLQQKLDEFGEQLSKVISLICVAVWLINIGHFNDPVHGGSWLRGAIYYFKIAV 300
rabbit    ATEQDKTPLQQKLDEFGEQLSKVISLICVAVWLINIGHFNDPVHGGSWIRGAIYYFKIAV 300
mouse     ATEQDKTPLQQKLDEFGEQLSKVISLICVAVWLINIGHFNDPVHGGSWFRGAIYYFKIAV 300
human     ATEQDKTPLQQKLDEFGEQLSKVISLICVAVWLINIGHFNDPVHGGSWFRGAIYYFKIAV 300

                         |         |         |         |         |         |
horse     ALAVAAIPEGLPAVITTCLALGTRRMAKKNAIVRSLPSVETLGCTSVICSDKTGTLTTNQ 360
rabbit    ALAVAAIPEGLPAVITTCLALGTRRMAKKNAIVRSLPSVETLGCTSVICSDKTGTLTTNQ 360
mouse     ALAVAAIPEGLPAVITTCLALGTRRMAKKNAIVRSLPSVETLGCTSVICSDKTGTLTTNQ 360
human     ALAVAAIPEGLPAVITTCLALGTRRMAKKNAIVRSLPSVETLGCTSVICSDKTGTLTTNQ 360

                         |         |         |         |         |         |
horse     MSVCKMFIVDKVDGDLCILNEFSITGSTYAPEGEILKNDKPVRAGQYDGLVEVATICALC 420
rabbit    MSVCKMFIIDKVDGDFCSLNEFSITGSTYAPEGEVLKNDKPIRSGQFDGLVELATICALC 420
mouse     MSVCKMFIIDKVDGDVCSLNEFSITGSTYAPEGEVLKNDKPVRAGQYDGLVELATICALC 420
human     MSVCKMFIIDKVDGDICLLNEFSITGSTYAPEGEVLKNDKPVRPGQYDGLVELATICALC 420
```

```
                         |         |         |         |         |         |
horse    NDSSLDFNEAKGVYEKVGEATETALTTLVEKMNVFNTDVRNLSKVERANACNSVIRQLMK 480
rabbit   NDSSLDFNETKGVYEKVGEATETALTTLVEKMNVFNTEVRNLSKVERANACNSVIRQLMK 480
mouse    NDSSLDFNETKGVYEKVGEATETALTTLVEKMNVFNTEVRSLSKVERANACNSVIRQLMK 480
human    NDSSLDFNEAKGVYEKVGEATETALTTLVEKMNVFNTDVRSLSKVERANACNSVIRQLMK 480
                         |         |          |         |         |         |
horse    KEFTLEFSRDRKSMSVYCSPAKS-RAAVGNKMFVKGAPEGVLDRCNYVRVGTTRVPMAGPV 540
rabbit   KEFTLEFSRDRKSMSVYCSPAKSSRAAVGNKMFVKGAPEGVIDRCNYVRVGTTRVPMTGPV 541
mouse    KEFTLEFSRDRKSMSVYCSPAKSSRAAVGNKMFVKGAPEGVIDRCNYVRVGTTRVPLTGPV 541
human    KEFTLEFSRDRKSMSVYCSPAKSSRAAVGNKMFVKGAPEGVIDRCNYVRVGTTRVPLTGPV 541

                         |         |         |         |         |         |
horse    KERILSVIKEWGTGRDTLRCLALATRDTPPKREDMILDDSARFMEYETDLTFIGVVGMLD 600
rabbit   KEKILSVIKEWGTGRDTLRCLALATRDTPPKREEMVLDDSSRFMEYETDLTFVGVVGMLD 601
mouse    KEKIMSVIKEWGTGRDTLRCLALATRDTPPKREEMVLDDSAKFMEYEMDLTFVGVVGMLD 601
human    KEKIMAVIKEWGTGRDTLRCLALATRDTPPKREEMVLDDSARFLEYETDLTFVGVVGMLD 601

                         |         |         |         |         |         |
horse    PPRKEVTGSIQLCRDAGIRVIMITGDNKGTAIAICRRIGIFGENEEVADRAYTGREFDDL 660
rabbit   PPRKEVMGSIQLCRDAGIRVIMITGDNKGTAIAICRRIGIFGENEEVADRAYTGREFDDL 661
mouse    PPRKEVTGSIQLCRDAGIRVIMITGDNKGTAIAICRRIGIFSENEEVTDRAYTGREFDDL 661
human    PPRKEVTGSIQLCRDAGIRVIMITGDNKGTAIAICRRIGIFGENEEVADRAYTGREFDDL 661

                         |         |         |         |         |         |
horse    PLAEQREACRRACCFARVEPSHKSKIVEYLQSFDEITAMTGDGVNDAPALKKAEIGIAMG 720
rabbit   PLAEQREACRRACCFARVEPSHKSKIVEYLQSYDEITAMTGDGVNDAPALKKAEIGIAMG 721
mouse    PLAEQREACRRACCFARVEPSHKSKIVEYLQSYDEITAMTGDGVNDAPALKKAEIGIAMG 721
human    PLAEQREACRRACCFARVEPSHKSKIVEYLQSYDEITAMTGDGVNDAPALKKAEIGIAMG 721

                         |         |         |         |         |         |
horse    SGTAVAKTASEMVLADDNFSTIVAAVEEGRAIYNNMKQFIRYLISSNVGEVVCIFLTAAL 780
rabbit   SGTAVAKTASEMVLADDNFSTIVAAVEEGRAIYNNMKQFIRYLISSNVGEVVCIFLTAAL 781
mouse    SGTAVAKTASEMVLADDNFSTIVAAVEEGRAIYNNMKQFIRYLISSNVGEVVCIFLTAAL 781
human    SGTAVAKTASEMVLADDNFSTIVAAVEEGRAIYNNMKQFIRYLISSNVGEVVCIFLTAAL 781

                         |         |         |         |         |         |
horse    GLPEALIPVQLLWVNLVTDGLPATALGFNPPDLDIMDRPPRSPKEPLISGWLFFRYMAIG 840
rabbit   GLPEALIPVQLLWVNLVTDGLPATALGFNPPDLDIMDRPPRSPKEPLISGWLFFRYMAIG 841
mouse    GLPEALIPVQLLWVNLVTDGLPATALGFNPPDLDIMDRPPRSPKEPLISGWLFFRYMAIG 841
human    GLPEALIPVQLLWVNLVTDGLPATALGFNPPDLDIMDRPPRSPKEPLISGWLFFRYMAIG 841

                         |         |         |         |         |         |
horse    GYVGAATVGAAAWWFLFAEDGPHVTYSQLTHFMKCNEHNPDFEGVDCEVFEAPEPMTMAL 900
rabbit   GYVGAATVGAAAWWFMYAEDGPGVTYHQLTHFMQCTEDHPHFEGLDCEIFEAPEPMTMAL 901
mouse    GYVGAATVGAAAWWFLYAEDGPHVSYHQLTHFMQCTEHNPEFDGLDCEVFEAPEPMTMAL 901
human    GYVGAATVGAAAWWFLYAEDGPHVNYSQLTHFMQCTEDNTHFEGIDCEVFEAPEPMTMAL 901

                         |         |         |         |         |         |
horse    SVLVTIEMCNALNSLSENQSLVRMPPWVNIWLVGSICLSMSLHFLILYVDPLPMIFKLEA 960
rabbit   SVLVTIEMCNALNSLSENQSLMRMPPWVNIWLLGSICLSMSLHFLILYVDPLPMIFKLKA 961
mouse    SVLVTIEMCNALNSLSENQSLLRMPPWVNIWLLGSICLSMSLHFLILYVDPLPMIFKLRA 961
human    SVLVTIEMCNALNSLSENQSLLRMPPWVNIWLLGSICLSMSLHFLILYVDPLPMIFKLRA 961

                         |         |         |
horse    LDLTHWLMVLKISFPVILLDEVLKFVARNYLEG 993
rabbit   LDLTQWLMVLKISLPVIGLDEILKFIARNYLEG 994
mouse    LDFTQWLMVLKISLPVIGLDELLKFIARNYLEG 994
human    LDLTQWLMVLKISLPVIGLDEILKFVARNYLEG 994
```

**Table S1. Amino acid sequence variations of horse SERCA1a versus rabbit, mouse, and human orthologs, correlated with functional effects of 18 site-directed mutants and one human skin disease mutation.** There are 48 residue variations between horse and rabbit SERCA1a sequences, which are highlighted in *yellow*, plus a 1-residue deletion in horse at position 504, highlighted in *orange* (*left columns*; see also **Figure 2*A*** and **Figure S1**). Residues which result in perturbation of SERCA activity when subjected to site-directed mutagenesis are listed (*right columns*), as compiled from references 2–8 (2-8). A molecular model of horse SERCA (**Figure 3**, **Figure S2**) and the crystal structure of rabbit SERCA were used to examine the side-chain interactions of (i) residue 54, which is the position of a human SERCA2b variation correlated with Darier's skin disease (7), and (ii) residue 959/960, which encodes a negatively-charged sidechain at position 959 in horse SERCA (E959) versus a positively-charged sidechain in the analogous position (960) encoded by rabbit (K960), mouse (R960), and human (R960) orthologs (**Figure S1**).

| Residue | Horse | Rabbit | Mouse | Human | Site-Directed Mutant | Effect | Ref |
|---|---|---|---|---|---|---|---|
| 6 | S | S | S | A | S6L-SERCA1a, rabbit | WT activity | 2 |
| 8 | T | S | S | T | S8A-SERCA1a, rabbit | low expresion (COS-1) | 2 |
| 14 | A | A | S | A | | | |
| 22 | A | T | T | T | | | |
| 27 | E | D | D | D | | | |
| 32 | H | H | H | N | | | |
| 38 | P | H | P | L | H38A-SERCA1a, rabbit | WT activity | 3 |
| 43 | T | A | A | A | A43G-SERCA1a, rabbit | WT activity | 3 |
| 48 | S | S | S | T | S48A-SERCA1a, rabbit | Ca-ATPase increase | 3 |
| 54 | V | I | V | I | I54V -SERCA1a, rabbit<br>I54V-SERCA2b, human | Ca-ATPase decrease;<br>Darier's skin disease | 3,4 |
| 85 | V | I | V | I | | | |
| 192 | E | E | D | E | E192A-SERCA1a, rabbit | WT activity | 5 |
| 220 | L | L | V | L | | | |
| 225 | A | T | T | T | | | |
| 229 | N | S | S | G | | | |
| 289 | L | I | F | F | | | |
| 369 | V | I | I | I | | | |
| 376 | L | F | V | I | | | |
| 378 | I | S | S | L | S378R-SERCA2a, rabbit | WT activity | 6 |
| 395 | I | V | V | V | V395A-SERCA2a, rabbit | WT activity | 6 |
| 402 | V | I | V | V | V402A-SERCA2a, rabbit | WT activity | 6 |
| 404 | A | S | A | P | | | |
| 407 | Y | F | Y | Y | | | |
| 413 | V | L | L | L | | | |
| 430 | A | T | T | A | | | |
| 458 | D | E | E | D | | | |
| 461 | N | N | S | S | | | |
| 504 | – | S | S | S | | | |
| 522 | L | I | I | I | | | |

| | | | | | | | |
|---|---|---|---|---|---|---|---|
| 536 | M | M | L | L | | | |
| 537 | A | T | T | T | | | |
| 543 | R | K | K | K | | | |
| 545 | L | L | M | M | | | |
| 546 | S | S | S | A | | | |
| 574 | D | E | E | E | | | |
| 576 | I | V | V | V | | | |
| 581 | A | S | A | A | | | |
| 582 | R | R | K | R | | | |
| 584 | M | M | M | L | | | |
| 588 | T | T | M | T | | | |
| 594 | I | V | V | V | | | |
| 607 | T | M | T | T | | | |
| 642 | G | G | S | G | | | |
| 648 | A | A | T | A | | | |
| 693 | F | Y | Y | Y | | | |
| 856 | L | M | L | L | | | |
| 857 | F | Y | Y | Y | Y858A-SERCA1a, rabbit | WT activity | 7 |
| 863 | H | G | H | H | | | |
| 865 | T | T | S | N | T866A-SERCA1a, rabbit | WT activity | 7 |
| 867 | S | H | H | S | | | |
| 874 | K | Q | Q | Q | Q875A-SERCA1a, rabbit | WT activity | 7 |
| 876 | N | T | T | T | T877A-SERCA1a, rabbit | WT activity | 7 |
| 878 | H | D | H | D | D879A-SERCA1a, rabbit | WT activity | 7 |
| 879 | N | H | N | N | | | |
| 880 | P | P | P | T | P881A-SERCA1a, rabbit | WT activity | 7 |
| 881 | D | H | E | H | | | |
| 883 | E | E | D | E | E884A-SERCA1a, rabbit | WT activity | 7 |
| 885 | V | L | L | I | | | |
| 889 | V | I | V | V | | | |
| 922 | V | M | L | L | | | |
| 933 | V | L | L | L | | | |
| 959 | E | K | R | R | T960A-SERCA1a, chicken | low expression; uncoupled transport/ATPase | 8 |
| 963 | L | L | F | L | | | |
| 965 | H | Q | Q | Q | | | |
| 974 | F | L | L | L | | | |
| 978 | L | G | G | G | | | |
| 982 | V | I | L | I | | | |
| 986 | V | I | I | V | | | |

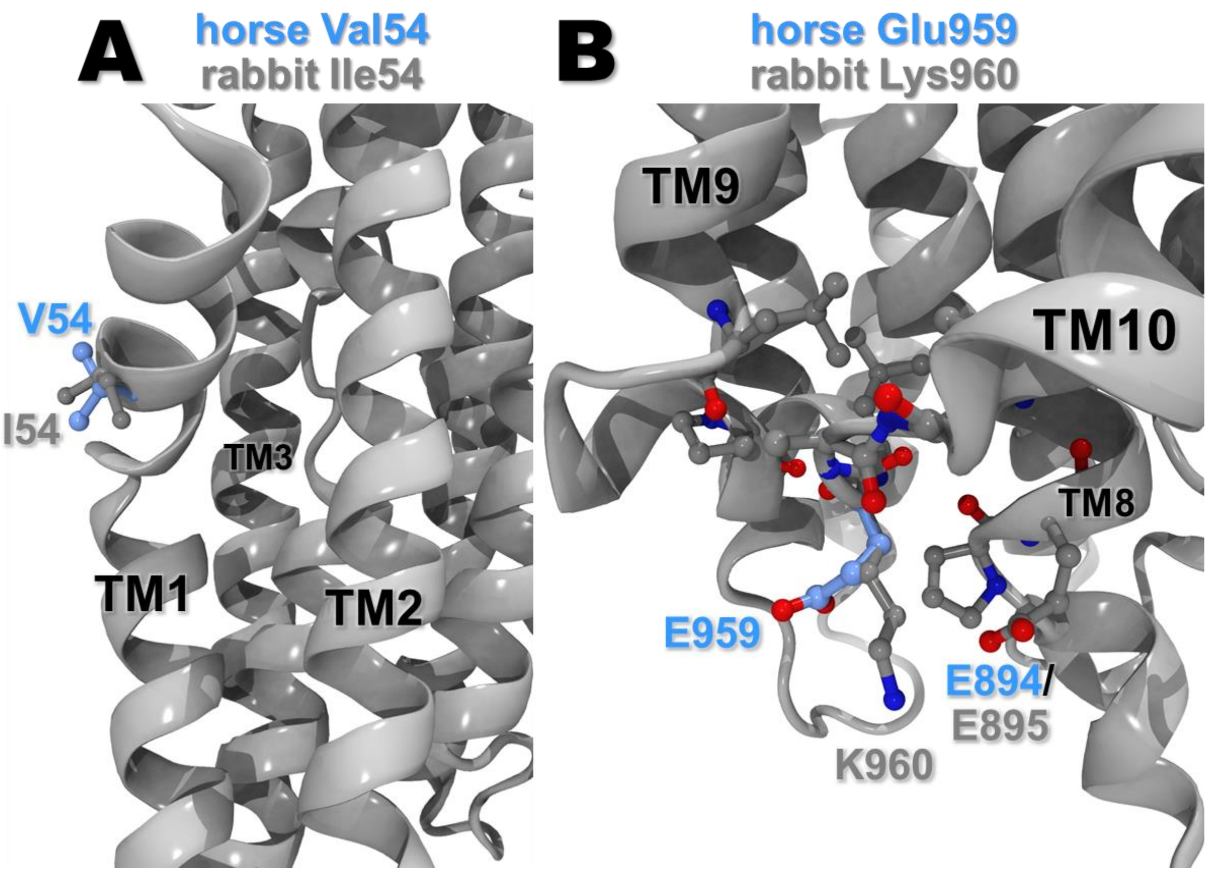


**Figure S2**. **Molecular modeling of horse SERCA, with structural analysis of two key variations in amino acid sequence versus rabbit SERCA.** *A*, molecular modeling of SERCA was used to compare residue V54 of horse SERCA1a with I54 of rabbit SERCA1a in the E1▪Mg b crystal structure (PDB ID code 3W5A (9)) (see also **Figure 2*A***). In rabbit SERCA1a, the engineered site-directed mutant I54V results in decreased ATPase activity (4). In human SERCA2b, the I54V mutation is associated with Darier's skin disease (7). Modeling and structural analysis of horse and rabbit SERCA1a indicate that V54 and I54 sidechains show similar positions facing the lipids just below the water–phospholipid headgroup interface (myoplasmic leaflet) in the E2•TG and E1P•2Ca•ADP structural states (rabbit SERCA structures PDB ID codes 1IWO and 2ZBD, respectively) (10,11). In the E1•2Ca crystal structure of rabbit SERCA1a (PDB ID codes 1SU4 (12)), I54 is located in the interface between TM1 and TM3, whereby rabbit I54 makes van der Waals contacts with side chains of residues Trp50 and Val53 on TM1, plus the Cα atom of Gly257 on TM3. In the molecular model of horse SERCA1a, the Val54 sidechain does not reach far enough to contact Gly257, thereby decreasing van der Waals contributions here in horse SERCA1a. *B*, structural comparison of residues E959 and K960 in horse and rabbit SERCA1a (equivalent position in the amino acid sequence of the two orthologs). Modeling of SERCA1a indicates that horse E959 and rabbit K600 sidechains show different positions near the water–phospholipid bilayer luminal interface due to opposite electrostatic force (repulsive versus attractive) with residue E894/E895 on TM8, per horse and rabbit amino acid sequence numbering, respectively. Chicken SERCA1a encodes T960 at this position, and the engineered site-directed mutant T960A partially uncouples ATPase activity from $Ca^{2+}$ transport, producing a low efficiency of pump energetics, i.e., a low Ca/ATP coupling ratio (8).

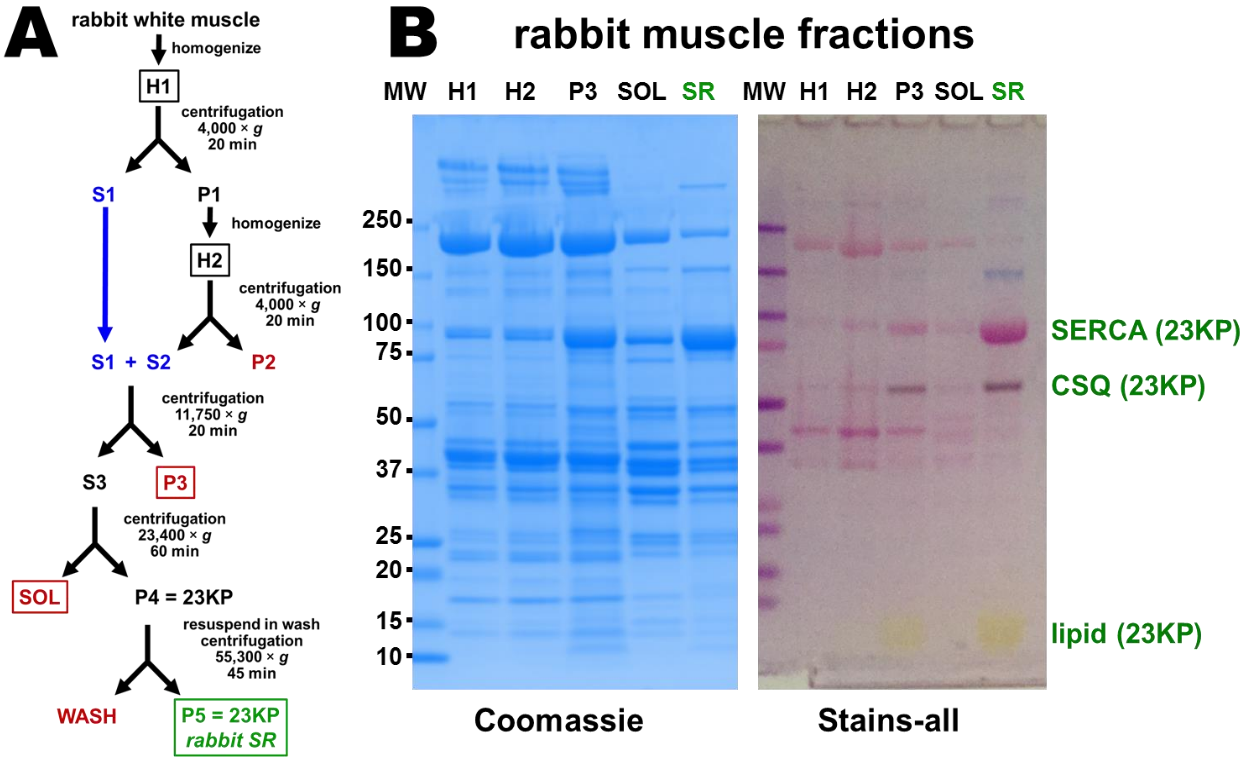


**Figure S3**. **Standard purification of SR vesicles from rabbit muscle using mechanical homogenization and differential centrifugation.** Unfractionated SR vesicles, with minimal glycogen phosphorylase contamination (**Fig. 4**), were isolated using the classic protocol of Ikemoto *et al*. (13). *A*, rabbit white muscle (fast-twitch skeletal) was homogenized by two rounds of Polytron generator (H1, H2). Five muscle fractions were characterized biochemically, as indicated by *boxed text*: homogenate 1 (H1), homogenate 2 (H2), 11,700 × *g* pellet (P3), 234000 × *g* supernatant (SOL), and a 23,400 × *g* pellet which is washed once to yield rabbit "SR vesicles" (P5 = 23KP). *B*, rabbit muscle fractions were electrophoresed through 4–15% Laemmli gels and stained with Coomassie blue (*left*) or Stains-all (*right*). The amount of protein loaded was 10 µg per lane. The molecular mass of protein gel markers (kDa) are indicated on the *left*. Gel bands are identified on the *right*, along with identification of the muscle fraction with predominant enrichment for each band.

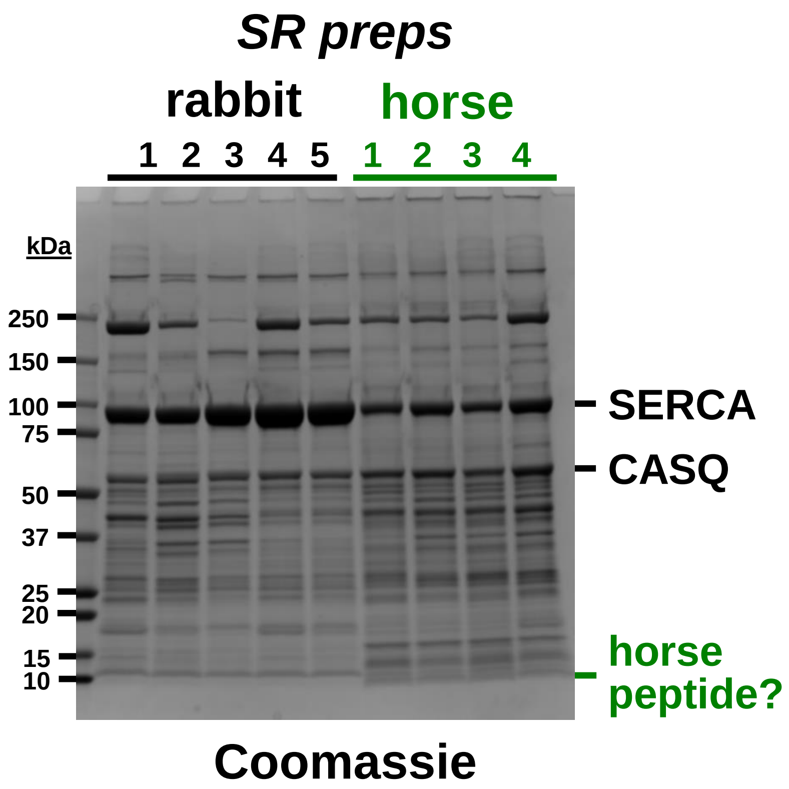


**Figure S4. Coomassie gel of rabbit and horse SR preps utilized in this study.** Five rabbit SR preps (washed 23KP = P5, per the prep flowchart in **Figure S3**) and four horse SR preps (10KP = P3, as in the prep flowchart in **Figure 4**) were electrophoresed through a 4–15% Laemmli gel and stained with Coomassie blue. The amount of protein loaded was 15 µg per lane, as determined by BCA assay with BSA as protein standard. The molecular mass of protein gel markers (kDa) are indicated on the *left*. Gel bands are identified on the *right*.

Coomassie densitometry determined that **horse SR vesicles show** (a) **a level of SERCA expression** that is ~45% relative to SERCA expression in rabbit SR vesicles (SERCA content per total SR protein**)**, (b) **a level of CASQ expression** that is ~120% relative to CASQ expression in rabbit SR vesicles (CASQ content per total SR protein), and (c) **a ratio of CASQ/SERCA expression levels** that is ~3.0 higher than the ratio of CASQ/SERCA in rabbit SR vesicles.

### SERCA immunoblot

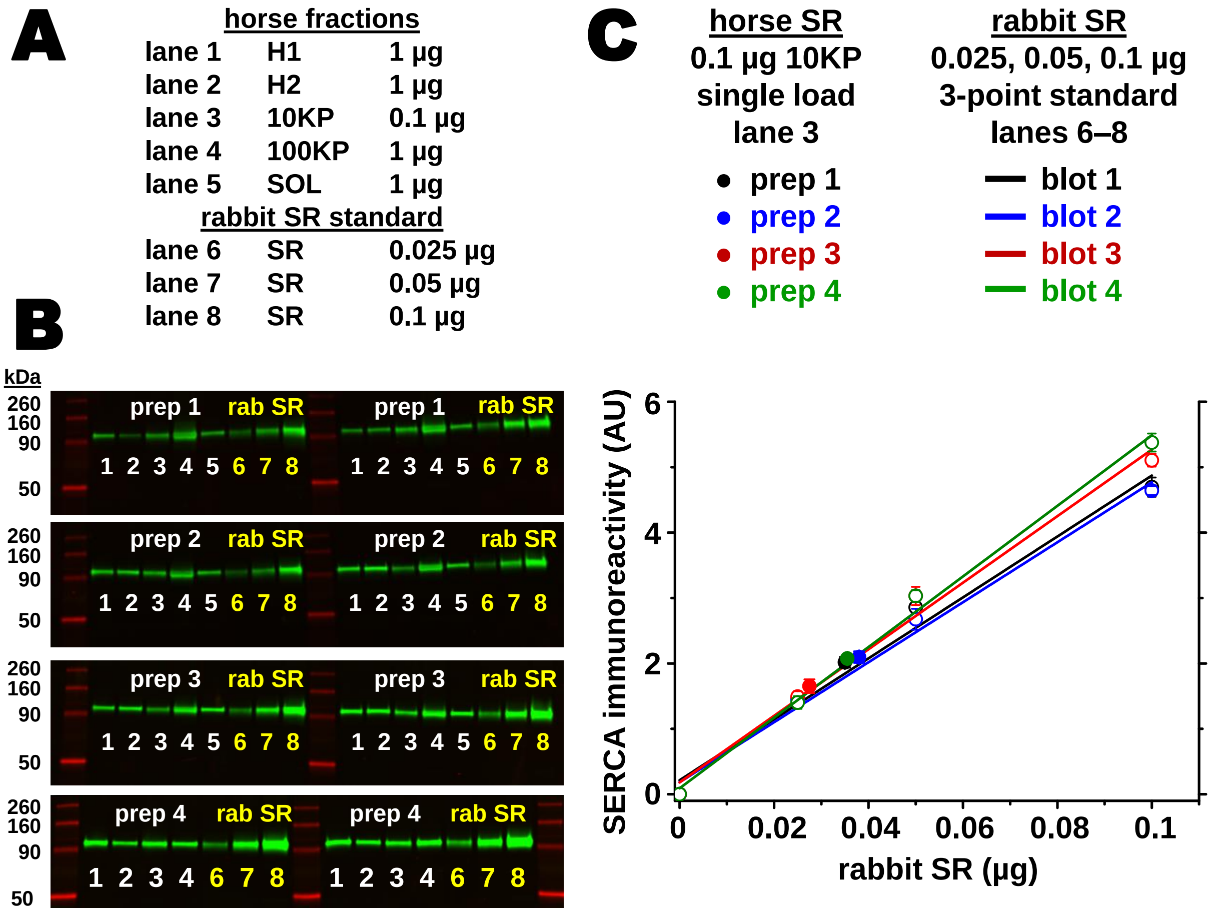


**Figure S5**. **Semi-quantitative immunoblot analysis of SERCA expression in horse muscle fractions.** *A*, SERCA immunoblots analyzed horse muscle fractions (H1, H2, 10KP, 100KP, SOL) from four preps, and SR vesicles from rabbit muscle were used as a quantitative standard. The protein load (µg) in each lane is listed. *B*, samples were electrophoresed through 4–15% Laemmli gels. Four horse preps were analyzed in duplicate on each blot (*left*, *right*), along with rabbit SR. Immunoblotting was performed using the mAb VE12$_1$G9 (primary) (14) and goat anti-mouse-IgG pAb labeled with a 800-nm fluorophore (secondary). Immunolabeling was quantified using the LI-COR laser scanner system in the near-infrared fluorescence mode. The intensity scaling of each fluorescence image is the same for all four immunoblots. The molecular mass of protein gel markers are indicated on the *left*. *C*, linear regression analysis (*open circles*: average ± SEM) of rabbit SR standard (0.025, 0.05, and 0.1 µg loads) on immunoblots 1–4. Semi-quantitative analysis (*filled circle*: duplicate average ± SEM) of SERCA content in the 10KP fraction (0.1 µg load) of horse preps 1–4. In some cases, the error bars (SEM) are smaller than the symbol (open or filled circles).

<u>Semi-quantitative immunoblotting</u> determined that **horse SR vesicles show a level of SERCA expression** that is 35 ± 6.6% relative to SERCA expression in rabbit SR vesicles (SERCA content per total SR protein).

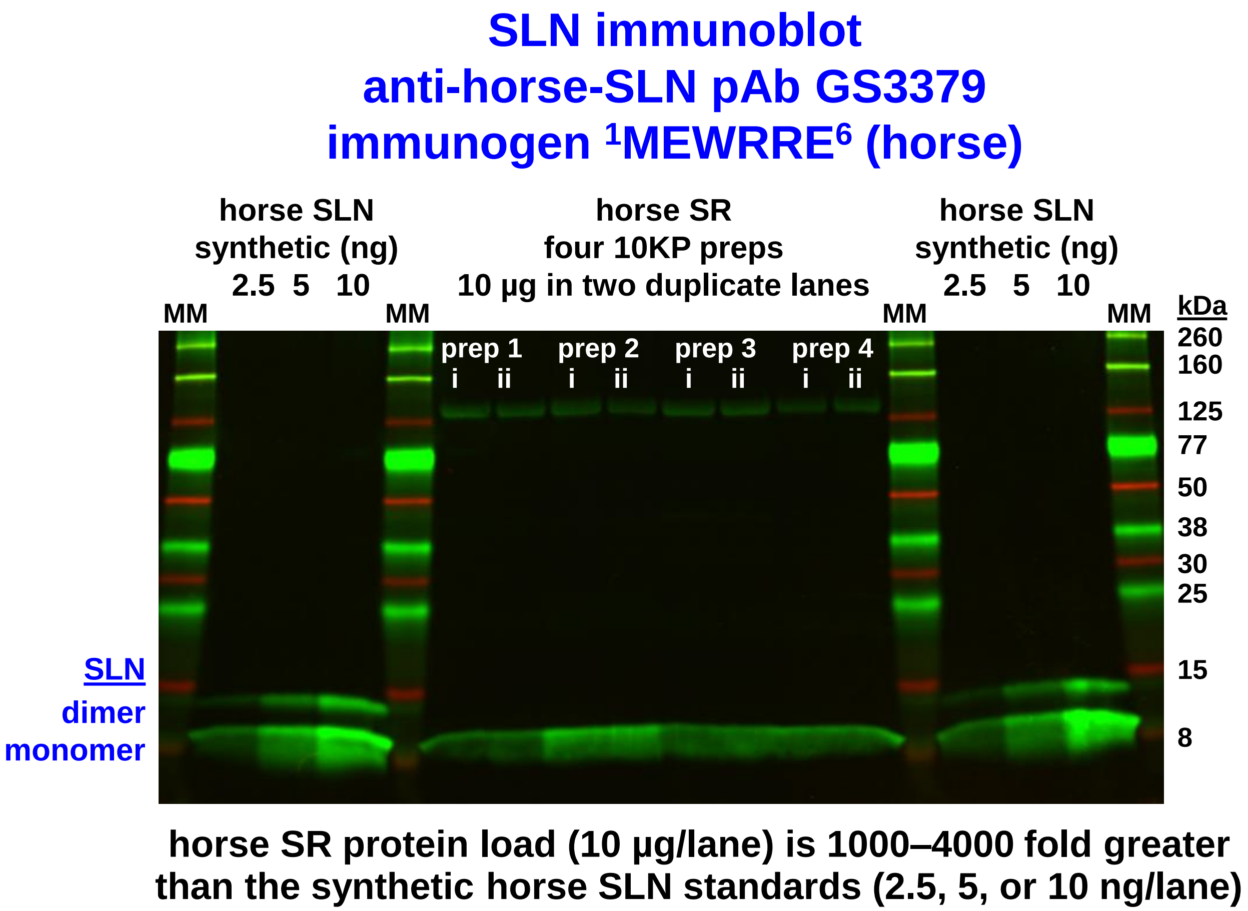


**Figure S6**. **Immunoblot analysis using custom anti-horse-SLN pAb GS3379 detects minimal expression of SLN protein in horse SR.** The primary antibody was the custom anti-horse-SLN pAb GS3379, with immunogen comprising horse residues $^{1}$MEWRRE$^{6}$ (15). Samples were electrophoresed through a 4–20% Laemmli gel. Horse SR vesicles were assayed (N = 4 preps; 10 µg protein per lane; each prep run in duplicate lanes). Synthetic horse SLN was used as a semi-quantitative protein standard (2.5, 5, or 10 ng/lane) in duplicate sets on *left* and *right* sides on the gel. Immunoblotting was performed using pAb GS3379 (primary) and goat anti-rabbit-IgG pAb labeled with 800-nm fluorophore (secondary). Immunolabeling was quantified using the LI-COR laser scanner system in the near-infrared fluorescence mode. The molecular mass of protein gel markers (kDa) are indicated on the *right*. The proposed molecular species of SLN (monomer, dimer) are indicated on the *left*.

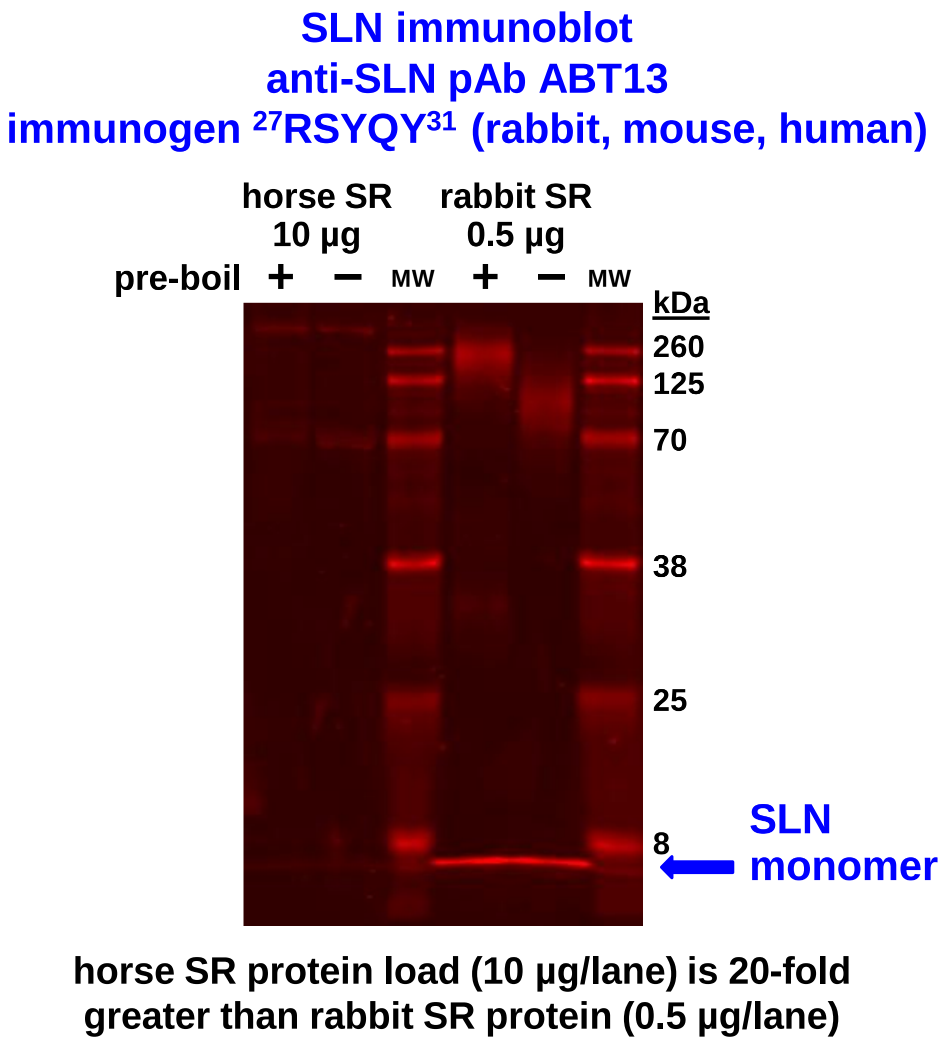


**Figure S7**. **Immunoblot analysis using commercial anti-rabbit/mouse/human-SLN pAb ABT13 identifies SLN protein expression in rabbit SR, but not horse SR.** The primary antibody was the commercial anti-SLN pAb ABT13, with immunogen comprising rabbit/mouse/human SLN residues $^{27}$RSYQY$^{31}$, whereas horse SLN encodes $^{26}$RSYQ$^{29}$ (15). Samples were electrophoresed through a 10% Laemmli gel. Horse SR vesicles were loaded at 10 µg protein per lane (*left*), and rabbit SR vesicles were loaded at 0.5 µg protein per lane (*right*), a 20-fold lower amount than horse SR. Immunoblotting was performed using pAb ABT13 (primary) and goat anti-rabbit-IgG pAb labeled with 680-nm fluorophore (secondary). Immunolabeling was quantified using the LI-COR laser scanner system in the near-infrared fluorescence mode. One sample of each SR set was heated at 100 °C for 2 min in Laemmli sample buffer prior to electrophoresis (+ pre-boil). The molecular mass of protein gel markers (kDa) are indicated on the *right*.

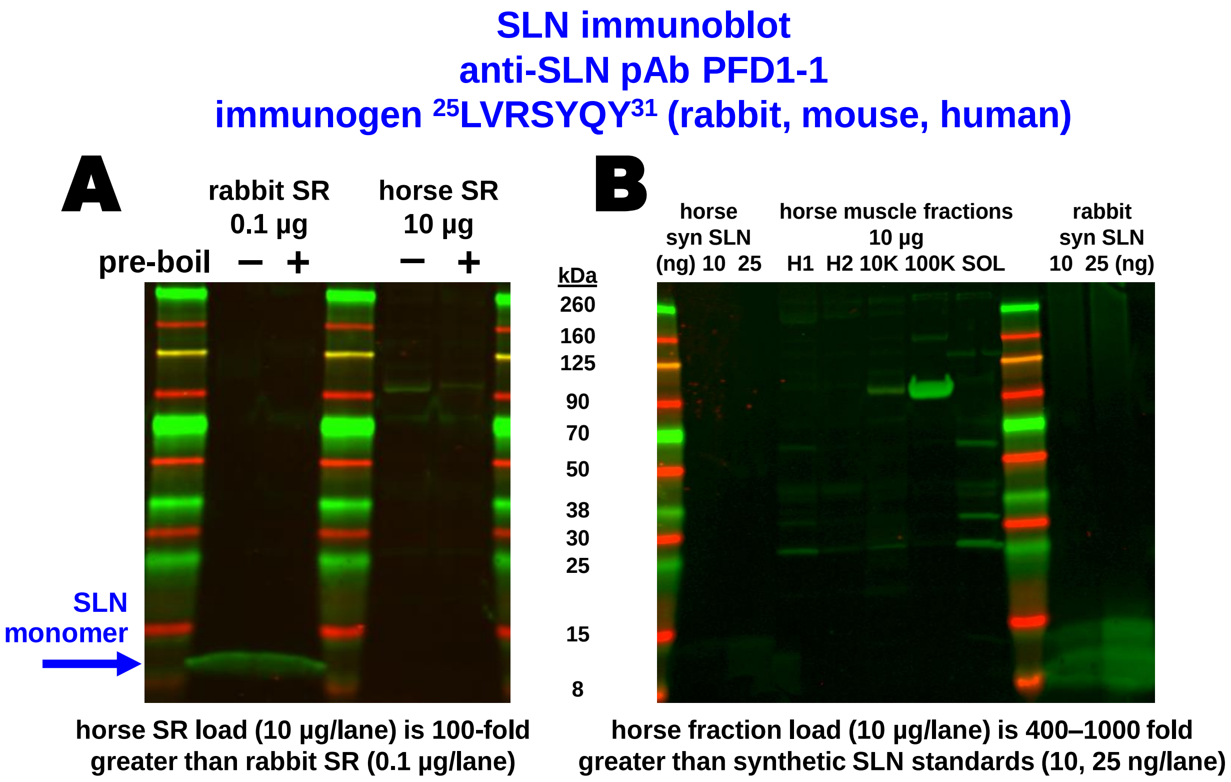


**Figure S8**. **Immunoblot analysis using custom anti-rabbit/mouse/human-SLN pAb PFD1-1 identifies SLN protein expression in rabbit SR, but not horse SR.** The primary antibody was the custom anti-SLN pAb PFD1-1 (16) with immunogen comprising rabbit/mouse/human SLN residues $^{25}$LVRSYQY$^{31}$, whereas horse SLN encodes $^{24}$LVRSYQ$^{29}$ (15). Samples were electrophoresed through 4–20% Laemmli gels. Horse SR vesicles were loaded at 10 µg protein per lane (*left*), and rabbit SR vesicles were loaded at 0.5 µg protein per lane (*right*), a 20-fold lower amount than horse SR. Immunoblotting was performed using pAb ABT13 (primary) and goat anti-rabbit-IgG pAb labeled with 680-nm fluorophore (secondary). Immunolabeling was quantified using the LI-COR laser scanner system in the near-infrared fluorescence mode. One sample of each SR set was heated at 100 °C for 2 min in Laemmli sample buffer prior to electrophoresis (+ pre-boil). The molecular mass of protein gel markers (kDa) are indicated in the *middle*.

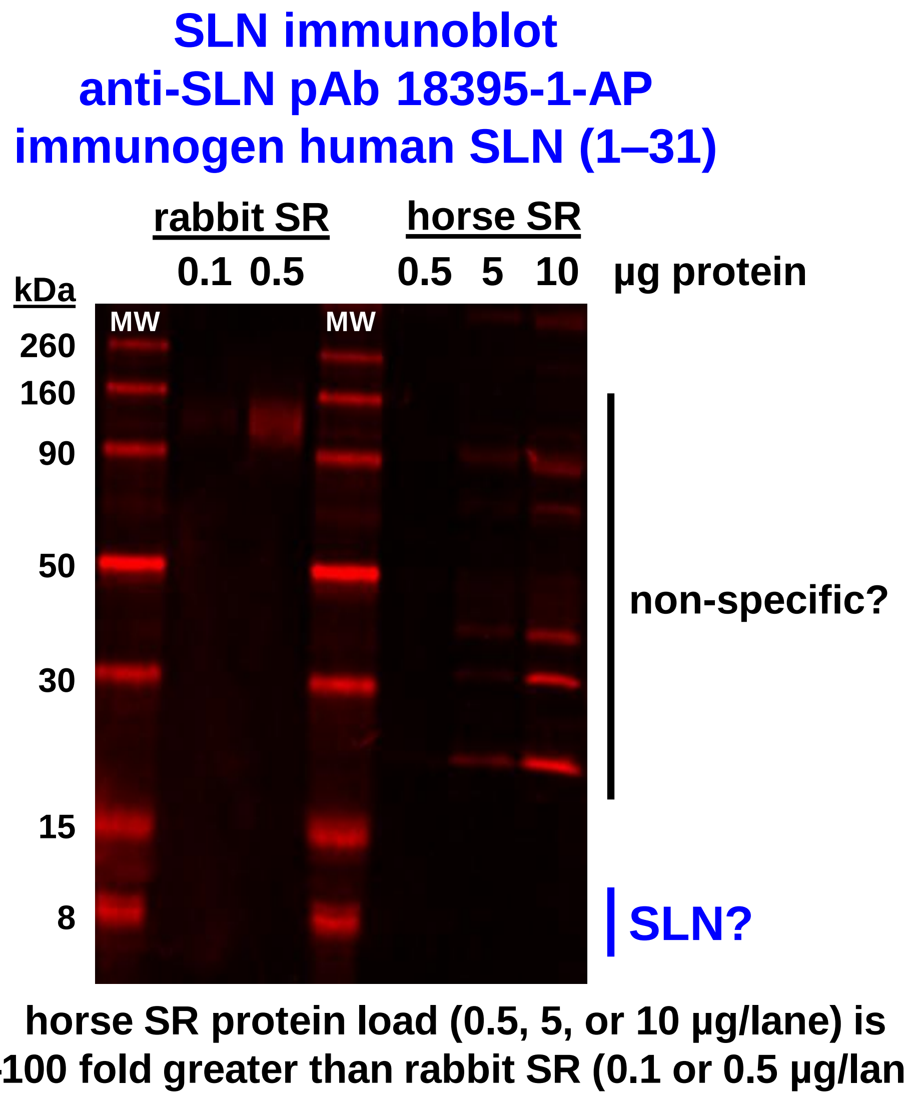


**Figure S9**. **Immunoblot analysis using commercial anti-human-SLN pAb 18395-1-AP does not detect horse or rabbit SLN, but instead shows non-specific binding to other proteins in horse and rabbit SR.** The primary antibody was the commercial anti-SLN pAb18395-1-AP with immunogen comprising human SLN residues 1-31 (see **Fig. 2B**). Samples were electrophoresed through a 4–20% Laemmli gel. Rabbit SR vesicles were loaded at 0.1 and 0.5 µg protein per lane (*left)*, and horse SR vesicles were loaded at 0.5, 5, and 10 µg protein per lane (*right*). Immunoblotting was performed using pAb 18395-1-AP (primary) and goat anti-rabbit-IgG pAb labeled with 680-nm fluorophore (secondary). Immunolabeling was quantified using the LI-COR laser scanner system in the near-infrared fluorescence mode. The molecular mass of protein gel markers (kDa) are indicated on the *left.*

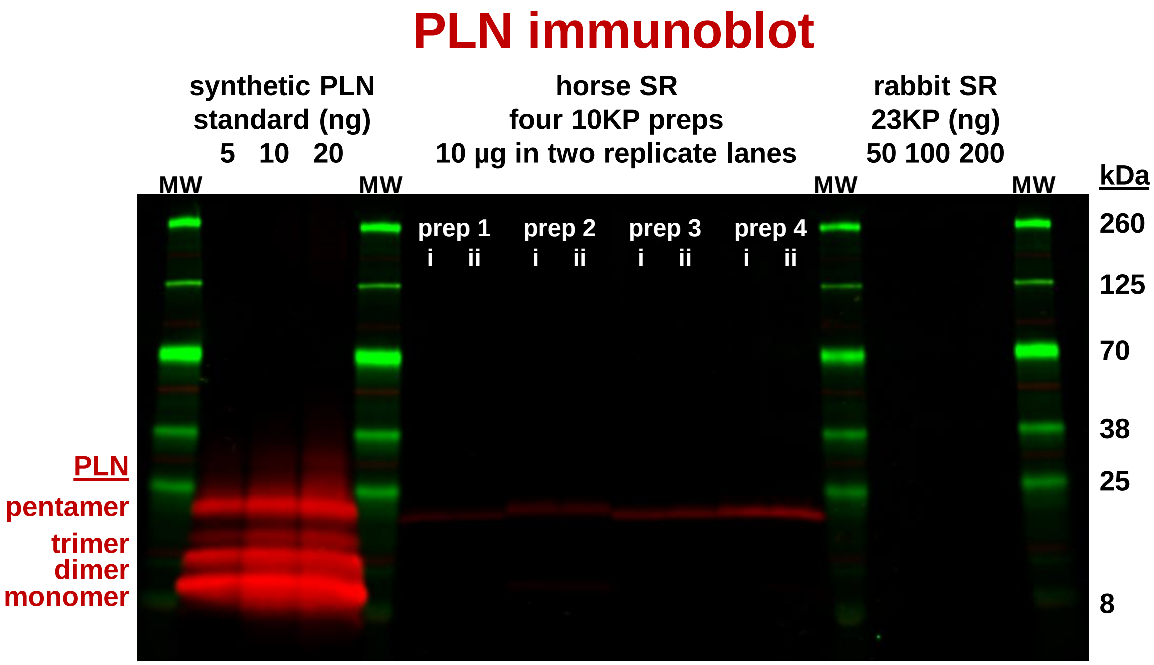


***center*: horse SR protein load (10 µg/lane) is:**
***left*: 500–2000 fold greater than synthetic PLN standard (5, 10, or 20 ng/lane)**
***right*: 50–200 fold greater than rabbit SR protein (50, 100, or 200 ng/lane)**

**Figure S10**. **Immunoblot analysis using anti-universal-PLN mAb 2D12 detects minimal expression of PLN protein in horse and rabbit SR.** The primary antibody was the commercial anti-PLN mAb 2D12 with epitope $^{7}$LTRSAAIR$^{13}$ (17,18) a sequence which is identical in the three PLN orthologs: horse (15), rabbit (GenBank accession code Y00761.1 (19)), and human PLN (GenBank M63603.1 (20)). Samples were electrophoresed through a 4–20% Laemmli gel. Horse SR vesicles (N = 4 preps; 10 µg protein per lane; each prep run in duplicate lanes) and rabbit SR vesicles (50, 100, 200 ng) were assayed. Synthetic human PLN (5, 10, 20 ng) was used as a quantitative standard (21,22). Immunoblotting was performed using mAb 2D12 (primary) and goat anti-mouse-IgG pAb labeled with a 680-nm fluorophore (secondary). Immunolabeling was quantified using the LI-COR laser scanner system in the near-infrared fluorescence mode. The molecular mass of protein gel markers (kDa) are indicated on the *right*. The proposed molecular species of PLN (monomer, dimer, trimer, and pentamer) are indicated on the *left*.

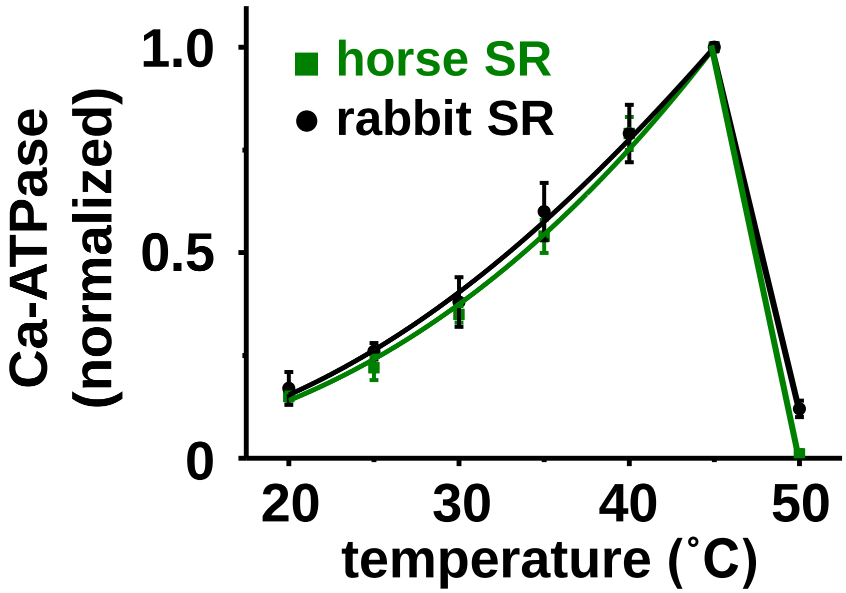


**Figure S11**. **Horse and rabbit SERCA show similar temperature dependence of $Ca^{2+}$-activated ATPase activity.** SR vesicles from horse and rabbit muscle were assayed for ATP hydrolysis in the presence of saturating concentration of substrates (100 µM $Ca^{2+}$ and 5 mM Mg-ATP), in the presence of $Ca^{2+}$ ionophore A23187. The $Ca^{2+}$-ATPase activity of horse and rabbit SERCA shows ~6-fold thermal activation from 20 °C to 45 °C, and then sharp thermal inactivation from 45 °C to 50 °C.

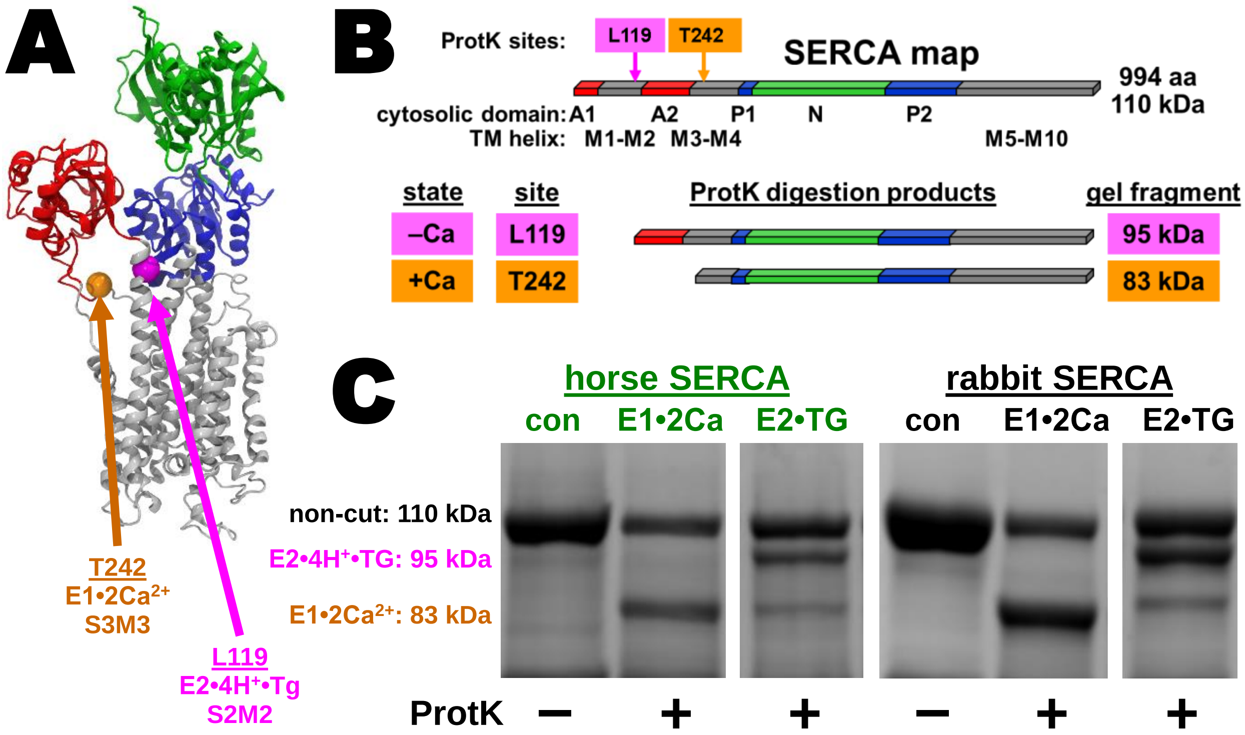


**Figure S12**. **Horse and rabbit SERCA show similar** $Ca^{2+}$**-dependent cleavage by ProtK.** Horse SERCA1a encodes residues L119 and T242, which are identical in the rabbit SERCA1a sequence (i.e., ProtK sites in the absence and presence of $Ca^{2+}$, respectively). *A*, location of conformation-specific ProtK sites in the x-ray crystal structure of rabbit SERCA in the calcium-free E2•TG state (PDB ID code 1IWO (10)). *B*, location of conformation-specific ProtK sites in the primary topology map of rabbit SERCA. *C*, horse and rabbit SR vesicles were digested with ProtK, and proteolytic fragments of SERCA were analyzed by SDS-PAGE and Coomassie staining. ProtK digestions were run in the presence of 100 µ$Ca^{2+}$ to stabilize the $Ca^{2+}$-bound state (E1•2Ca) or in the presence of 1 mM EGTA and 1 µM TG to stabilize the $Ca^{2+}$-free state (E2•TG). The molecular mass of SERCA and diagnostic ProtK fragments are indicated on the *left*. SERCA samples were run on the same Coomassie gel, and the four gel slices shown are presented with the same scale of image intensity.
